# Increasing the resolution and precision of psychiatric GWAS by re-imputing summary statistics using a large, diverse reference panel

**DOI:** 10.1101/496570

**Authors:** Chris Chatzinakos, Donghyung Lee, Na Cai, Vladimir I. Vladimirov, Bradley T. Webb, Brien P. Riley, Jonathan Flint, Kenneth S. Kendler, Kerry J. Ressler, Nikolaos P. Daskalakis, Silviu-Alin Bacanu

## Abstract

Genotype imputation across populations of mixed ancestry is critical for optimal discovery in large-scale genome-wide association studies (GWAS). Methods for direct imputation of GWAS summary statistics were previously shown to be practically as accurate as summary statistics produced after raw genotype imputation, while incurring orders of magnitude lower computational burden. Given that direct imputation needs a precise estimation of linkage-disequilibrium (LD) and that most of the methods using a small reference panel e.g., ~2,500 subject coming from the 1000 Genome Project, there is a great need for much larger and more diverse reference panels. To accurately estimate the LD needed for an exhaustive analysis of any cosmopolitan cohort, we developed DISTMIX2. DISTMIX2: i) uses a much larger and more diverse reference panel and ii) estimates weights of ethnic mixture based solely on Z-scores (when AFs are not available). We applied DISTMIX2 to GWAS summary statistics from the Psychiatric Genetic Consortium (PGC). DISTMIX2 uncovered signals in numerous new regions, with most of these findings coming from the rarer variants. Rarer variants provide much sharper location for the signals compared with common variants, as the LD for rare variants extends over a lower distance than for common ones. For example, while the original PGC post-traumatic stress disorder (PTSD) study found only 3 marginal signals for common variants, we now uncover a very strong signal for a rare variant in *PKN2*, a gene associated with neuronal and hippocampal development. Thus, DISTMIX2 provides a robust and fast (re)imputation approach for most Psychiatric GWAS studies.

## 1. Introduction

Genotype imputation (B. L. Browning & Browning, 2009; B.N. Howie, P. Donnelly, & J. Marchini, 2009; Nicolae, 2006; Servin & Stephens, 2007) methods are commonly used to increase the genomic resolution for large-scale multi-ethnic GWAS meta-analyses (Consortium, 2015; Ripke et al., 2013; Ripke et al., 2011; Sklar et al., 2011; Sladek et al., 2007) by predicting genotypes at unmeasured markers based on cosmopolitan reference panels, e.g. 1000 Genomes (1KG) (Abecasis et al., 2012). However, genotypic imputation is computationally burdensome and requires access to subject-level genetic data, which is harder and slower to obtain than summary statistics.

To overcome these limitations, researchers proposed summary statistics-based imputation methods, e.g. DIST (Lee, Bigdeli, Riley, Fanous, & Bacanu, 2013) and ImpG (Pasaniuc et al., 2013). These methods can directly impute summary statistics (two-tailed Z-scores) for unmeasured Single Nucleotide Polymorphisms (SNPs) from summary statistics of genome-wide association studies (GWASs) or called variants from sequencing studies. The methods were shown to i) substantially reduce the computational burden and ii) be practically as accurate as commonly used genotype imputation methods. These methods were successfully applied in gene-level joint testing of functional variants (Lee et al., 2014) and functional enrichment analyses (Pickrell, 2014). However, these first wave of direct imputation methods were only amenable for imputation in ethnically homogeneous cohorts.

To accommodate cosmopolitan cohorts, DIST method was extended (Lee et al., 2015) to **D**irectly **I**mputing summary **S**tatistics for unmeasured SNPs from **Mix**ed ethnicity cohorts (DISTMIX). It: i) predicted a study’s proportions (weights) of ethnicities from a multi-ethnic reference panel based only on cohort allele frequencies (AFs) for (common) SNPs from the studied cohort or taking prespecified ethnic weights, ii) computed an ethnicity-weighted correlation matrix based on the estimated/prespecified weights and genotypes of ethnicities from the reference panel and then, iii) used the weighted correlation matrix for accurate imputation. Currently, due to privacy concerns (Homer et al., 2008), cohort AFs are lately only rarely provided. To circumvent the lack of AFs, the ARDISS method (Togninalli, Roqueiro, Investigators, & Borgwardt, 2018) extended the inference to cosmopolitan cohorts using a gaussian field approach based only on Z-scores. However, ARDISS software provides only a 1KG-derived reference panel. Unlike DISTMIX that is implemented as a standalone C++ software, ARDISS is implemented in Python that is more temperamental in installing/working with various versions of libraries.

Besides the lack of AFs, another issue that occurred in practical applications of DISTMIX is its reliance on 1KG reference panel which was both small and European-centric, while many meta-analyses have non-trivial fractions of non-European subjects (Cai et al., 2017; Ripke et al., 2013). Since its initial publishing, larger reference cohorts were sequenced and published, e.g. Haplotype Reference Consortium (HRC) (McCarthy et al., 2016) and CONVERGE (Consortium, 2015). Notably, the >11K Han Chinese cohort from the CONVERGE consortium complement nicely HRC panel and provides accurate LD information for the second most studied continental population besides European.

In this paper we propose DISTMIX2 method/software, which addresses the above shortcomings by including two critical components. First, we provide a novel method to accurately estimate ethnic weights of the cohort which uses only summary statistics, e.g. Z-scores. Second, we build a larger, more diverse reference panel with 33K subjects, which combines the subjects from the publicly available part of HRC and CONVERGE. While implementing the above 2 features, DISTMIX2: i) adequately controls the false positive rate, and ii) provides much improved resolution when compared to methods based on older reference panels. Based on simulated data sets we provide guidance on the significance thresholds for rare and/or low information variants. Furthermore, in a practical application to reported summary statistics from studies of psychiatric disorders, we uncover numerous regions harboring signals. Most of these novel signals are associated with rarer variants that could not be robustly interrogating using the smaller panels from previous methods. Some of the new findings provide strong signals in new regions for traits that reported only marginal signals. In the quest to provide enhanced resolution for ethnically mixed GWAS studies, DISTMIX2 provides a robust and substantially faster alternative to the laborious genotype imputation.

## 2. Method

### 2.1 Larger and more diverse reference panel

To facilitate imputation of rarer variants, the current version uses the 33,000 subjects (33K) as reference panel. It consists of 20,281 Europeans, 10,800 East Asians, 522 South Asians, 817 Africans and 533 Native Americans (Text S1, Table S1 in SI). The reference panel includes the publicly available 22,691 subjects from Haplotype Reference Consortium (HRC) and 10,262 CONVERGE. For CONVERGE subjects, we used province of origin to assign them to 4 population (China North East - CNE, China Central East - CCE, China South East - CSE and China Central South - CCS). HRC subjects coming from the small Orkney (ORK) island provided the basis for an extra European population, i.e. ORK. Subjects from 1KG in HRC sample, CONVERGE and ORK along with their a) population label, b) first 20 ancestry principal components were used to train a quadratic discriminant model for predicting population label from principal components. Subsequently, to have more homogeneous populations in the panel, all available subjects were assigned(reassigned) population labels based on model prediction. Consequently, a subject might be re-assigned to a different (but related) population.

Finally, our reference panel contains twenty-six million SNPs. To have reasonably accurate SNP LD estimators, we eliminate the rarest SNPs which did not have at least: i) 20 alleles in European or East Asian superpopulations or ii) 5 in African, South Asian and America native superpopulations.

### 2.2 Converge haplotypes

#### 2.2.1 DNA sequencing

DNA was extracted from saliva samples using the Oragene protocol. A barcoded library was constructed for each sample. Sequencing reads obtained from Illumina Hiseq machines were aligned to Genome Reference Consortium Human Build 37 patch release 5 (GRCh37.p5) with Stampy (v1.0.17) (Lunter & Goodson, 2011) using default parameters, after filtering out reads containing adaptor sequencing or consisting of more than 50% poor quality (base quality <= 5) bases. Samtools (v0.1.18) (Li et al., 2009) was used to index the alignments in BAM format (Li et al.) and Picardtools (v1.62) was used to mark PCR duplicates for downstream filtering. The Genome Analysis Toolkit’s (GATK, version 2.6). Base quality score recalibration (BQSR) was then applied to the mapped sequencing reads using BaseRecalibrator in Genome Analysis Toolkit (GATK, basic version 2.6) (DePristo et al., 2011) with the known insertion and deletion (INDEL) variations in 1000 Genomes Projects Phase 1 (Genomes Project et al., 2010) and known single nucleotide polymorphisms (SNPs) from dbSNP (v137, excluding all sites added after v129) excluded from the empirical error rate calculation. GATKlite (v2.2.15) was then used to output sequencing reads with the recalibrated base quality scores while removing reads without the “proper pair” flag bit set by Stampy (1-5% of reads per sample) using the --read_filter ProperPair option (if the “proper pair” flag bit is set for a pair of reads, it means both reads in the mate-pair are correctly oriented, and their separation is within 5 standard deviations from the mean insert size between mate-pairs).

#### 2.2.2 Variant calling, imputation, and phasing

Variant discovery and genotyping (for both SNPs and INDELs) at all polymorphic SNPs in 1000G Phase1 East Asian (ASN) reference panel(Genomes Project et al., 2012) was performed simultaneously using post-BQSR sequencing reads from all samples using the GATK’s UnifiedGenotyper (version 2.7-2-g6bda569). Variant quality score recalibration (VQSR) was then performed with GATK’s VariantRecalibrator (v2.7-4-g6f46d11) in SNP variant calls using the SNPs in 1000 Genomes Phase 1 ASN Panel (Genomes Project et al., 2010) as the known, truth and training sets. A sensitivity threshold of 90% to SNPs in the 1000G Phase1 ASN panel was applied for SNP selection for imputation after optimizing for Transition to Transversion (TiTv) ratios in SNPs called. Genotype likelihoods (GLs) were calculated at selected sites using a sample-specific binomial mixture model implemented in SNPtools (version 1.0), and imputation was performed at those sites without a reference panel using BEAGLE (version 3.3.2) (S. R. Browning & Browning, 2007). The second round of imputation was performed with BEAGLE on the same GLs, but only at biallelic SNPs polymorphic in the 1000G Phase 1 ASN panel using the 1000G Phase 1 ASN haplotypes as a reference panel. The genotypes derived from Beagle imputation were phased using Shapeit (version 2, revision 790) (Delaneau, Howie, Cox, Zagury, & Marchini, 2013). Genetic maps were obtained from the Impute2 (B. N. Howie, P. Donnelly, & J. Marchini, 2009) website. Chromosomes 13 - 22 and X were phased using 12 threads and default parameters. Chromosomes 1-12 were phased using 12 threads in four chunks that overlap by 1MB. The phased chunks were ligated together using ligateHAPLOTYPES, available from the Shapeit website. A final set of allele dosages and genotype probabilities was generated from these two datasets by replacing the results in the former with those in the latter at all sites imputed in the latter. We then applied a conservative set of inclusion threshold for SNPs for genome-wide association study (GWAS): a) p-value for violation HWE > 10^−6^, b) Info score > 0.9, c) MAF in CONVERGE > 0.5% to arrive at the final set of 6,242,619 SNPs. Details can be found in (Cai et al., 2017).

### 2.3 Automatic detection of cohort composition

Our group has previously described, in DISTMIX paper (Lee et al., 2015), a method to estimate the ethnic composition when the cohort allele frequencies (AF) are available. However, lately some consortia do not provide such measure; they often provide only the AF for Caucasian / European cohorts. *Consequently, there is a great need to estimate the ethnic composition of the cohort even when no AFs are provided*.

Below is the theoretical outline of such method. Suppose that the cohort genotype is a mixture of genotypes belonging *k* ethnic groups from the reference panel. The *G_ij_* denotes the genotype for the *i*-th subject at the *j*-th SNP which belongs to the *l*-th group, let 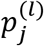 be the frequency of the reference allele frequency for this SNP in the *l*-th group. Let 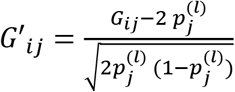 be the normalized genotype, i.e. the transformation to a variable with zero mean and unit variance. Near *H*_0_, SNP Z-score statics *Z_j_*′s have the approximately the same correlation matrix as the genotypes used to construct it, *G*_∗*j*_’s (Lee et al., 2014); given that *G*′_∗*j*_’s are linear combination (with positive slope) of *G*_∗*j*_′s, it follows that Z-scores have the same correlation structure *G*′_∗*j*_′s. However, given that both *G*_∗*j*_′*s* and *Z_j_′s* have unit variance, it follows that the two have the same covariance (i.e. not only the same correlation) structure. Therefore, for any *s* ≥ 1 *E*(*Z_j_ Z_j+s_*) = *E*(*G*′_∗*j*_ *G*′_∗ *j*+s_), which, due independence of genotypes in different ethnic groups becomes:

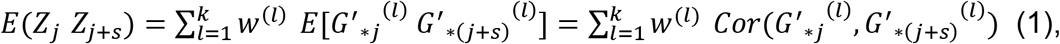

where *w*^(*l*)^ is the expected fraction of subjects from the entire cohort that belong to the *l*-th group.

While *Cor*(*G*′_∗*j*_^(*l*)^, *G*′_∗(*j*+s)_^(*l*)^) is unknown, it can be easily approximated using their reference panel counterparts. Thus, the weights, *w*^(*l*)^, can be estimated by simply regressing the product of reasonably close SNP *Z*-scores, *Z′_j_ Z′_j+s_*, on correlations between normalized genotypes at the same SNP pairs for all subpopulations in the reference panel. To increase bias power, we chose the parameter *s*, such as to maximize the variance of the within-panel ethnic group correlations while keeping *j* + *s*-th SNP no more than 50Kb away from *j*–th SNP. Because some GWAS might have numerous large signals, e.g. latest height meta-analysis (Ripke et al., 2013; Wood et al., 2014), a more accurate estimation of the weights is very likely to be obtained by substituting expected gaussian quantiles for 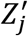 (see **Nonparametric robust estimation of weights subsection**).

Due to the strong LD among SNPs, the estimation of the correlation using all SNPs in a genome might lead to a poor regression estimate in (1). To avoid this, we sequentially split GWAS SNPs into 1000 non-overlapping SNP sets, e.g. first set consists of the 1st, 1001st, 2001st, etc. map ordered SNPs in the study. The large distances between SNPs in the same set make them quasi-independent which, thus, improves the accuracy of the estimated correlation. *W* = (*w*^(*l*)^) is subsequently estimated as the average of the weights obtained from the 1000 SNP sets. Finally, we set to zero the negatives weights and normalize the remaining weights to sum to 1 (Chatzinakos et al., 2017). This method should be even more useful when we already know the approximate continental (EUR, ASN, SAS, AFR and AMR) weights (as estimated from study information) but it is not always clear how these proportions should be allocated among continental subpopulations. This further apportioning of continental weights is likely to be extremely important when the GWAS cohorts contain many admixed populations, e.g. African Americans and American native populations. Consequently, when continental proportions are provided by the users, we use our automatic detection to distribute these weights to the most likely subpopulations in the reference panel. To eliminate unforeseen artifacts, we strongly recommend to the users to provide continental proportions when AFs are not available.

### 2.4 Nonparametric robust estimation of weights

To estimate robust weights and to avoid false positives, we apply a two-step, robust algorithm to the *Z*-scores of the SNPs. **First,** let *Z_σ_* = (*z*_*σ*_1__, *z*_*σ*_2__,…, *z_σ_m__*), where *σ* indicates the permutation of indices for sorting in increasing order Z-scores, *Z*, for the *m* SNPs. **Second,** 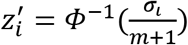, where *Φ*^−1^ is the inverse normal cumulative distribution function. Subsequently, these transformed risk scores are used for estimating ethnic weights.

## 3. Simulation

To estimate the accuracy and false positive rates of DISTMIX2, for five different cosmopolitan studies scenarios, we simulated (under *H*_0_: no association between trait and variants) 100 cosmopolitan cohorts of 10,000 subjects for autosomal SNPs in Ilumina 1M panel (Lee et al., 2015) using 1KG haplotype patterns (Text S1, Table S2 in SI). The subject phenotypes were simulated independent of genotypes as a random Gaussian sample. SNP phenotype-genotype association summary statistics were computed from a correlation test.

The accuracy of the procedure was assessed by masking 5% of the SNPs (Experiment 1, Table 1). Subsequently, the true values and the imputed values at these masked SNPs were used to compute: i) their correlation and ii) the mean squared error of the imputation. We assess these measures at four different levels of MAF. To compare the Type I error rate of our proposed method, DISTMIX2, we estimated the relative Type I error (the empirical divided by the nominal Type I error rate) as a function of the nominal Type I error rate, for the same four MAF levels for all the cohorts. Finally, for all the combinations between MAFs and Info we performed DISTMIX2 analyses with three different parameters for the length of the predicted window (the length of the predicted window also depends on the minimum number of measured SNPs it encompasses).

**Table 1.**
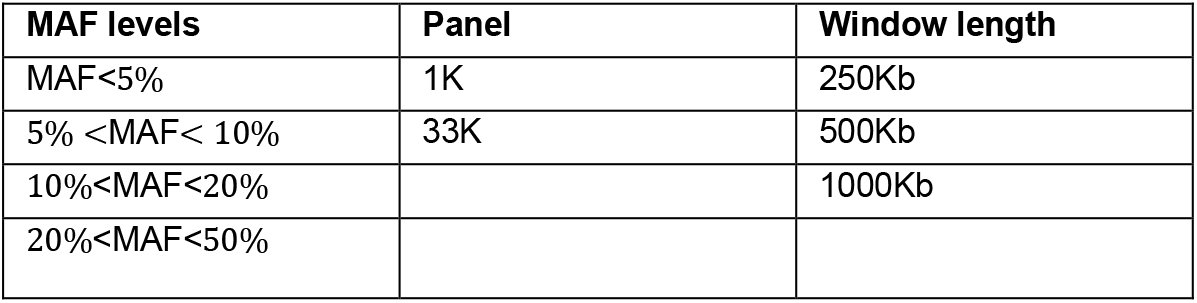
Experiment 1 parameter settings.

To assess the reliability of DISTMIX2 results for rare and very rare variants, for the above cohorts, we also estimate DISTMIX2 size of the test for very low MAFs (rare variants), (Experiment 2, Table 2). The size of the test is assessing for 5 imputation Info intervals and 6 MAF intervals.

**Table 2.** Experiment 2 variable parameter settings. Fixed parameters for this experiment: 33K panel and 500Kb length of predicting window.

| MAF levels | Info levels |
| --- | --- |
| 0.05% <MAF< 0.5% | Info< 20% |
| 0.5% <MAF< 1% | 20%<Info<40% |
| 1% <MAF<2% | 40%<Info<60% |
| 2% <MAF< 5% | 60%<Info<80% |
| 5% <MAF< 10% | Info>80% |
| 10% <MAF< 50% |  |

However, given that: 1) the simulated cohorts might not reflect real data and 2) these data sets do not have the sample sizes needed to detect very rare SNPs (e.g. MAFs < 0.05%), which is important for DISTMIX2 inference in practical applications, we used real data sets to create so-called nullified data sets (Experiment 3, Table 3). These nullified data are based on 20-real and mostly Caucasian GWAS SCZ, ADHD, AUT, MDD and sixteen GWAS meta-analyses that are not yet publicly available. This approximation for null data is obtained by substituting the expected quantile of the Gaussian distribution for the (ordered) Z-score. We note that, while the quantile estimation adjusts the noncentrality parameter (enrichment) of the statistics to zero, it does not change the order of the statistics. One effect of this fact is that imputing statistics within/near the peak signals in original GWASs might result in increased false positive rates and, thus, the genome-wide false positive rates might appear to be moderately inflated.

**Table 3.** Experiment 3 variable parameter settings. Fixed parameters for this experiment: 33K panel and 500Kb window length.

| MAF levels | Info levels |
| --- | --- |
| MAF < 0.05% | Info < 20% |
| 0.05% <MAF < 0.5% | 20% <Info < 40% |
| 0.5% <MAF < 1% | 40% <Info < 60% |
| 1% <MAF < 2% | 60% <Info < 80% |
| 2% <MAF < 5% | Info > 80% |
| 5% <MAF < 10% |  |
| 10% <MAF < 50% |  |

## 4. Practical Application

We applied DISTMIX2 to a subset of the psychiatric summary datasets available for download from Psychiatric Genetics Consortium (PGC-http://www.med.unc.edu/pgc/), i.e. schizophrenia (SCZ), attention deficit hyperactive disorder (ADHD), autism (AUT), eating disorder (ED), bipolar (BIP) disorder, major depressive disorder (MDD) and post-traumatic stress disorder (PTSD) (see Table 4 for references). Based on the results from simulations under the null hypothesis (Experiment 1), for all these practical applications we used: a) the larger 33K size panel and b) a length of the predicted window (500Kb). To improve the imputation of the unmeasured SNPs for SCZ, we denote as “measured SNPs” only those with very high information (Info>0.997). For the ADHD, AUT, BIP, MDD and PTSD data sets, because the imputation information is not available, we accept as measured SNPs the set consisting of the intersection between SNPs in each GWAS and the above SCZ’s “measured” SNPs. Where available (e.g. MDD), we also filtered out SNPs with effective sample sizes below the maximum.

**Table 4.**
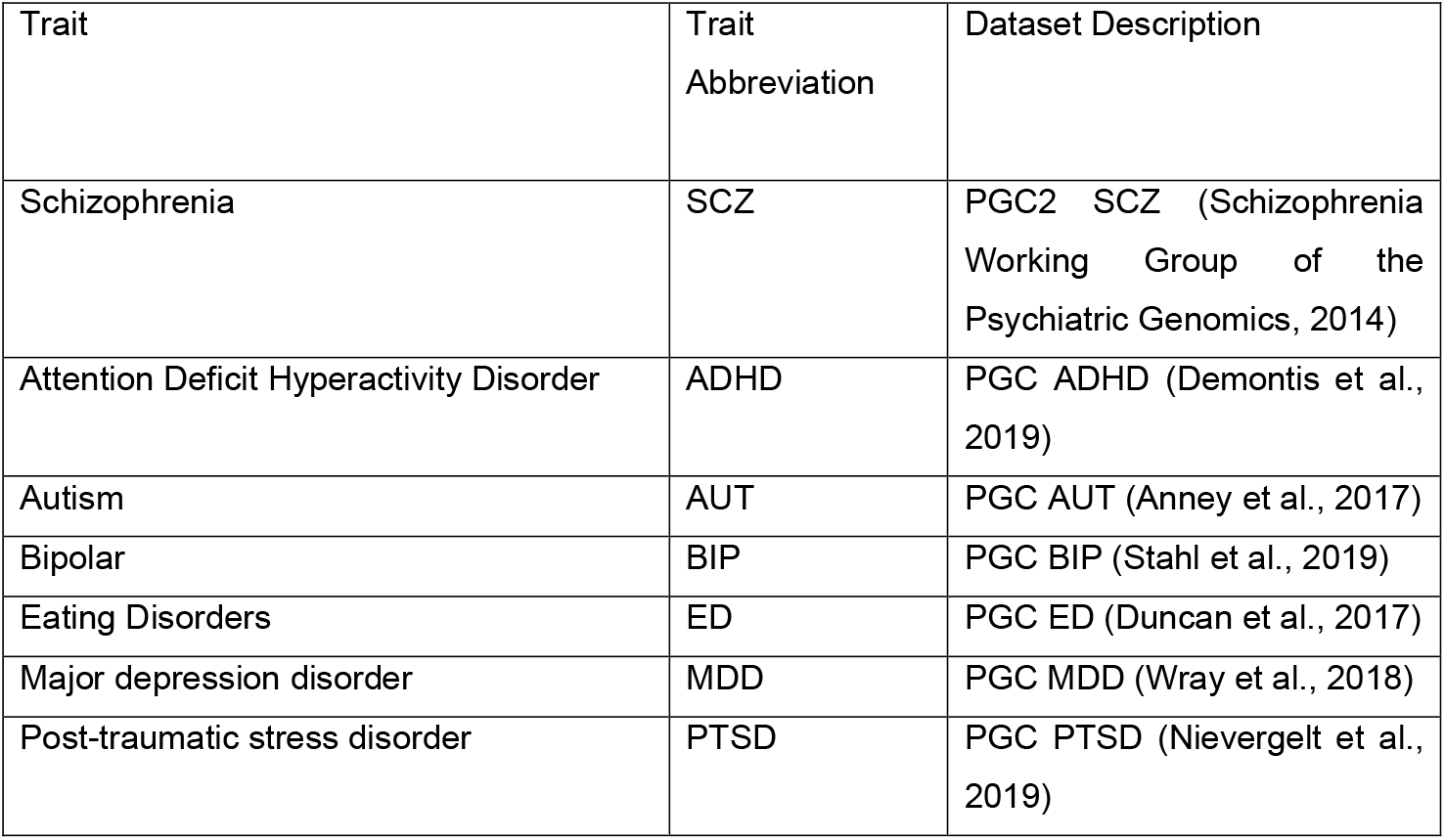
Real dataset description.

## 5. Results

For Illumina 1M SNPs (Marenne et al., 2011) that were masked, and then imputed (see Method evaluation section), DISTMIX2 with our novel automatic ethnic weight detection (see Method section), controls the false positive rates at or below nominal threshold, even at very low type I error, e.g. 10^−6^ (Text S2, Fig S1 in SI). *R*^2^ between true values and estimated ones is above 0.92 for our five simulated mixed-cohort scenarios (Text S2, Figs S2-S6 in SI). Also, DISTMIX2 imputed statistics have very good mean squared error (RMS) (Text S2, Figs S7-S11 in SI). For the above three measurements (size of the test, *R*^2^ and RMS) the setting of 250Kb for the length of the predicted window was the least precise, while 500Kb and 1000Kb had practically identical precision.

For rare and very rare variants, the size of the test is up to 300-1,000X higher than the nominal one and even up to 5,000-10,000X for cohorts that have large fractions of subpopulations that are underrepresented in the reference panel (e.g. Americans, Africans etc.), especially for the setting Minor Allele Frequency (MAF), 0.05%<MAF<0.5% and Information (Info), Info<0.2 (Text S2, Figures S12-S47 in SI).

For the “nullified” data sets, e.g. those obtained from real data sets by substituting the study Z-scores by their expected quantile under the null hypothesis (*H*_0_) (Method evaluation section and Text S2, Figs S42-S48 in SI), DISTMIX2 controls reasonably well the size of the test - up to 20X higher than the nominal rate (even for SNPs with low MAFs and low Info).The minimum GWAS p-values for the nullified data sets that were imputed ranged between 8.13 ∗ 10^−7^ and 1.11 ∗ 10^−11^. By fitting a normal distribution to −*log*_10_ (minimum p-values), we estimated the mean to be 8.655 and the standard deviation to be 1.172. Using as criterion the conservative three standard deviations above the mean, we obtain from these realistic data a 12.17 as the upper bound for the −*log*_10_ (p-value). I.e. in DISTMIX2 practical applications (PGC traits), a conservative threshold for significance is 10^−12^, regardless of imputation Info and SNP MAF. *Consequently, in all applied analyses in this paper we add this very stringent threshold for DISTMIX2 imputed summary statistics*. Using as criterion the even more conservative five standard deviations above the mean (the very conservative Chebyshev inequality for the upper bound of the p-value of exceeding this threshold= 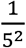 = 0.04), we obtain a 14.515 upper bound for the −*log*_10_ (p-values), i.e. a super-conservative significance threshold of 3 ∗ 10^−15^.

For the practical applications to PGC traits (Table 4), we construct Manhattan plots for all autosome chromosomes (1-22) and, individually, for chromosomes harboring novel signals (defined as imputed SNPs with statistically significant p-values that are at least 250Kb away from the reported GWAS signal) (Fig. 1-2, Text S3, Fig. S49-S59 in SI). Furthermore, in order to investigate the potential risk of genomic inflation, we construct Q-Q plots for the following three scenarios (i) all the SNPs, (ii) rare SNPs and (iii) common SNPS, for all the traits (Fig. 3, Text S3, Fig. S60-S78 in SI). Finally, we compared DISTMIX2 with ARDISS (Text S3, Fig. S79-S85 in SI). Since ARDISS software does not provide minor allele frequency and Info estimation we subset the imputed signals according to DISTMIX2 for *MAF* > 0.05. For all Manhattan plots we draw two dash lines denoting threshold for statistically significant signals. The red line is the default genome-wide threshold of *p* = 5 ∗ 10^−8^, which is applicable to signals from measured SNPs and common imputed SNPs with high Info values. The purple line at *p* = 10^−12^ is the threshold to be used for rare/very rare variants and/or variants with low information; it corresponds to the above mentioned upper bound for nullified data. As an illustration, we present PTSD Manhattan plot for all chromosomes, only for chromosome 1 and Q-Q plots for all signals (Fig.1, Fig.2-Fig.3 respectively).

**Fig. 1.**
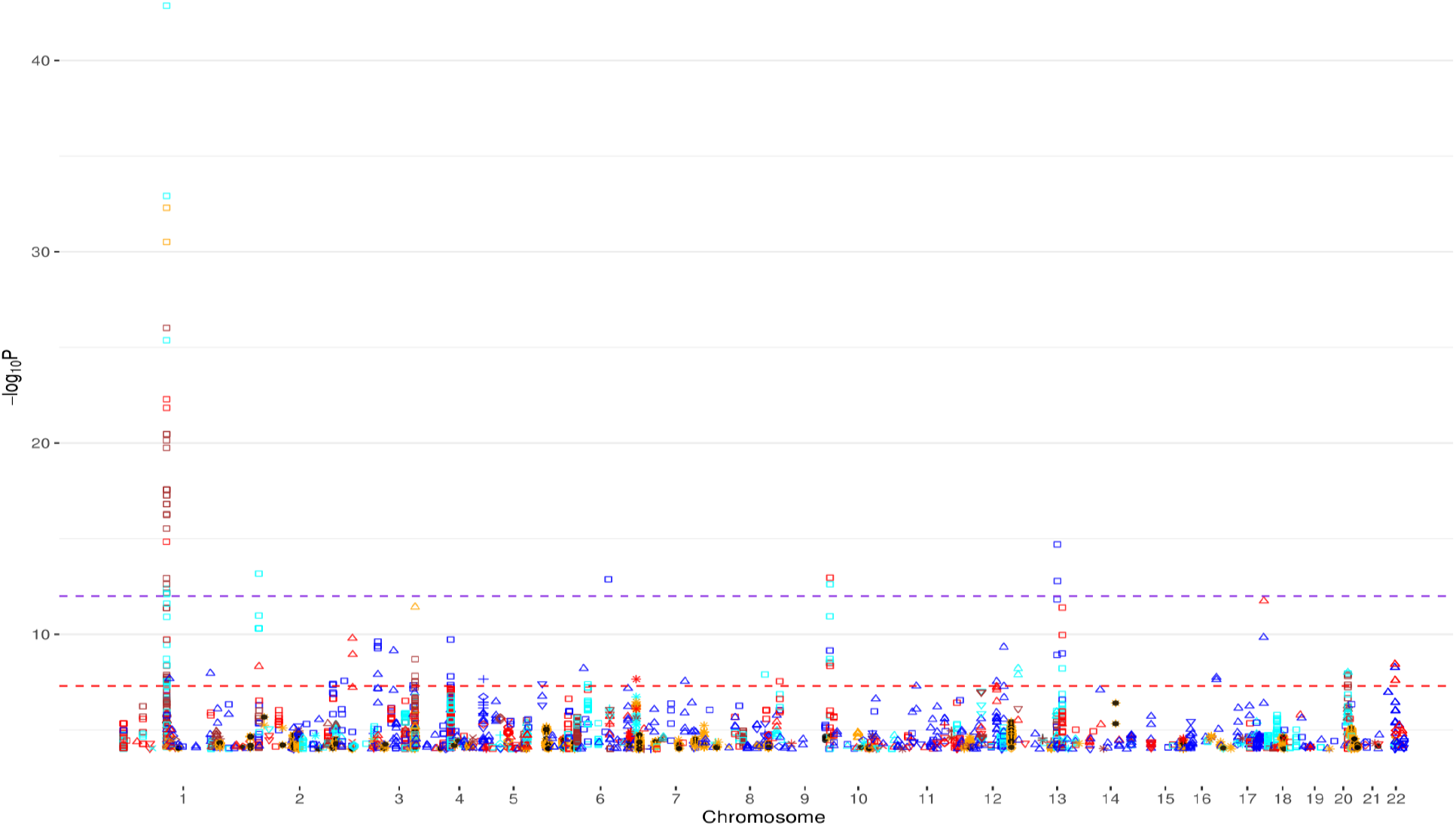
Manhattan plot for chromosomes 1-22 for PTSD. ● denotes reported signals from the original GWAS and the remain symbols and colors denote DISTMIX2 imputed signals. Among imputed signals blue denotes info<0.2, red denotes 0.2<info<0.4, cyan denotes 0.4<info<0.6, brown denotes 0.6<info<0.8, orange denotes info>0.8, □ denotes MAF <0.05%, △ denotes 0.05%<MAF<0.5%, ▽ denotes 0.5%<MAF<1%, + denotes 1%<MAF<2%, ♢ denotes 2%<MAF<5%, *x* denotes 5%<MAF<10% and ❊ denotes 10%<MAF<50%. The red line is the default genome-wide threshold of *p* = 5 ∗ 10^−8^, which is applicable common SNPs with moderate to large Info values. The purple line at *p* = 10^−12^ is the threshold to be used for rare and/or low Info variants.

**Fig 2.**
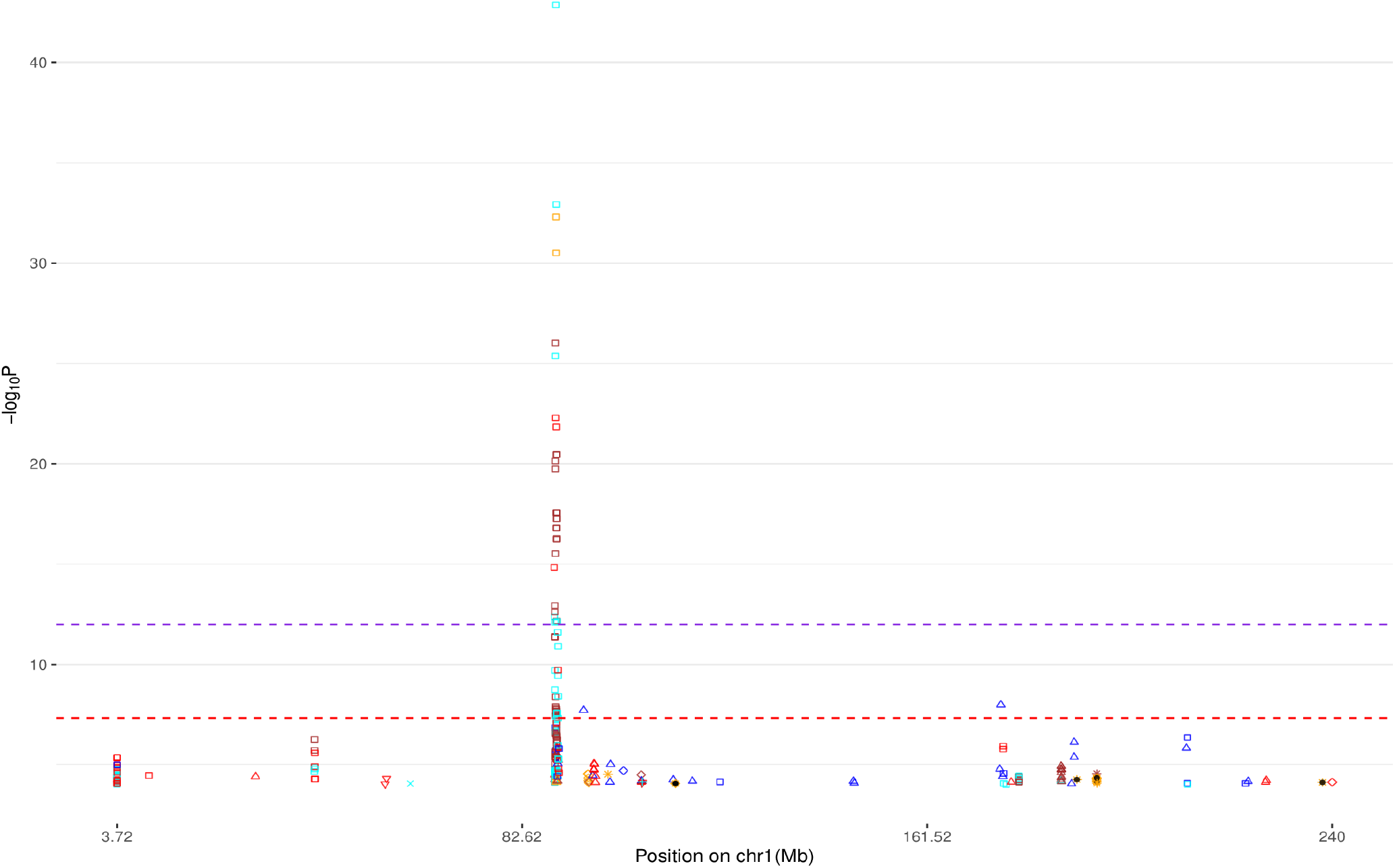
Manhattan plot for chromosome 1 for PTSD (see Fig 1. for background).

**Fig 3.**
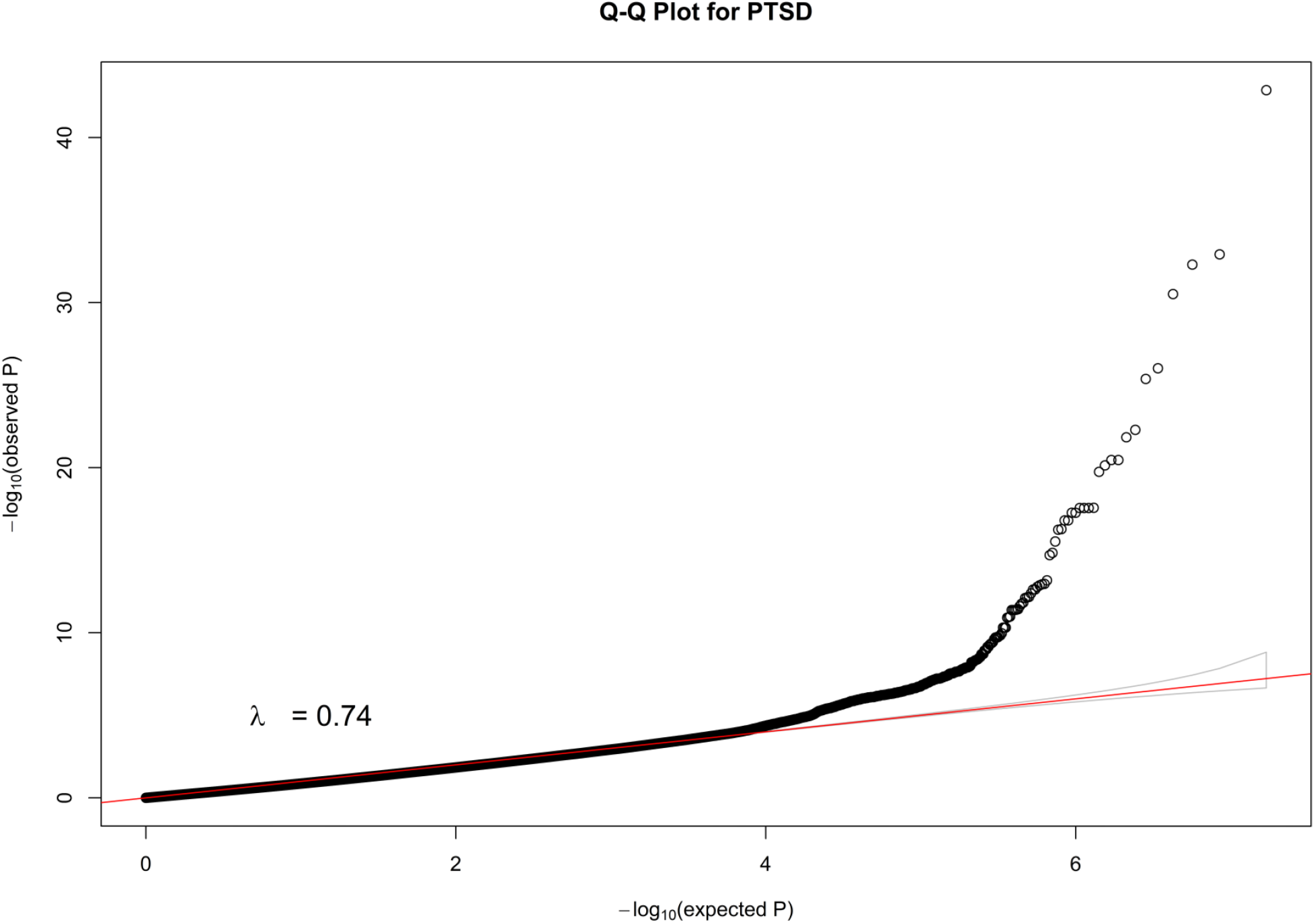
Q-Q plot for all SNPs of PTSD.

These applications of DISTMIX2 to PGC data sets suggests the existence of numerous new signals, most associated with rare SNPs (see Table 5). For instance, in chromosome 12 for schizophrenia (rs143374), with MAF=0.0007, Info=0.245 and p-value=9.26 ∗ 10^−46^ the magnitude of the p-value along with the lambda of the correspond Q-Q plot of the PTSD trait suggest that this signal is likely not to be an artifact (above the most stringent threshold), in chromosome 11 for ADHD (rs5681132) where the MAF=0.0004, the Info= 0.018 and p-value= 7.40 ∗ 10^−16^, in chromosome 22 for AUT (rs1380986), with MAF= 0.0006, Info= 0.498 and p-value= 8.01 ∗ 10^−15^, in chromosome 7 for BIP (rs76350051), with MAF=0.0004, Info=0.04 and p-value=2.47 ∗ 10^−37^, in chromosome 12 for MDD (rs567868887), with MAF=0.0009, Info=0.28 and p-value=1.57 ∗ 10^−55^, and in chromosome 1 for PTSD (rs150642422), with *MAF* = 0.0002, *Info* = 0.5512 and p-value=1.3 ∗ 10^−43^.

**Table 5.** Best three signals for each PGC dataset. Bolded red entries correspond to the most stringent threshold of *p* < 3 ∗ 10^−15^, not bolded red to the second most conservative threshold 3 ∗ 10^−15^ < *p* < 10^−12^ and not bolded blue 10^−12^ < *p* < 5 ∗ 10^−8^.

| Trait | rs_id | chr | bp | val | Info | MAF | Genes/Distance (Kbp) |
| --- | --- | --- | --- | --- | --- | --- | --- |
| ADHD | **rs568113293** | **11** | **54,899,533** | **$7.40 * 10^{-16}$** | **0.0189** | **0.00049** | **TRIM48,130.125**<br>**RP11-72M10.2,135.351** |
| | rs544637819 | 3 | 15,310,737 | $1.78 * 10^{-14}$ | 0.1543 | 0.00171 | SH3BP5, 0<br>SH3BP5-AS1, 4.737 |
| | chr6:30450452 | 6 | 30,450,452 | $6.44 * 10^{-13}$ | 0.0698 | 0.00151 | RANP1, 3.265<br>HLA-E, 6.792 |
| AUT | rs138098629 | 22 | 36,584,165 | $8.01 * 10^{-15}$ | 0.4980 | 0.00063 | APOL4, 1.007<br>MTND1P10, 8.732 |
| BIP | **rs76350051** | **7** | **64,164,245** | **$2.42 * 10^{-37}$** | **0.0417** | **0.00046** | **ZNF107, 0**<br>**RP11-561N12.7, 16.474** |
|  | **rs138549126** | **3** | **52,592,843** | **$6.65 * 10^{-16}$** | **0.073** | **0.00052** | **SMIM4, 0**<br>**PBRM1, 0** |
|  | **rs149257260** | **15** | **71,600,045** | **$1.40 * 10^{-15}$** | **0.4246** | **0.00017** | **THSD4, 0**<br>**RP11-592N21.1, 33.421** |
| ED | rs78958069 | 8 | 43,539,021 | $4.17 * 10^{-10}$ | 0.005 | 0.0002 | RP11-643N23.2, 10.803<br>AC134698.1, 123.105 |
| | rs144485994 | 20 | 4,963,320 | $5.18 * 10^{-9}$ | 0.15 | 0.0001 | SLC23A2, 0<br>RP5-1116H23.1, 30.846 |
| MDD | **rs567868887** | **12** | **31,931,432** | **$1.57 * 10^{-55}$** | **0.2800** | **0.00098** | **H3F3C, 12.691**<br>**RP11-467L13.5, 23.377** |
|  | **rs112241719** | **11** | **111,514,969** | **$8.14 * 10^{-45}$** | **0.4900** | **0.00025** | **SIK2, 0**<br>**AP000925.2, 26.711** |
|  | **rs182264017** | **1** | **188,992,506** | **$5.05 * 10^{-44}$** | **0.2775** | **0.00035** | **LINC01035, 0**<br>**CLPTM1LP1, 12.586** |
| PTSD | **rs150642422** | **1** | **89,223,553** | **$1.3 * 10^{-43}$** | **0.5512** | **0.0002** | **PKN2, 0**<br>**RNU6-125P, 58.909** |
|  | **rs7521099** | **1** | **88,875,371** | **$1.4 * 10^{-15}$** | **0.4521** | **0.0002** | **GBP3, 248.796** |
| | rs111229512 | 2 | 31,800,662 | $6.7 * 10^{-14}$ | 0.6325 | 0.0002 | BP7, 373.881<br>GBP4, 423.278 |
| SCZ | **rs559199817** | **3** | **17,267,731** | **$1.30 * 10^{-87}$** | **0.0213** | **0.00073** | **TBC1D5, 0**<br>**AC090644.1, 90.517** |
|  | **rs143337489** | **12** | **11,2089,686** | **$9.26 * 10^{-46}$** | **0.2464** | **0.00019** | **BRAP, 0**<br>**PCNPP1, 17.97** |
|  | **rs193224736** | **16** | **8,593,132** | **$3.79 * 10^{-21}$** | **0.28476** | **0.00018** | **RP11-483K5.3, 11.117**<br>**TMEM114, 26.37** |

When imputing in parallel SNPs regions of 40 Mbp, the analysis of each data set had a running time of less than 5 days on a cluster node with 4x Intel Xeon 6 core 2.67 GHz.

## 6. Discussion

DISTMIX2, is a software/method for “off-the-shelf” direct imputation of the unmeasured SNP statistics in cosmopolitan cohorts. The main features of the updated version are: 1) a much larger (33K subjects) and more diverse (includes ~11K Han Chinese) reference panel and 2) a procedure for estimating the ethnic composition of the cohort without the need for AF information. Using simulated and the very novel nullified (real) data sets we propose conservative and very conservative significance thresholds for low info and low MAF signal. Application of DISTMIX2 to PGC data sets provides numerous new signal regions, most harboring rarer variants.

It is noteworthy that we uncovered a potentially very strong signal in PGC PTSD (p<10^−42^) in a rare variant of *PKN2* gene, when the initial publication reported only 3 marginal signals on common variants. While *PKN2* has not been extensively characterized, it would be a potentially interesting target in PTSD given that it has been associated with Rho/Rock and mTOR cell pathways previously associated with fear learning and processing (Lachmann et al., 2011; Schmidt, Durgan, Magalhaes, & Hall, 2007; Wallace, Magalhaes, & Hall, 2011). It has also been associated with hippocampal functioning and development (Buchser, Slepak, Gutierrez-Arenas, Bixby, & Lemmon, 2010; Schmidt et al., 2007). All these cellular and neurobiological processes have been established as important in PTSD development and recovery (Maddox, Hartmann, Ross, & Ressler, 2019; Parsons & Ressler, 2013).

Due to our reassignment of subjects to subpopulations when constructing the 33K reference panel, the naive assignment of the pre-estimated weights to only specific subpopulations from the reference panel that are considered the closest ones to the perceived cohort composition, can greatly increase the type I error (false positives). For that reason, when AF is not available, we recommend that users provide continental cohort weights (i.e. European [EUR], East Asian [ASN], South Asian [SAS], African [AFR] and America native [AMR]) and our software automatically will allocate these meta-weights to the most likely within-continent subpopulations. However, when AF is available there is no need to provide this additional information.

DISTMIX2 maintains the type I error reasonably accurately, even for low MAFs and low Info variants, especially for mostly European and East Asian cohorts that are overrepresented in our reference panel. When MAF>5% (common variants), DISTMIX2 appears to maintain the false positive rates up to an order of magnitude higher than the nominal ones for all levels of information. Simulation results suggest that, when a larger part of study cohort consist of subpopulations underrepresented in our reference panel, it is reasonable to lower (by a factor of ~10,000) the genome-wide Bonferroni threshold of significance for p-values of imputed rarer variants. For imputed variants (especially rarer or with lower Info) in study of Europeans, we also use novel nullified data sets to propose a conservative threshold for significance of *p* = 10^−12^ and a very conservative threshold of *p* = 3 ∗ 10^−15^. This guidance is likely to be useful for similar methods when users enlarge their reference panels.

The length of the prediction window (250Kb, 500Kb, 1000Kb) is an important design parameter due to its implications for speed and precision of analyses. Simulations results suggest that, while the accuracies for 500Kb and 1000Kb estimates are very close, the computational burden increases ~2.5 times for the 1000kb window. For that reason, we recommend that researchers use a 500Kb prediction window.

While mentioned only briefly in this manuscript, for practical application we use as “measured” SNP in the input summary statistic file only the GWAS SNPs reported to have close to perfect information and/or effective sample size. Our approach is rooted in preserving the cardinal assumption, of our and all but one other imputation methods (Rueger, McDaid, & Kutalik, 2018), that the LD between SNP Z-scores is very well approximated by the LD of the same SNPs in the reference panels. It is well known that when there are non-negligible missing rates for the variant pair this assumption is not met (Rueger et al., 2018). While the LD of Z-scores can be estimated by making reasonably realistic assumptions about co-missingness patterns of such SNP pairs, to avoid even the rarer circumstances in which these assumptions might not be met, we decided to avoid such an approach. Consequently, we employed (and recommend) the conservative approach of deeming as measured only SNPs with close to perfect imputation information and/or effective sample sizes in the original GWAS.

In practical applications, the very low MAF and Info for some SNPs are associated with up to 4 orders of magnitude inflation in false positive rates, especially when the cohorts contain many subjects belonging to populations that are underrepresented in the reference panel. While signals for rarer SNPs from PGC data sets reported in this paper can be viewed as much “softer” signals than the ones associated with common and high Info variants, the very low p-values for some of them (e.g. p<10^−42^ in *PKN2* gene for PTSD) suggest that most of these signals are likely to be real. This suggestion is enhanced by the fact that, to avoid the pitfalls of estimating covariances from just very few minor alleles, we did not include in the imputation panel SNPs that do not have at least: i) 20 minor alleles in the Europeans or East Asians or ii) 5 minor alleles in all other continental groups. Nonetheless, we recognize that signals for these SNPs should be treated with more skepticism than the more common/higher Info variants and subjected to more stringent wet-lab validations.

## Software and data availability

DISTMIX2 executable and reference panel are freely available for academic use at https://github.com/Chatzinakos/DISTMIX2. The DISTMIX2 executable requires the user to input only the GWAS summary statistics.

## Notes

### Competing Interest Statement

The authors have declared no competing interest.

### Summary of Updates

Direct imputation of summary statistics was shown to be orders of magnitude less computationally intensive than the genotype imputation route. However, it requires a precise estimation of linkage disequilibrium (LD) for mixed ethnicity (cosmopolitan) cohorts. We propose a method (DISTMIX2) that: 1) automatically estimates the ethnic composition of the cohort and uses it for accurately estimating LD for SNPs, 2) uses a novel, large and diverse reference panel consisting of 33,000 subjects, including 11,000 Han Chinese and 3). DISTMIX2 can be a robust and fast (re)imputation approach for most Psychiatric GWAS studies. We believe that the proposed method is the first method that automatically detects cohort composition and uses this information, along with the diverse panel, to provide a robust and fast discovery tool for uncovering signals (even for rarer variants) in numerous new regions.

