## Supplemental Material for "Increasing the resolution and precision of psychiatric GWAS by re-imputing summary statistics using a large, diverse reference panel"

**S1. Reference panel description.**

**Table S1. Subpopulations in the reference panel and their continental cohort (super population).** EUR is the abbreviation for Europeans, AFR for Africans, ASN for Asians, AMR for Americans and SAS for south Asians.

| Population Abbreviation | Number of Subjects | Super Population | Population Description |
|---|---|---|---|
| ACB | 164 | AFR | African Caribbeans in Barbados |
| ASW | 162 | AFR | African Ancestry in Southwest US |
| BEB | 86 | SAS | Bengali from Bangladesh |
| CCE | 3,409 | ASN | China Central East |
| CCS | 2,613 | ASN | China Central South |
| CDX | 95 | ASN | Chinese Dai in Xishuangbanna, China |
| CEU | 6,360 | EUR | Utah residents with Northern and Western European ancestry |
| CLM | 98 | AMR | Colombians from Medellin, Colombia |
| CNE | 2,330 | ASN | China North East |
| CSE | 2,020 | ASN | China South-East |
| ESN | 140 | AFR | Esan in Nigeria |
| FIN | 3,529 | EUR | Finnish in Finland |
| GBR | 2,020 | EUR | British in England and Scotland |
| GIH | 110 | SAS | Gujarati Indian from Houston, Texas |
| GWD | 113 | AFR | Gambian in Western Divisions in the Gambia |
| IBS | 1,309 | EUR | Iberian Population in Spain |

| | | | |
|---|---|---|---|
| ITU | 95 | SAS | Indian Telugu from the UK |
| JPT | 107 | ASN | Japanese in Tokyo, Japan |
| KHV | 226 | ASN | Kinh in Ho Chi Minh City, Vietnam |
| LWK | 99 | AFR | Luhya in Webuye, Kenya |
| MSL | 87 | AFR | Mende in Sierra Leone |
| MXL | 187 | AMR | Mexican Ancestry from Los Angeles, USA |
| ORK | 5,772 | EUR | Orkney Island study |
| PEL | 110 | AMR | Peruvians from Lima, Peru |
| PJL | 121 | SAS | Punjabi from Lahore, Pakistan |
| PUR | 138 | AMR | Puerto Rican in Puerto Rico |
| STU | 110 | SAS | Sri Lankan Tamil from the UK |
| TSI | 1,291 | EUR | Toscani in Italia |
| YRI | 52 | AFR | Yoruba in Ibadan, Nigeria |

**S2. Assessing the Type I error rate**

In order to test DISTMIX2, we considered five different cosmopolitan cohort scenarios based on the 1000 Genomes haplotypic data: 1) $30\%$ CEU + $25\%$ CHS + $5\%$ PUR + $40\%$ YRI (Cohort 1), 2) $10\%$ ASW + $15\%$ CEU + $15\%$ CHB + $12.5\%$ CHS + $15\%$ GBR + $10\%$ MXL + $2.5\%$ PUR + $20\%$ YRI (Cohort 2), 3) $15\%$ ASW + $35\%$ CHB + $35\%$ GBR + $15\%$ MXL (Cohort 3), 4) $45\%$ ASW + $55\%$ GBR (Cohort 4) and 5) $55\%$ CHB + $45\%$ MXL (Cohort 5) (Table S2).

The accuracy of the procedure was assessed by masking $5\%$ of the SNPs (Experiment 1). To compare the Type I error rate of our proposed method, DISTMIX2, we estimated the relative Type I error (the empirical divided by the nominal Type I error rate) as a function of the nominal Type I error rate, for 4 different MAF scenarios (shown on $-log_{10}$ scale in Fig. S1a-S1d). Subsequently, the true values and the imputed ones at these masked SNPs were used to compute i) their correlation, and ii) mean squared error of the imputation (Fig. S2-S11).

To test the size of the test for DISTMIX2 for very low MAFs (rare variants), (Experiment 2), i) we imputed all the SNPs and again we estimated the relative Type I as a function of the nominal Type I error rate for all cohorts for 6 MAFs scenarios and with 5 Info (information of the imputed SNP) levels (Fig. S12-S41).

Moreover, given that the simulated cohorts might not reflect real data, we also create “nullified” data based on 20-real GWAS (SCZ, ADHD, MDD, AUT for abbreviation see Table 2, and fourteen not public available), (Experiment 3). This approximation for null data is obtained by substituting the expected quantile of the Gaussian distribution for the (ordered) Z-score. For these simulations, for the size of the test, we add one more MAF scenario and we retain the same Info levels (Fig. S42-S48).

**Table S2. Ethnic composition (weights) for Cohorts from 1000 Genomes haplotypic data.**

| Cohort | ASW | CEU | CHB | CHS | GBR | MXL | PUR | YRI |
|---|---|---|---|---|---|---|---|---|
| 1 | - | 0.30 | - | 0.25 | - | - | 0.05 | 0.40 |
| 2 | 0.10 | 0.15 | 0.15 | 0.125 | 0.15 | 0.10 | 0.025 | 0.20 |
| 3 | 0.15 | - | 0.35 | - | 0.35 | 0.15 | - | - |
| 4 | 0.45 | - | - | - | 0.55 | - | - | - |
| 5 | - | - | 0.55 | - | - | 0.45 | - | - |

Where CEU is the abbreviation for Utah Residents (CEPH) with Northern and Western European Ancestry, CHB for Han Chinese in Beijing, China, CHS for Southern Han Chinese.

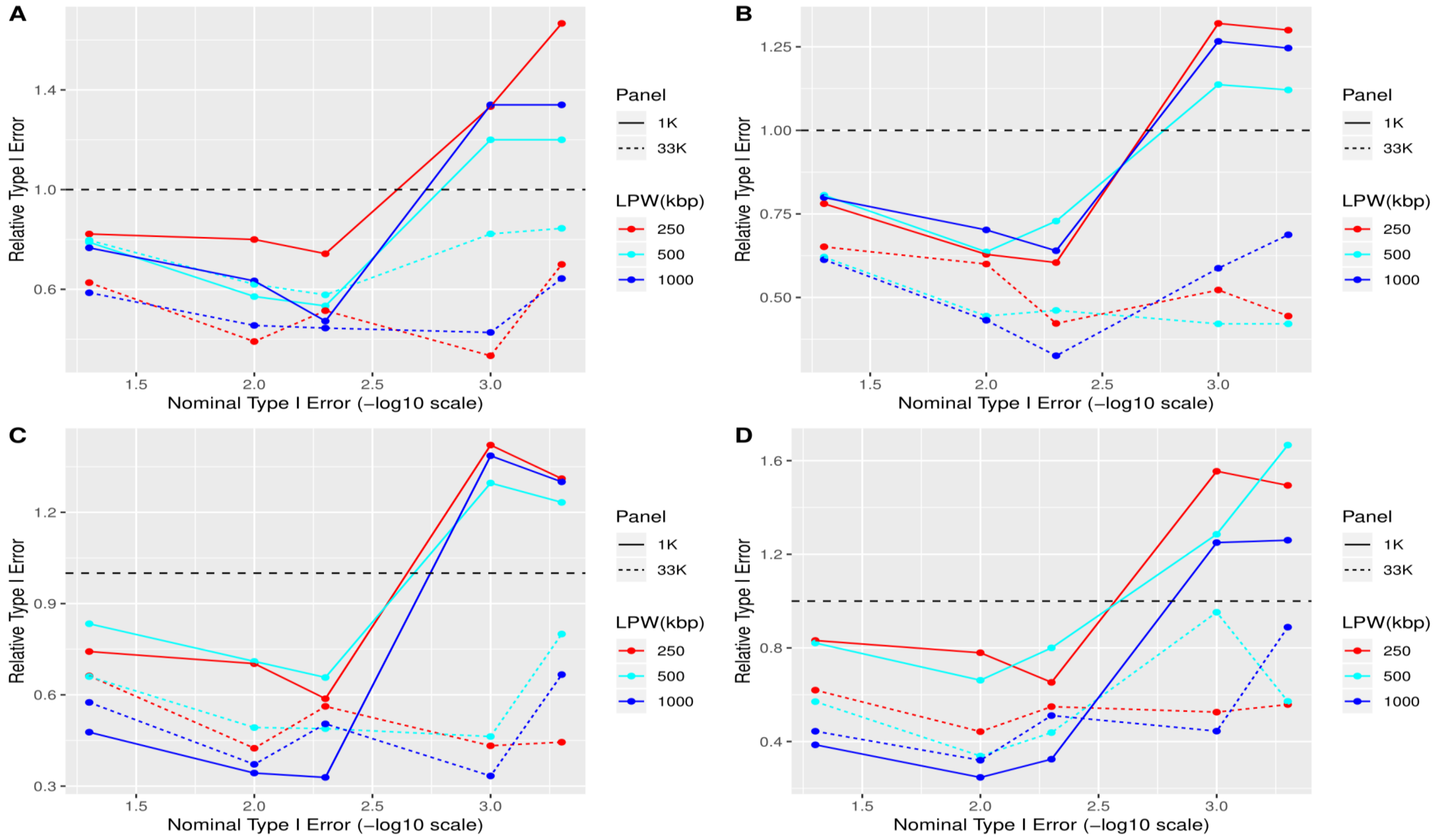


**Fig. S1. Relative size of the test (the quotient of empirical false positive rate and nominal type I error), for masked Illumina 1M markers in all cohorts. 1A designates MAF$< 5\%$, 1B - $5\% <$MAF$< 10\%$, 1C - $10\% <$MAF$< 20\%$ and 1D - MAF$> 20\%$. In legend, panel designates whether the statistic was under 1K or 33K reference panel. LPW(kbp) denotes the length of the prediction window measured in in thousands base pairs (kbp).**

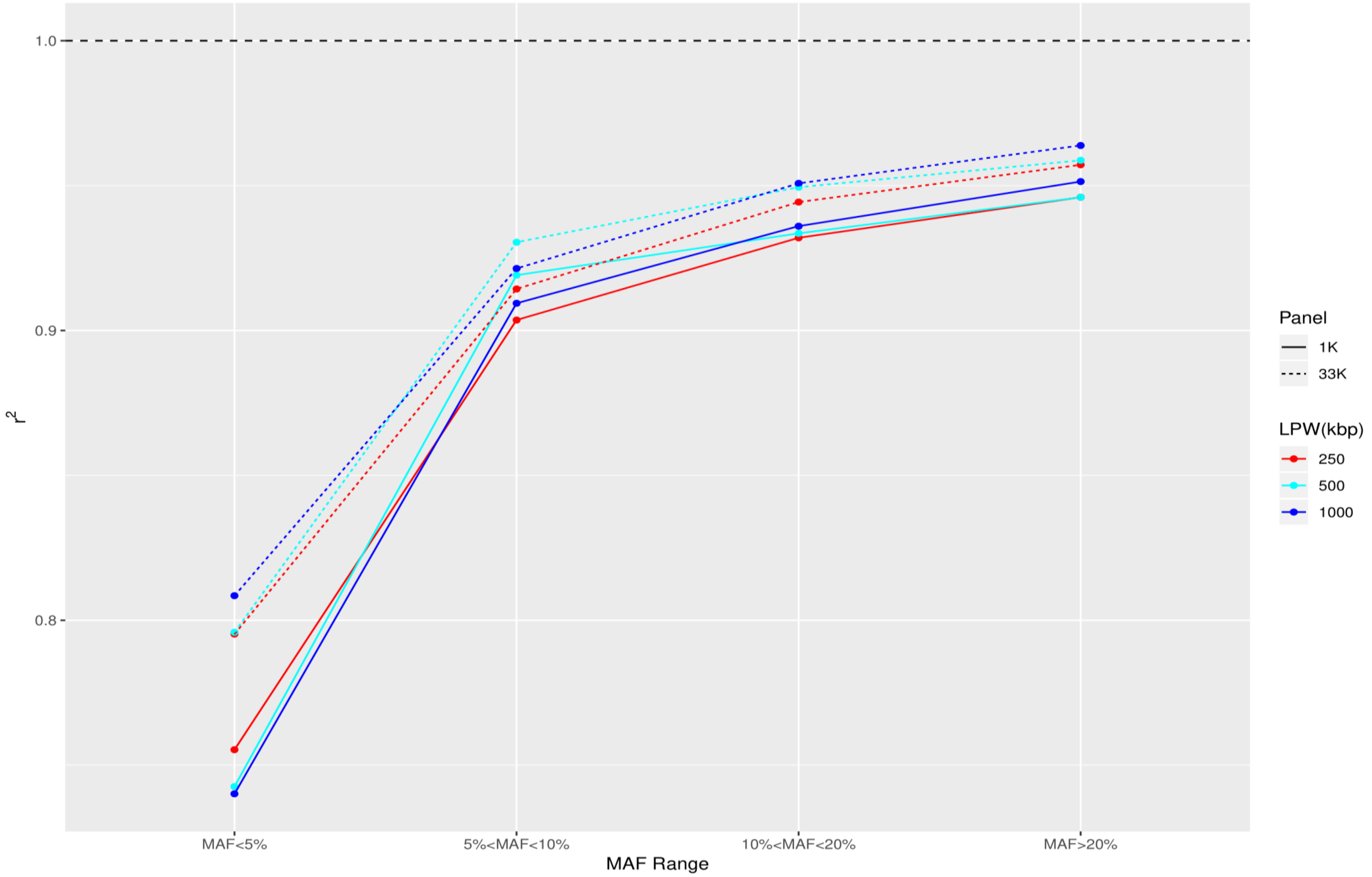


**Fig. S2 Correlation estimates beteen true and imputed Z-scores for Cohort 1 (30% CEU + 25% CHS + 5% PUR + 40% YRI). See Fig S1 for background and abbreviations**

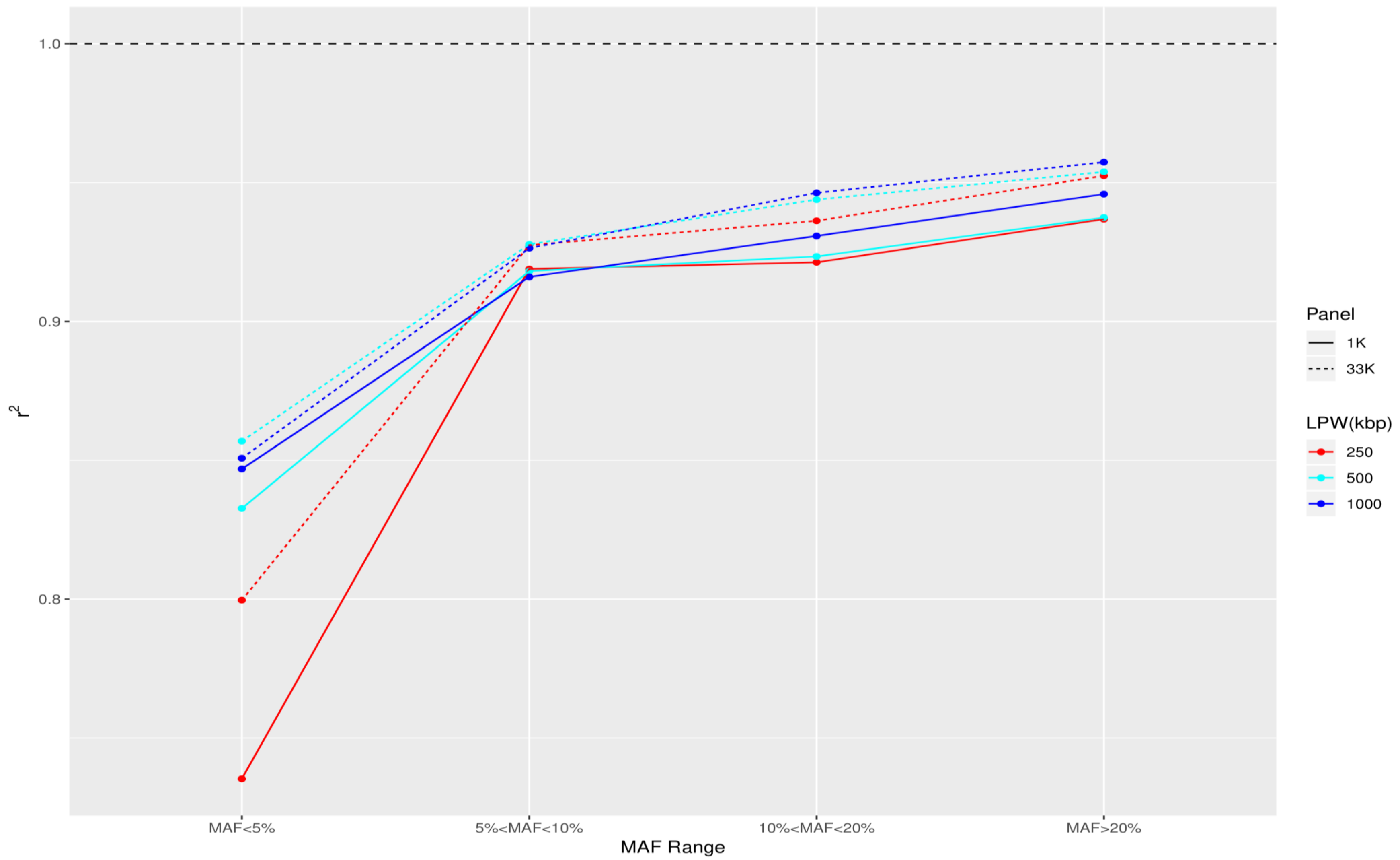


**Fig. S3. Correlation estimates beteen true and imputed Z-scores for Cohort 2 (10% ASW + 15% CEU + 15% CHB + 12.5% CHS + 15% GBR + 10% MXL + 2.5% PUR + 20% YRI). See Fig S1 for background and abbreviations.**

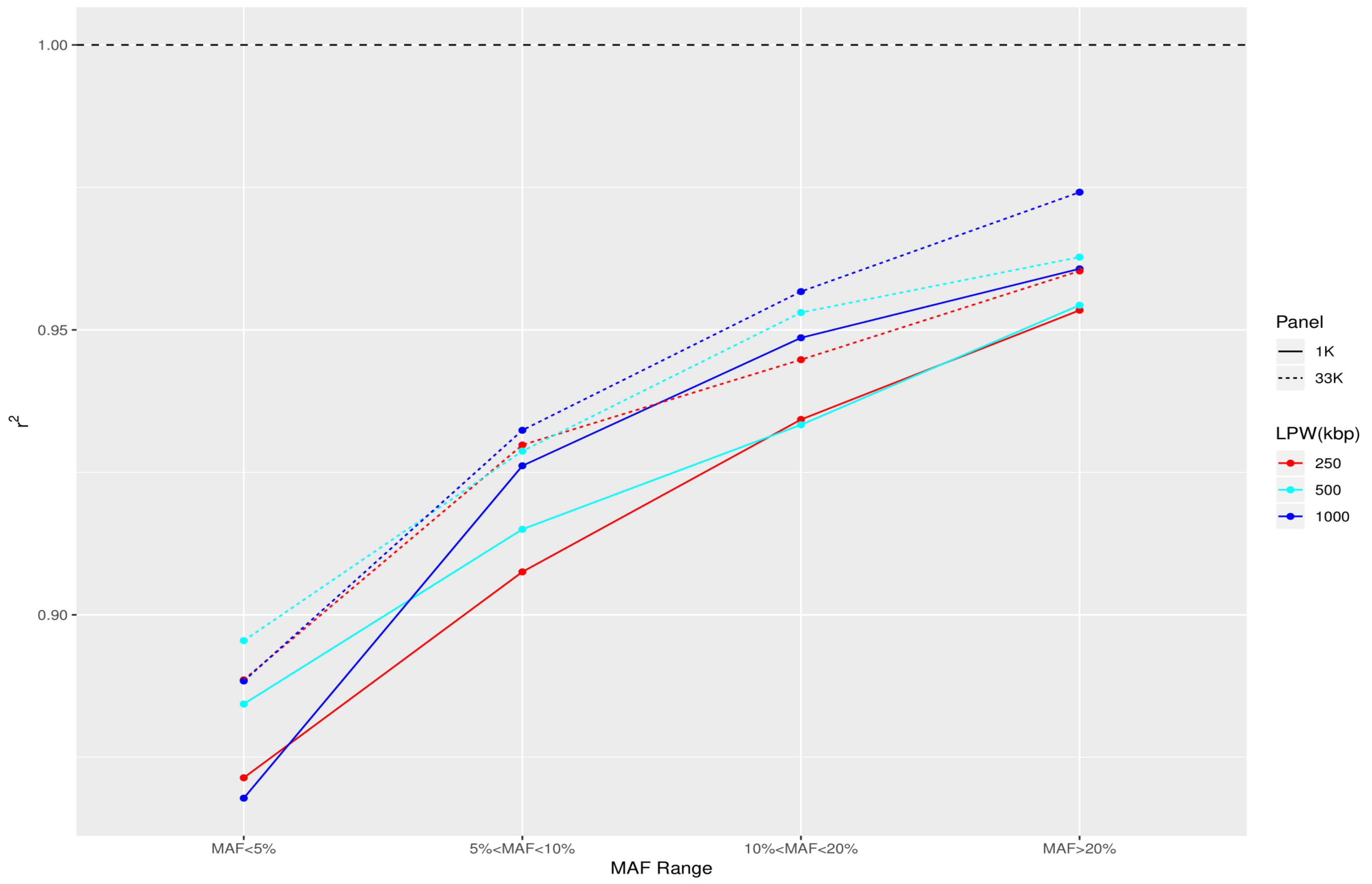


**Fig. S4. Correlation estimates beteen true and imputed Z-scores for Cohort 3 (15% ASW + 35% CHB + 35% GBR + 15% MXL). See Fig S1 for background and abbreviations.**

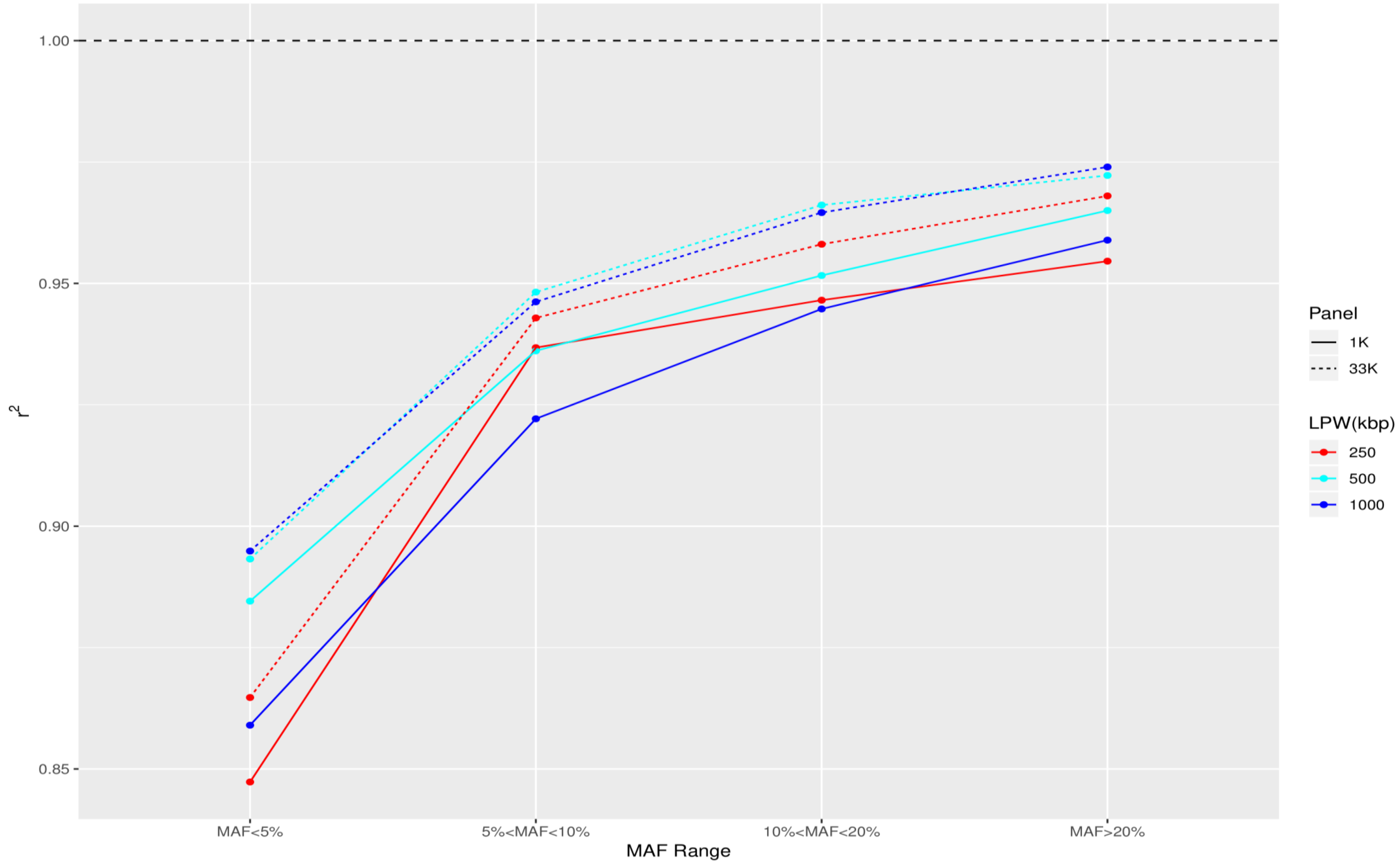


**Fig. S5. Correlation estimates beteen true and imputed Z-scores for Cohort 4 (45% ASW + 55% GBR). See Fig S1 for background and abbreviations.**

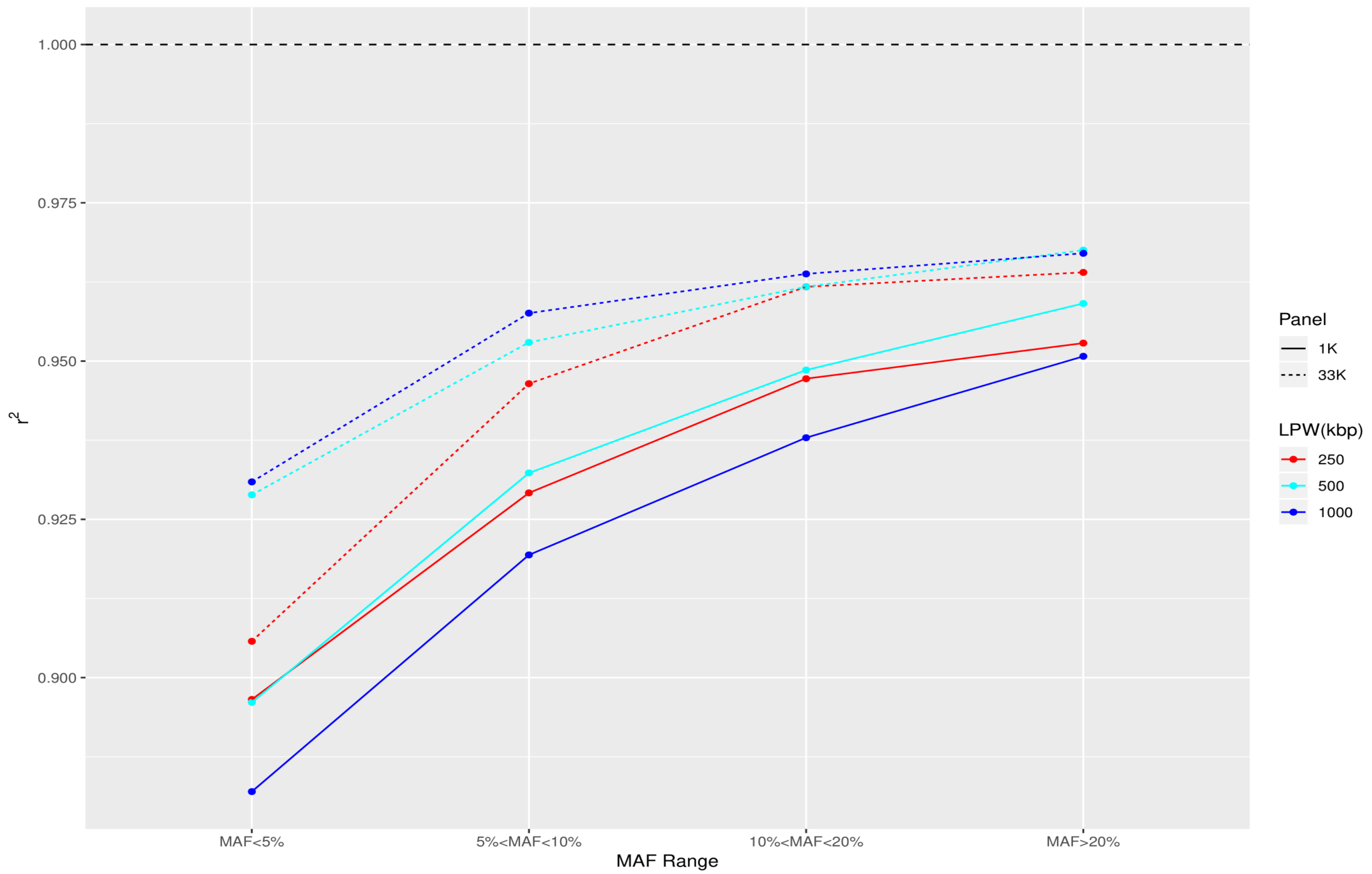


**Fig. S6. Correlation estimates beteen true and imputed Z-scores for Cohort 5 (55% CHB + 45% MXL). See Fig S1 for background and abbreviations.**

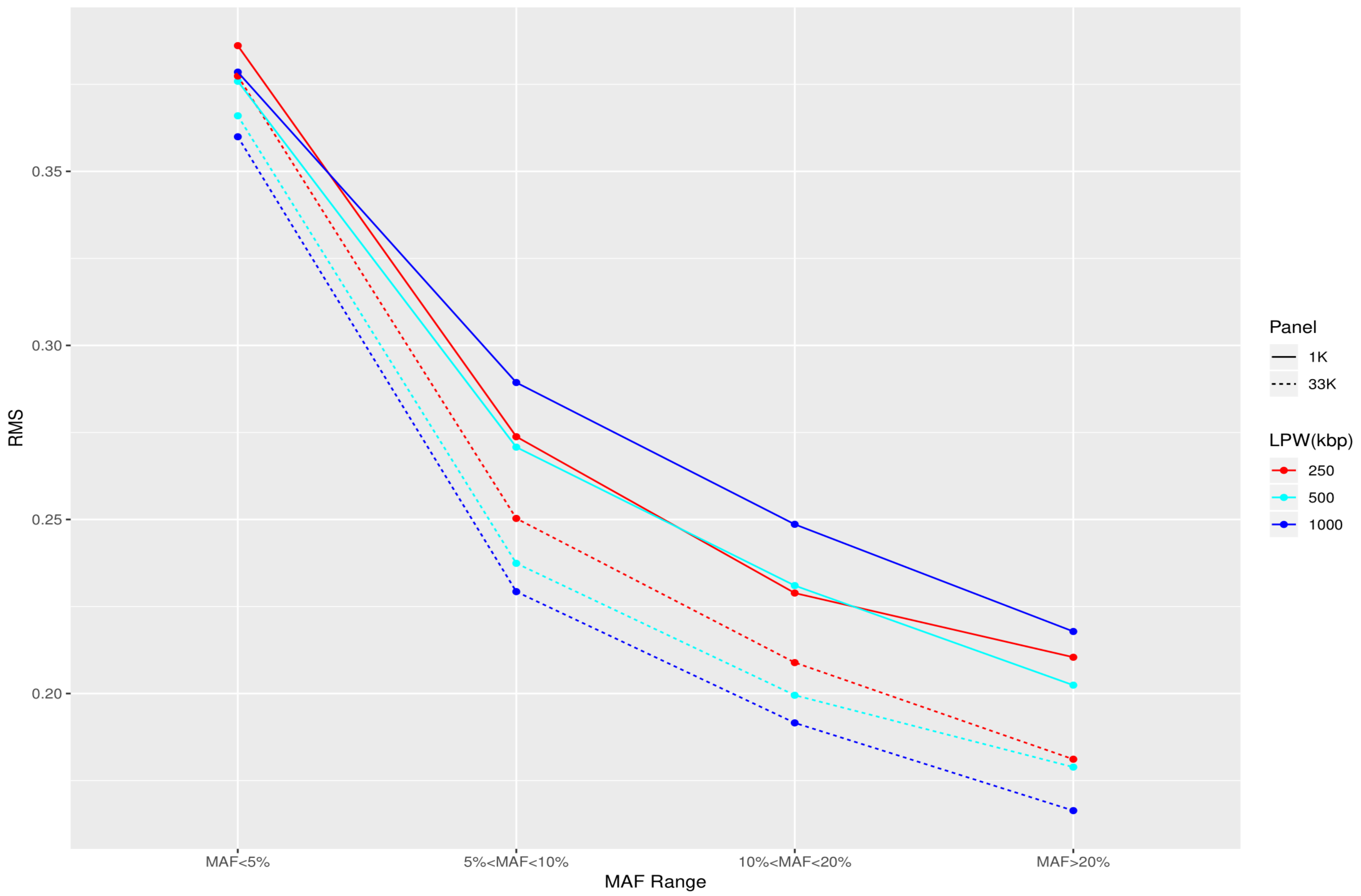


**Fig. S7. RMS of imputed Z-scores around their true value for Cohort 1 ancestry. See Fig S1 for background and abbreviations.**

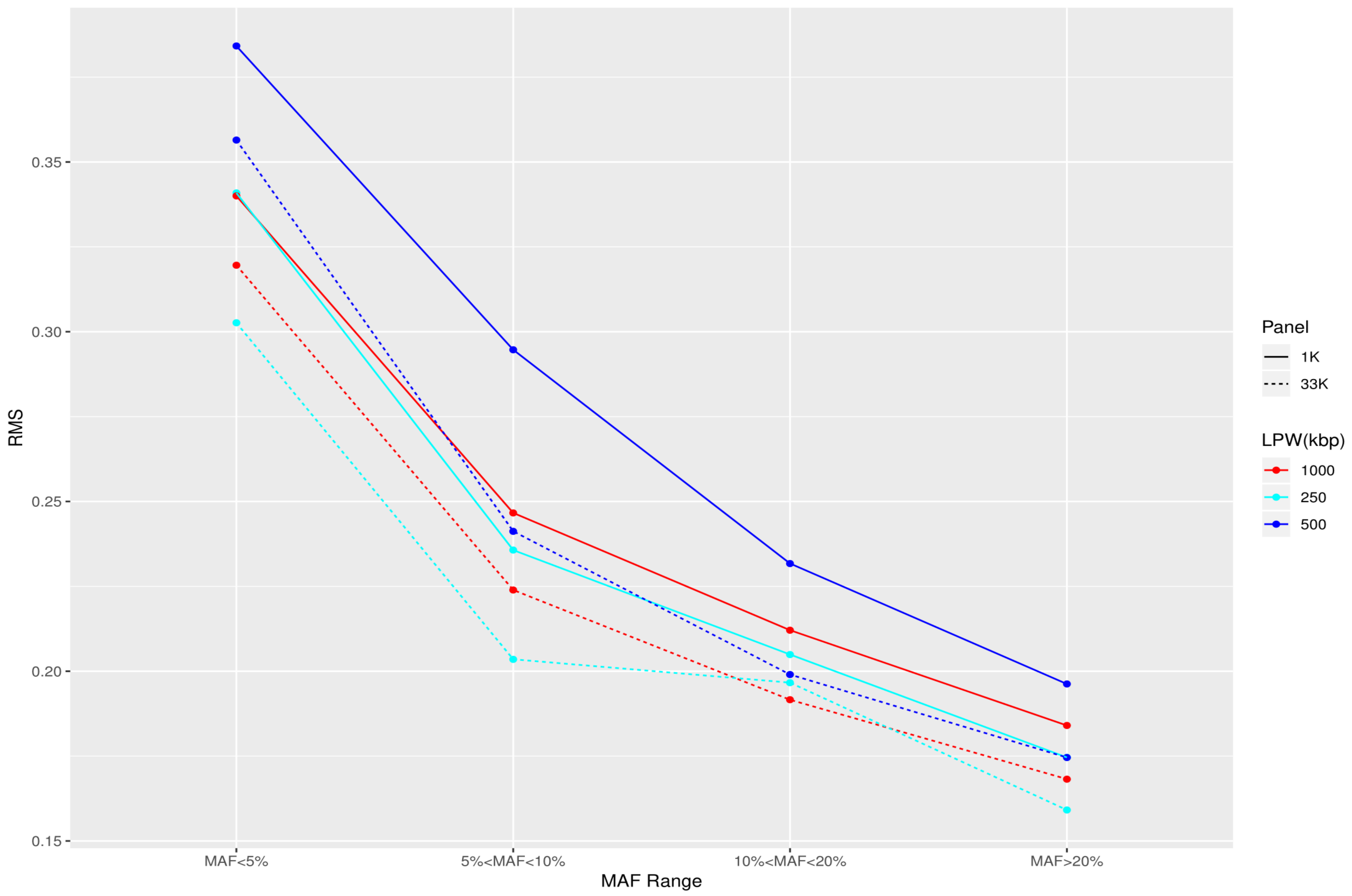


**Fig. S8. RMS of imputed Z-scores around their true value for Cohort 2. See Fig S1 for background and abbreviations.**

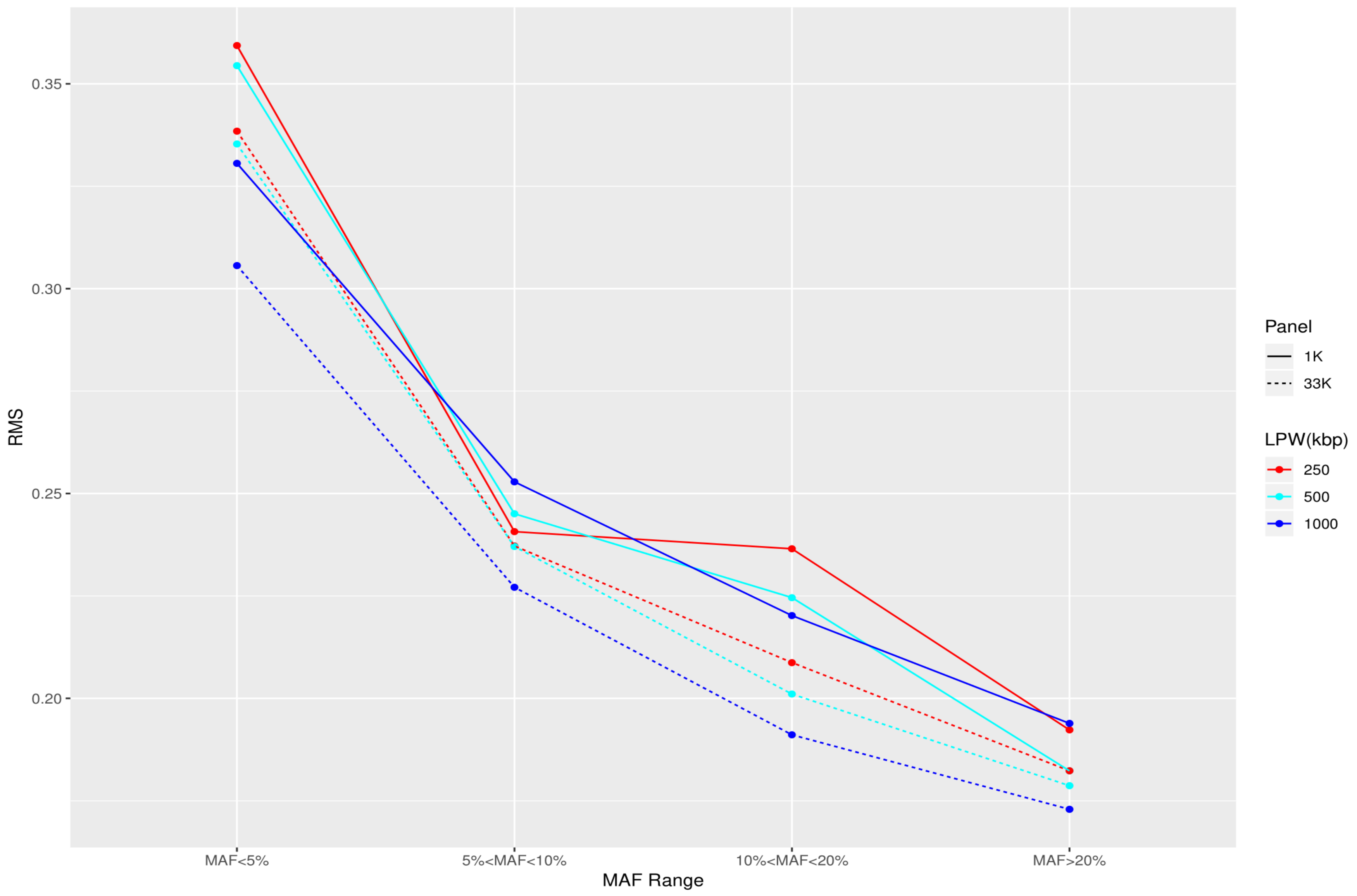


**Fig. S9. RMS of imputed Z-scores around their true value for Cohort 3. See Fig S1 for background and abbreviations.**

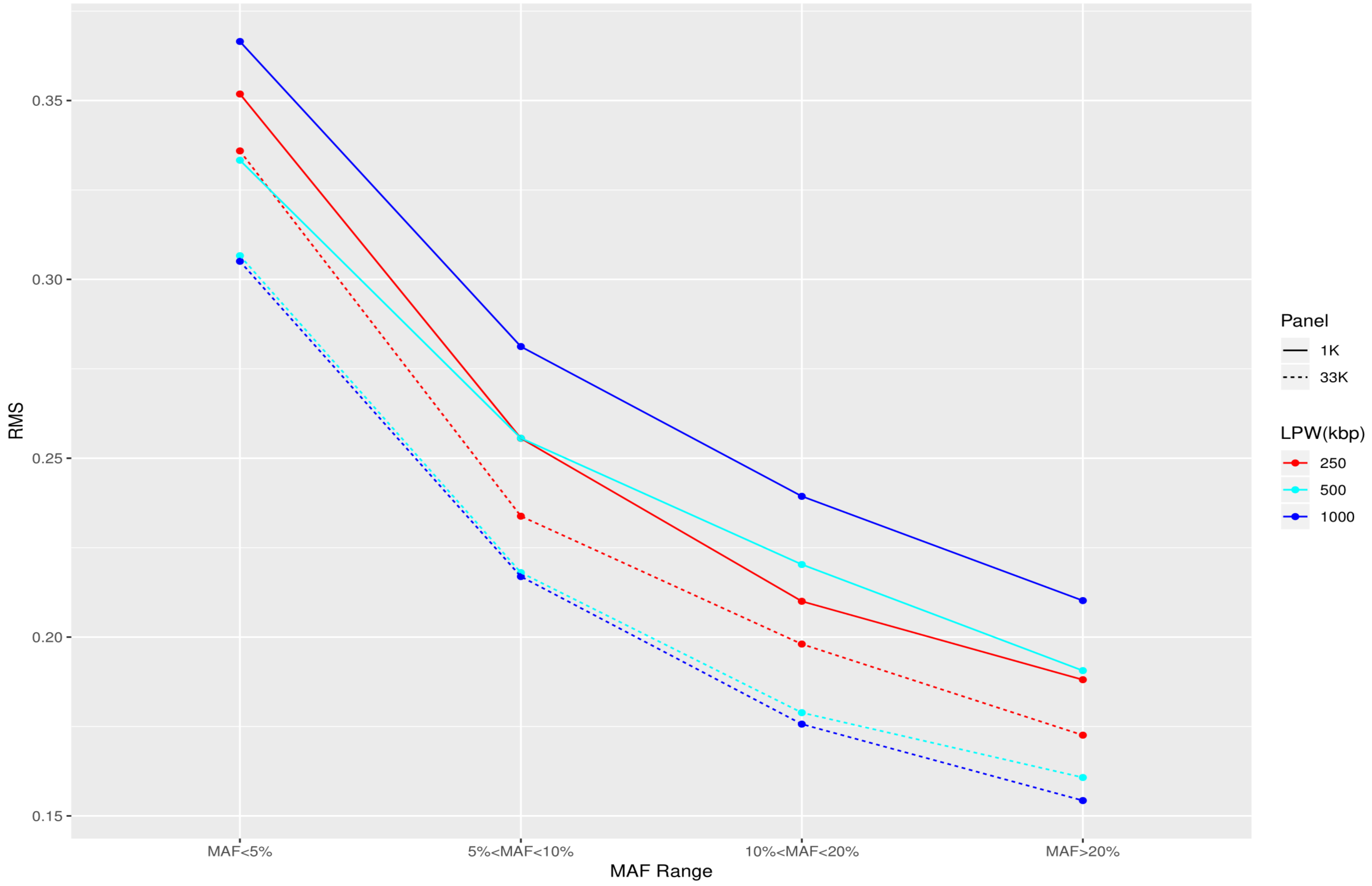


**Fig. S10. RMS of imputed Z-scores around their true value for Cohort 4. See Fig S1 for background and abbreviations.**

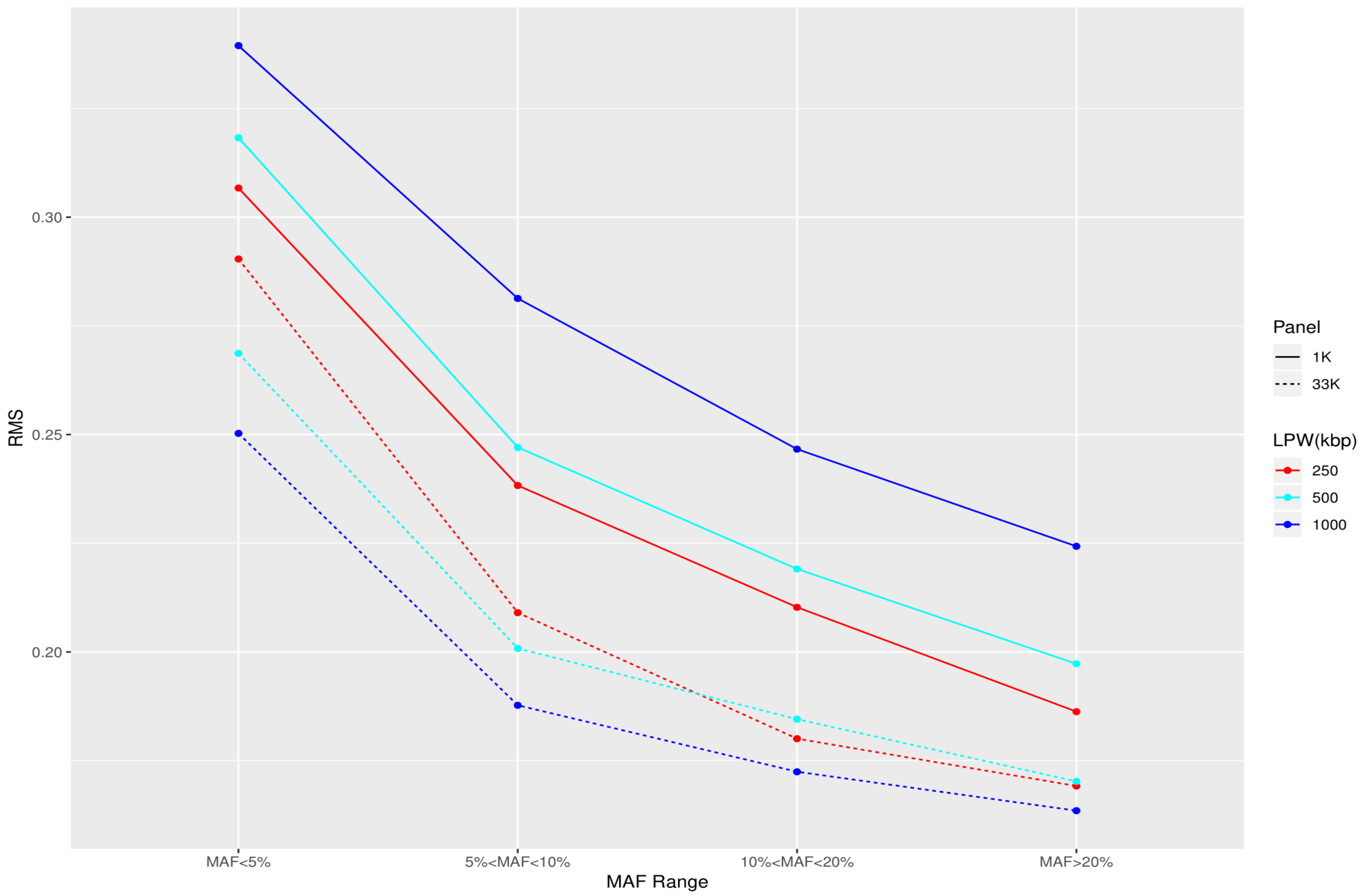


**Fig. S11. RMS of imputed Z-scores around their true value for Cohort 5. See Fig S1 for background and abbreviations.**

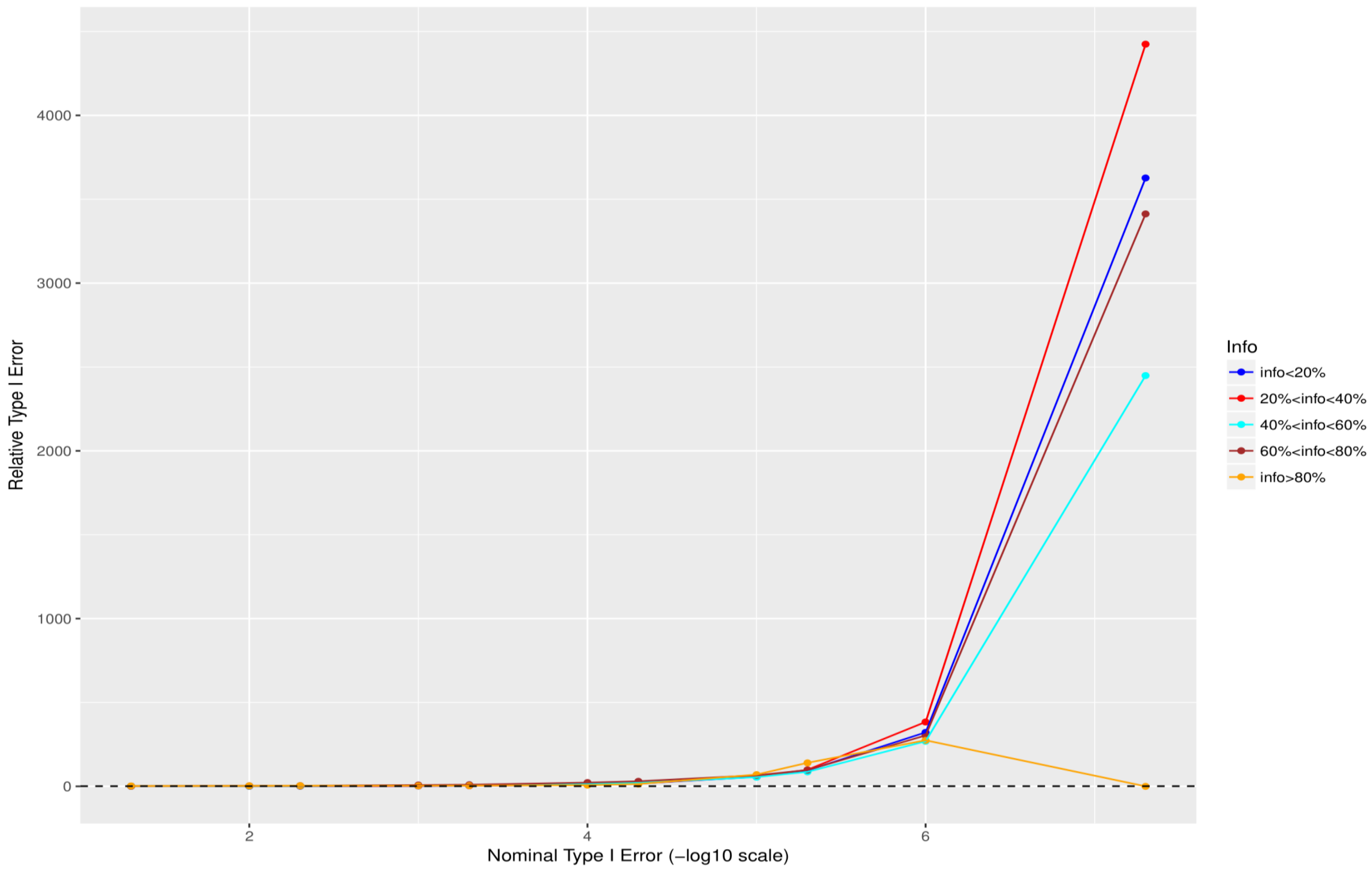


**Fig. S12 Relative size of the test - Cohort 1 for 0.05%<MAF<0.5% (the quotient of empirical false positive rate and nominal type I error).**

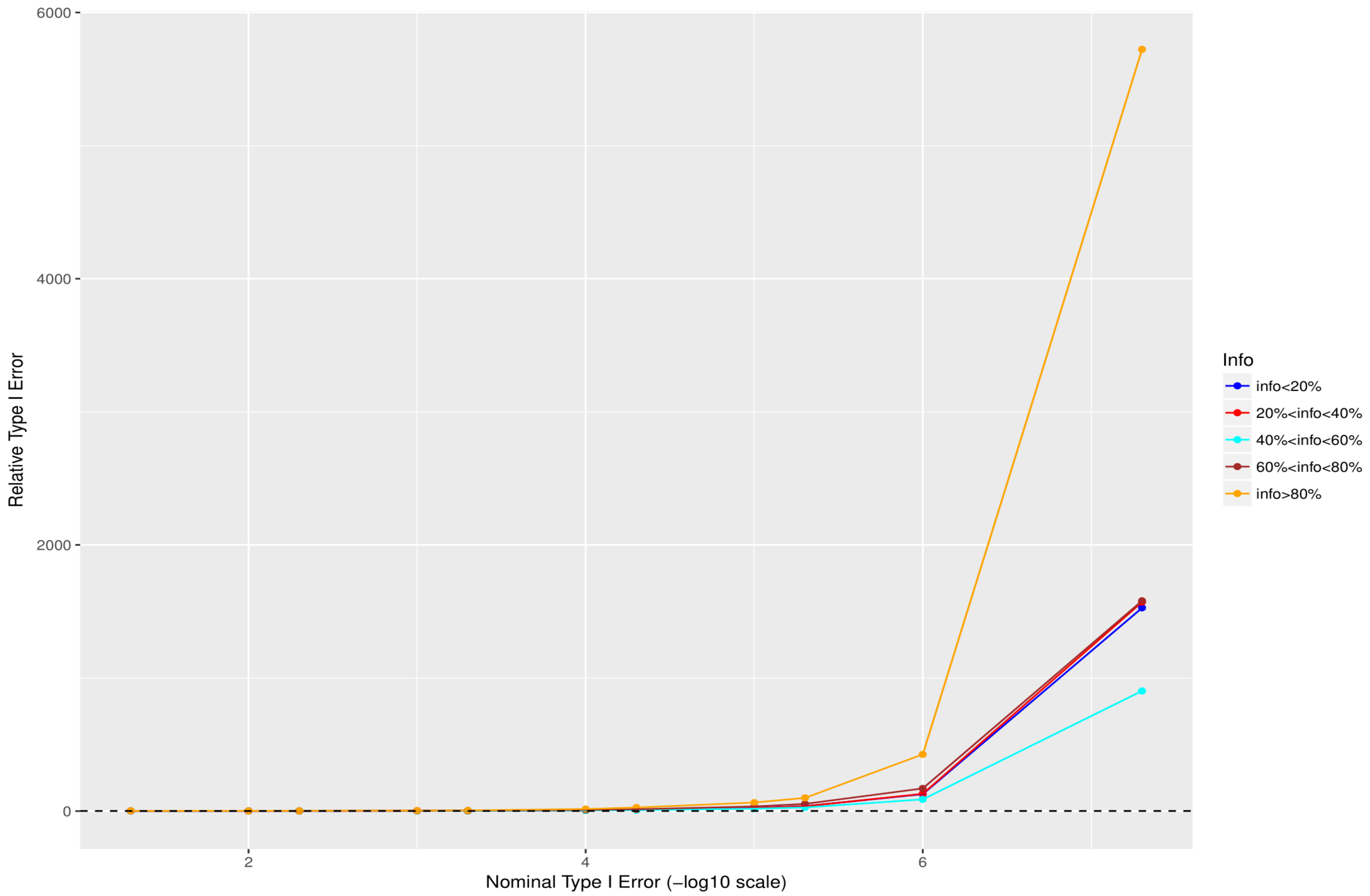


**Fig. S13 Relative size of the test - Cohort 1 for 0. 5%<MAF<1% (the quotient of empirical false positive rate and nominal type I error).**

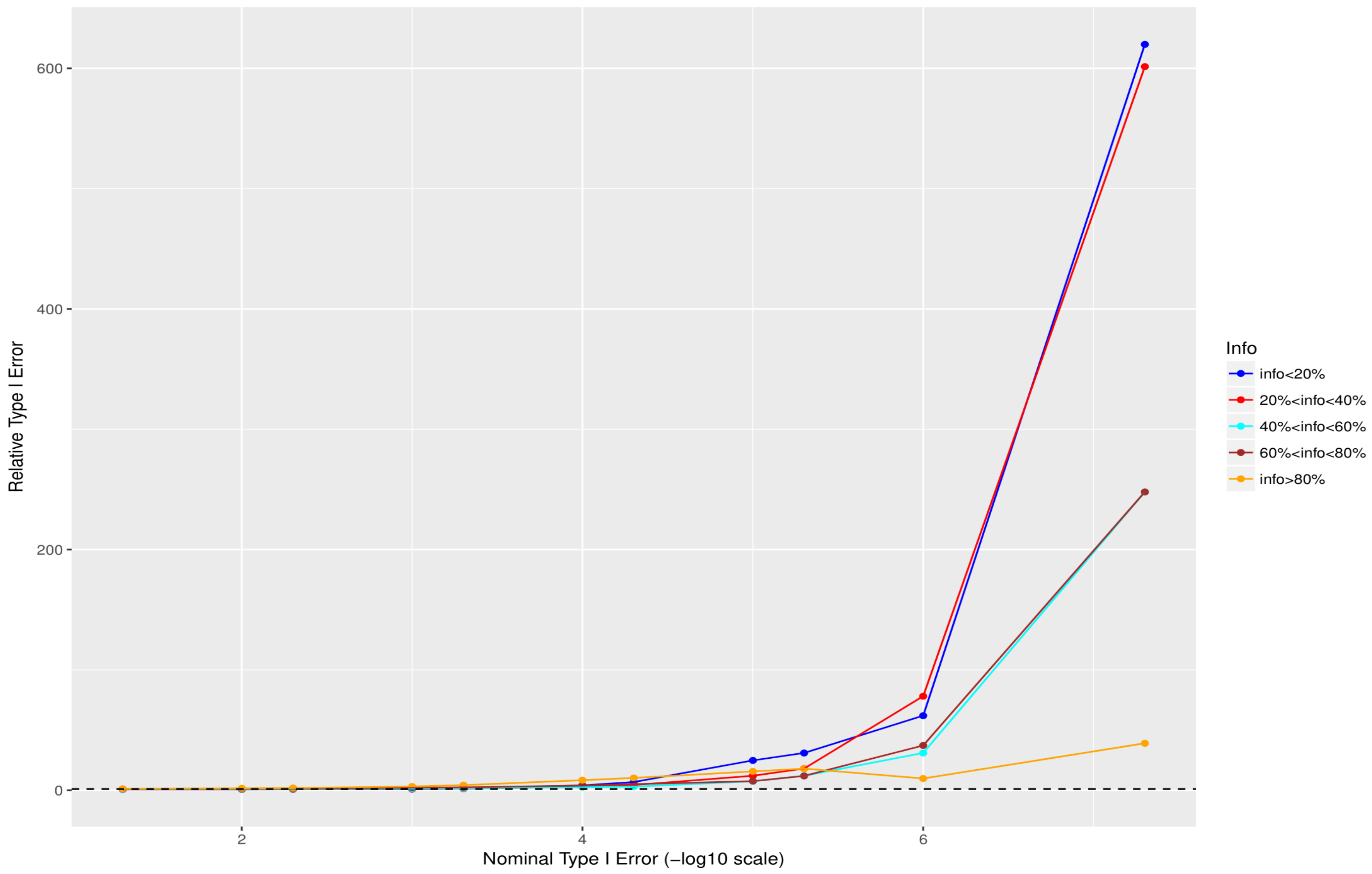


**Fig. S14 Relative size of the test - Cohort 1 for 1%<MAF<2% (the quotient of empirical false positive rate and nominal type I error).**

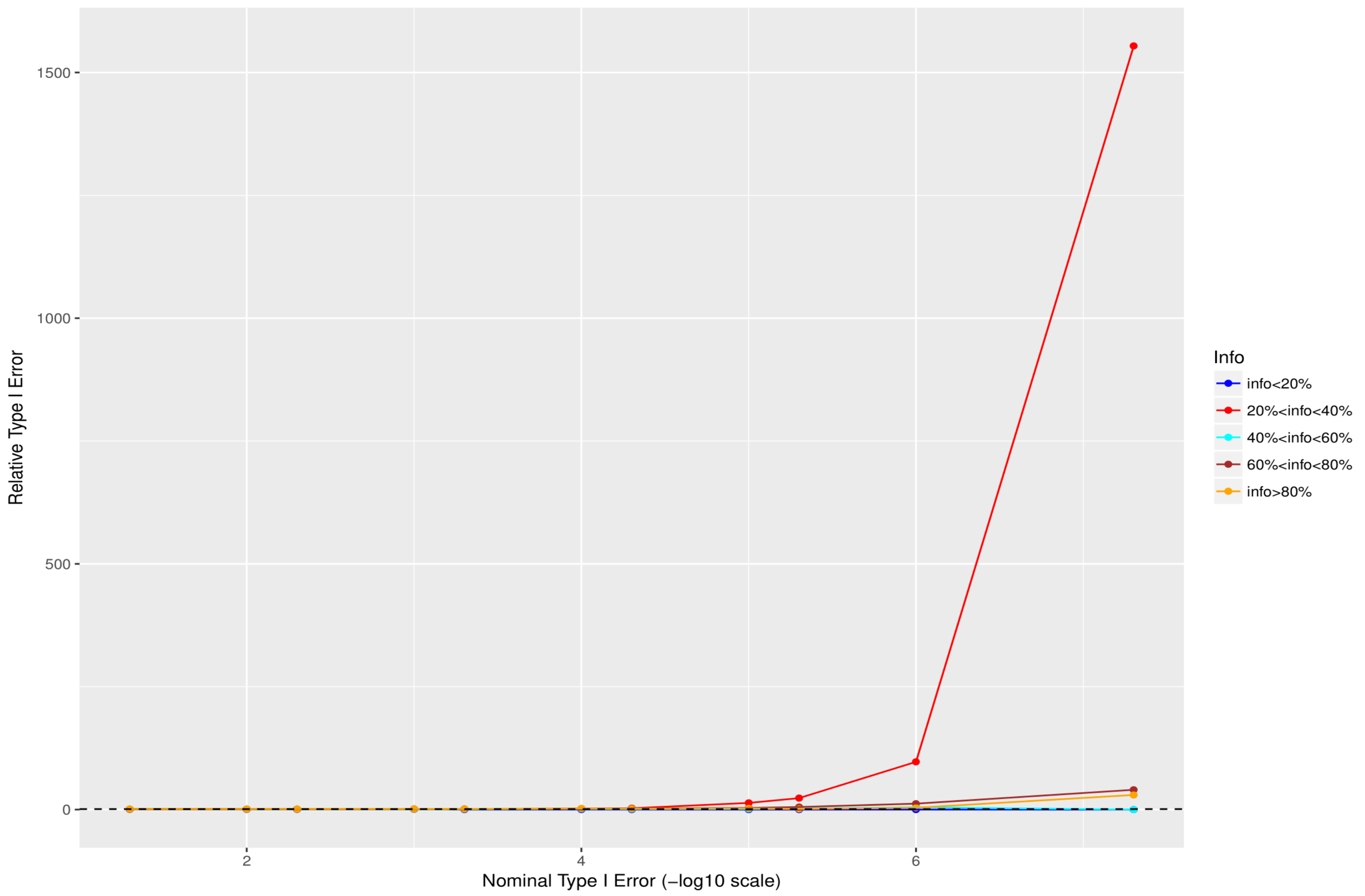


**Fig. S15 Relative size of the test - Cohort 1 for 2%<MAF<5% (the quotient of empirical false positive rate and nominal type I error).**

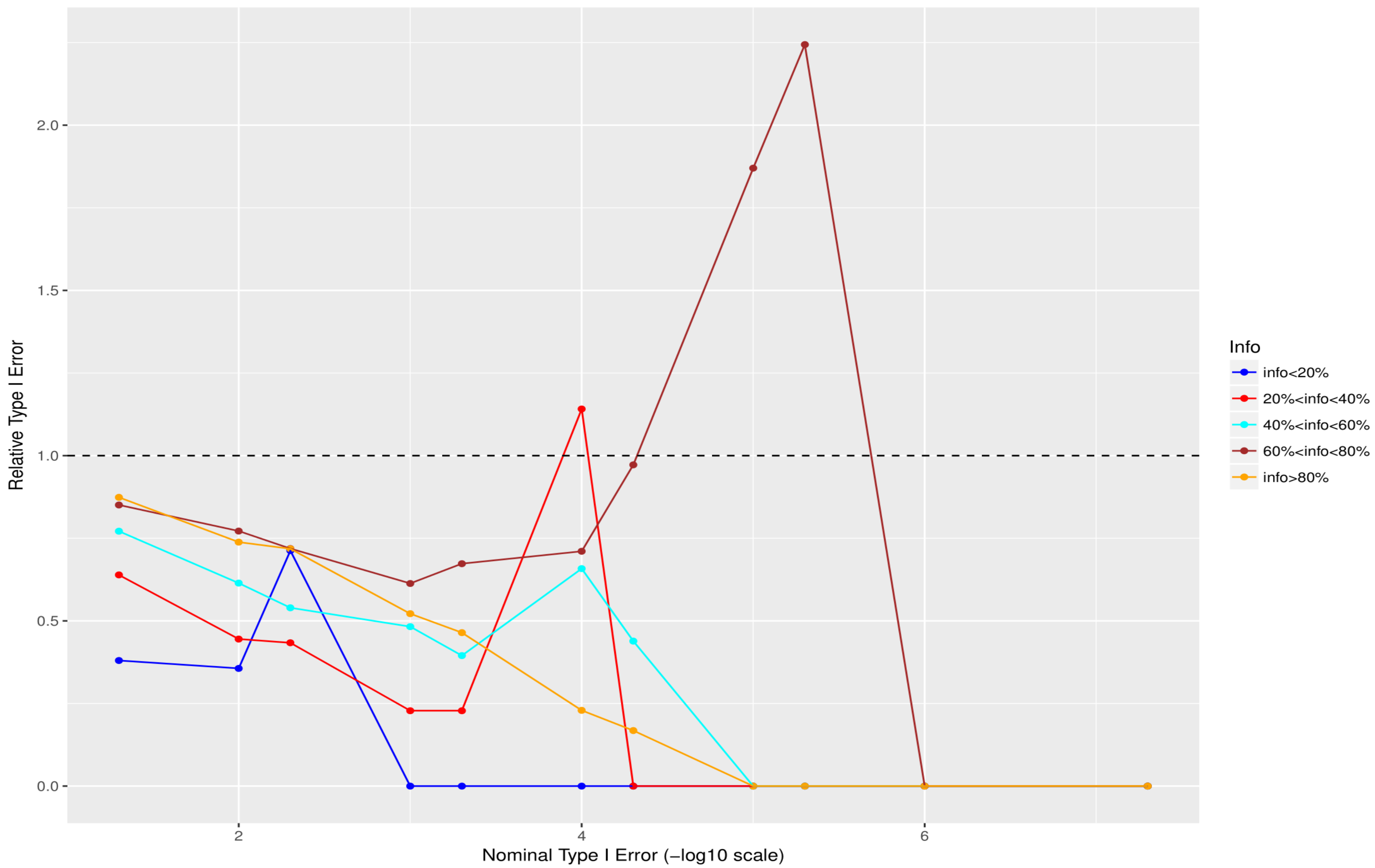


**Fig. S16 Relative size of the test - Cohort 1 for 5%<MAF<10% (the quotient of empirical false positive rate and nominal type I error).**

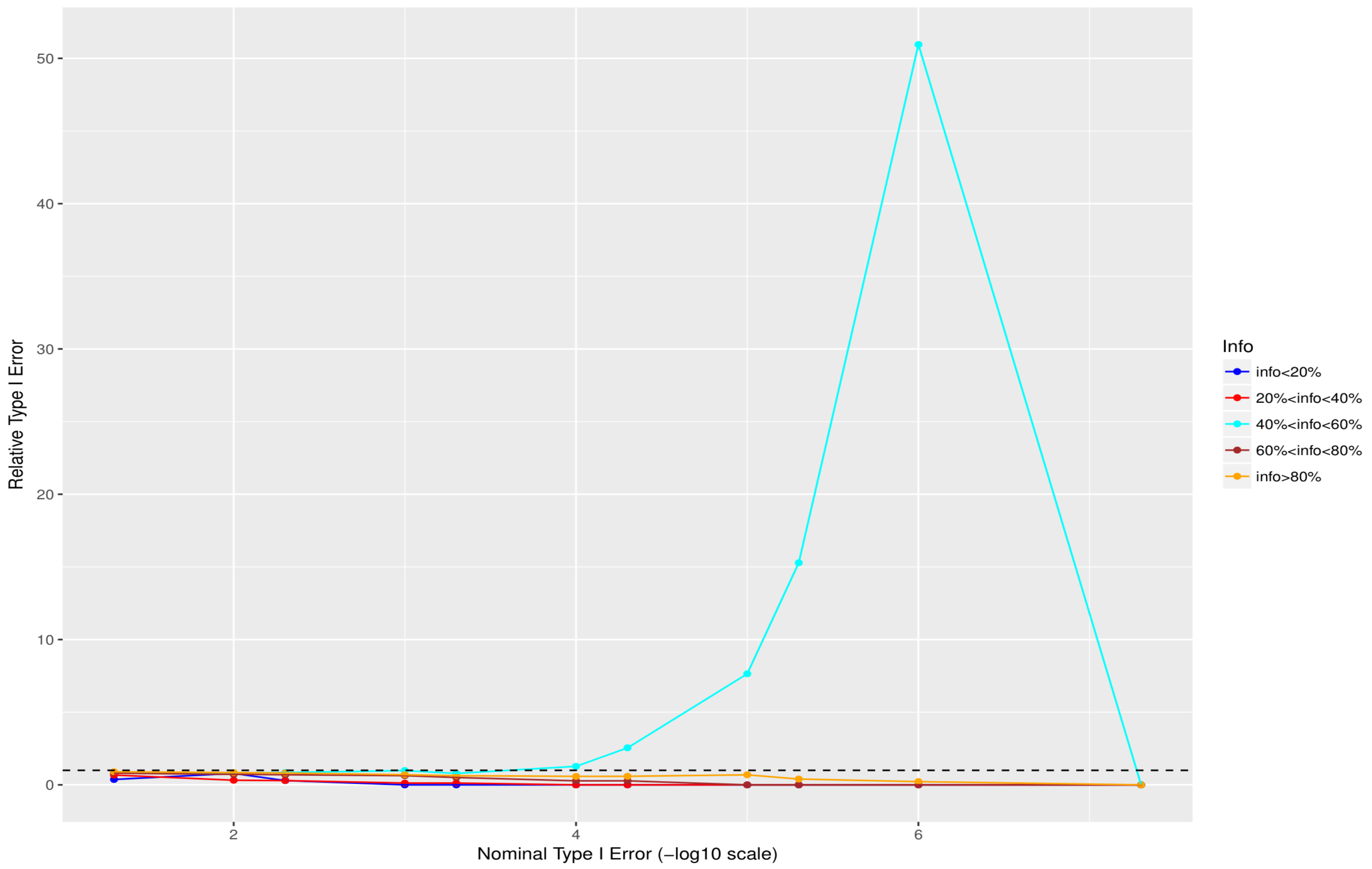


**Fig. S17 Relative size of the test - Cohort 1 for 10%<MAF<50% (the quotient of empirical false positive rate and nominal type I error).**

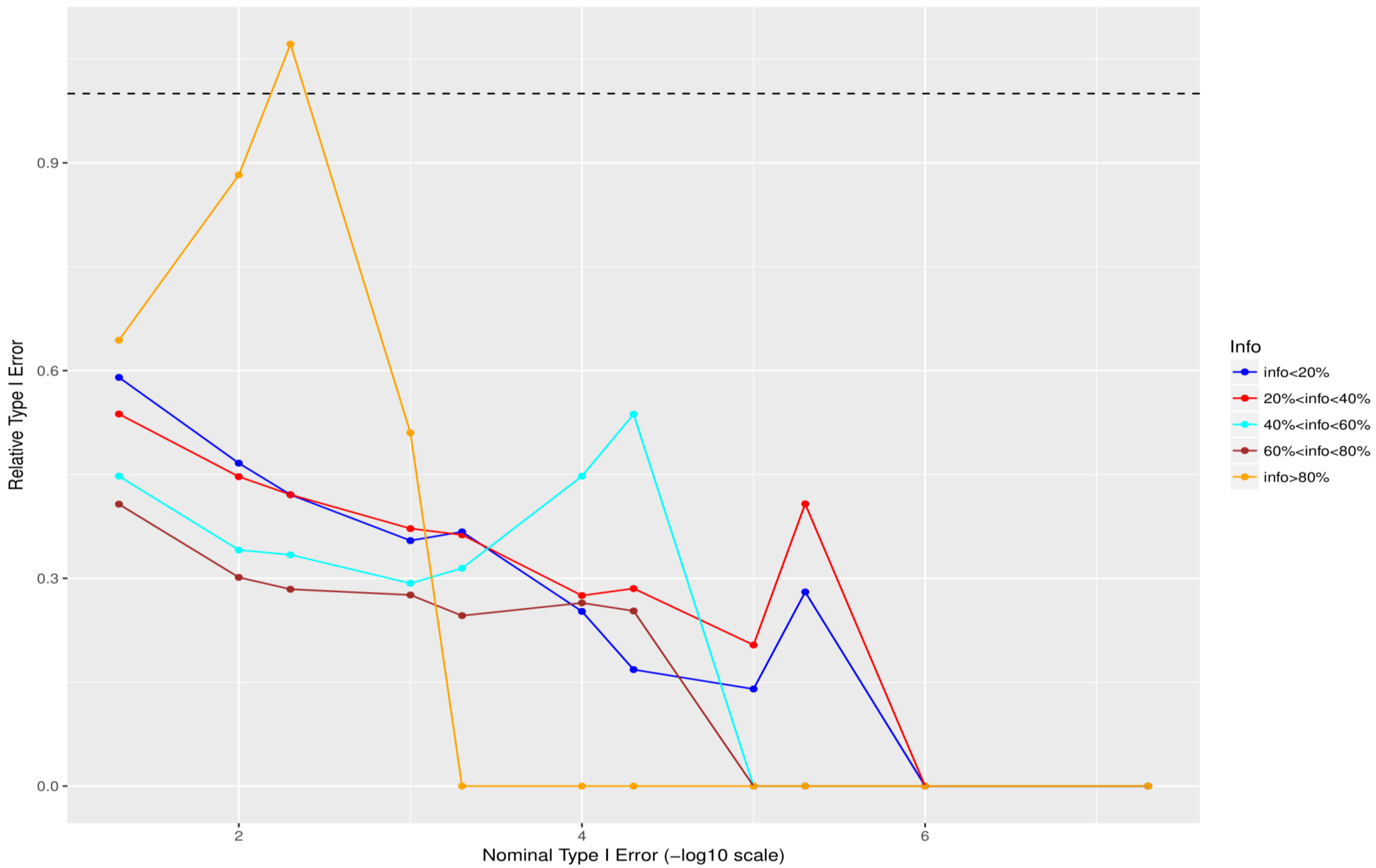


**Fig. S18 Relative size of the test - Cohort 2 for 0.05%<MAF<0.5% (the quotient of empirical false positive rate and nominal type I error).**

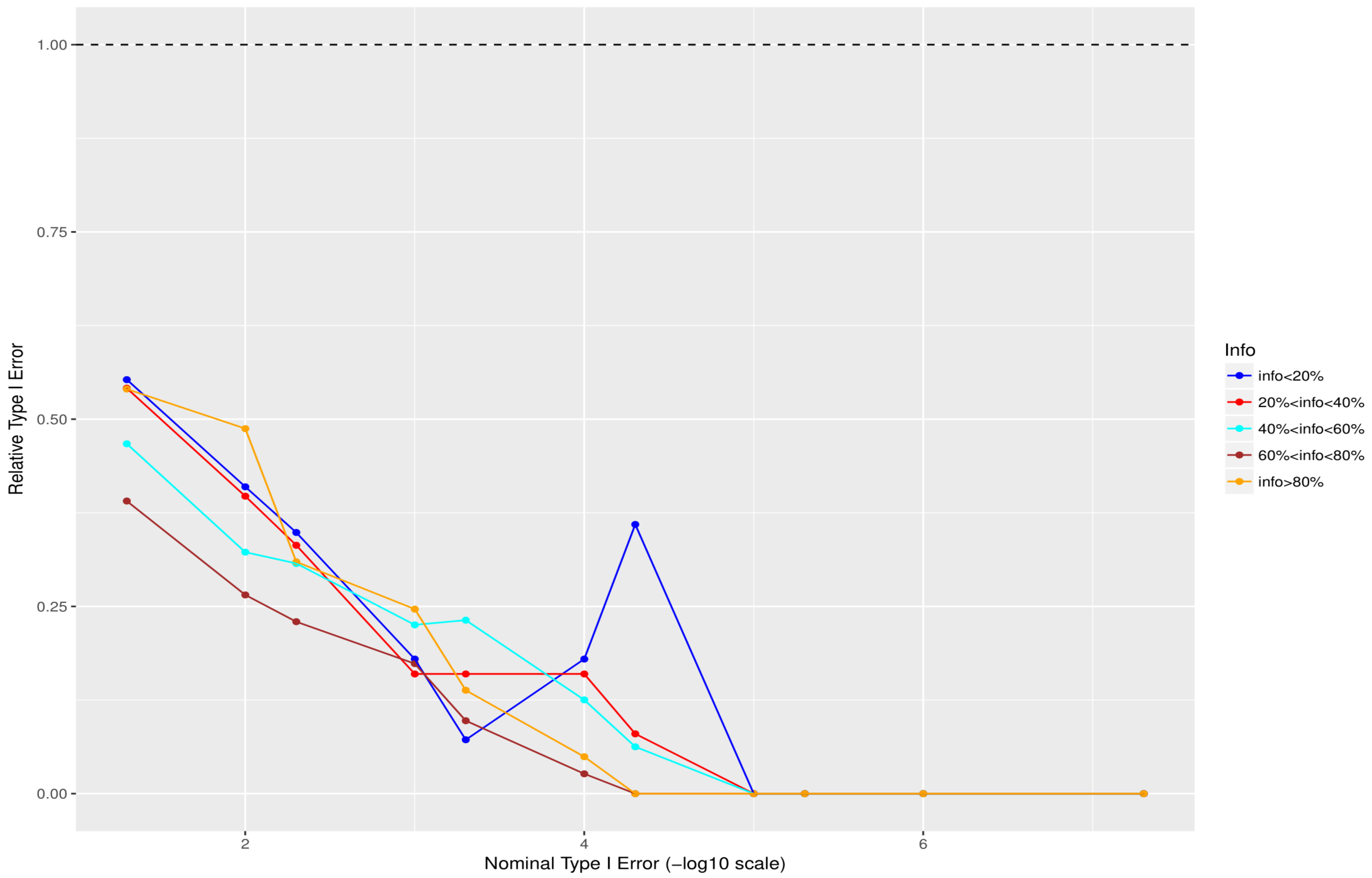


**Fig. S19 Relative size of the test - Cohort 2 for 0. 5%<MAF<1% (the quotient of empirical false positive rate and nominal type I error).**

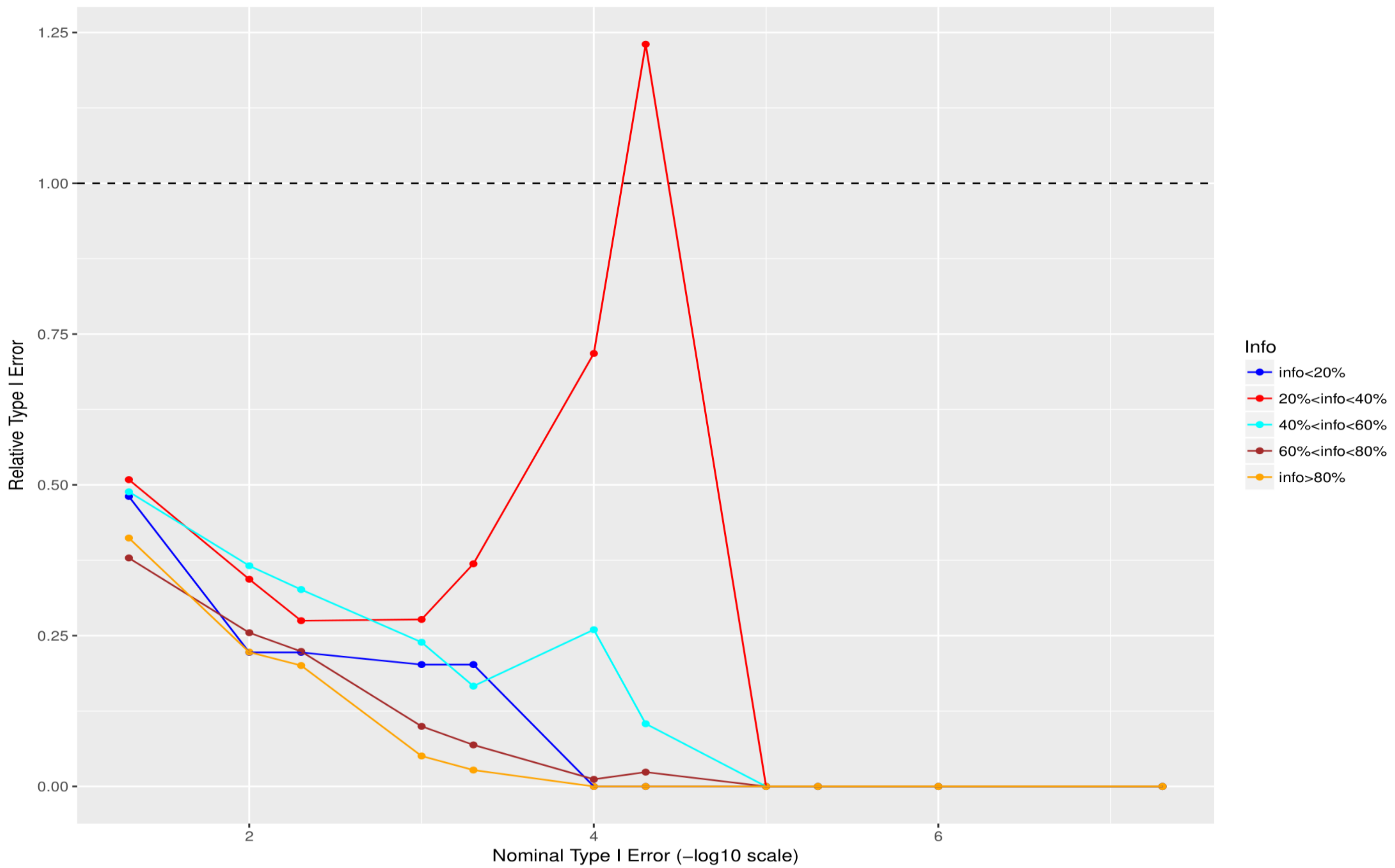


**Fig. S20 Relative size of the test - Cohort 2 for 1%<MAF<2% (the quotient of empirical false positive rate and nominal type I error).**

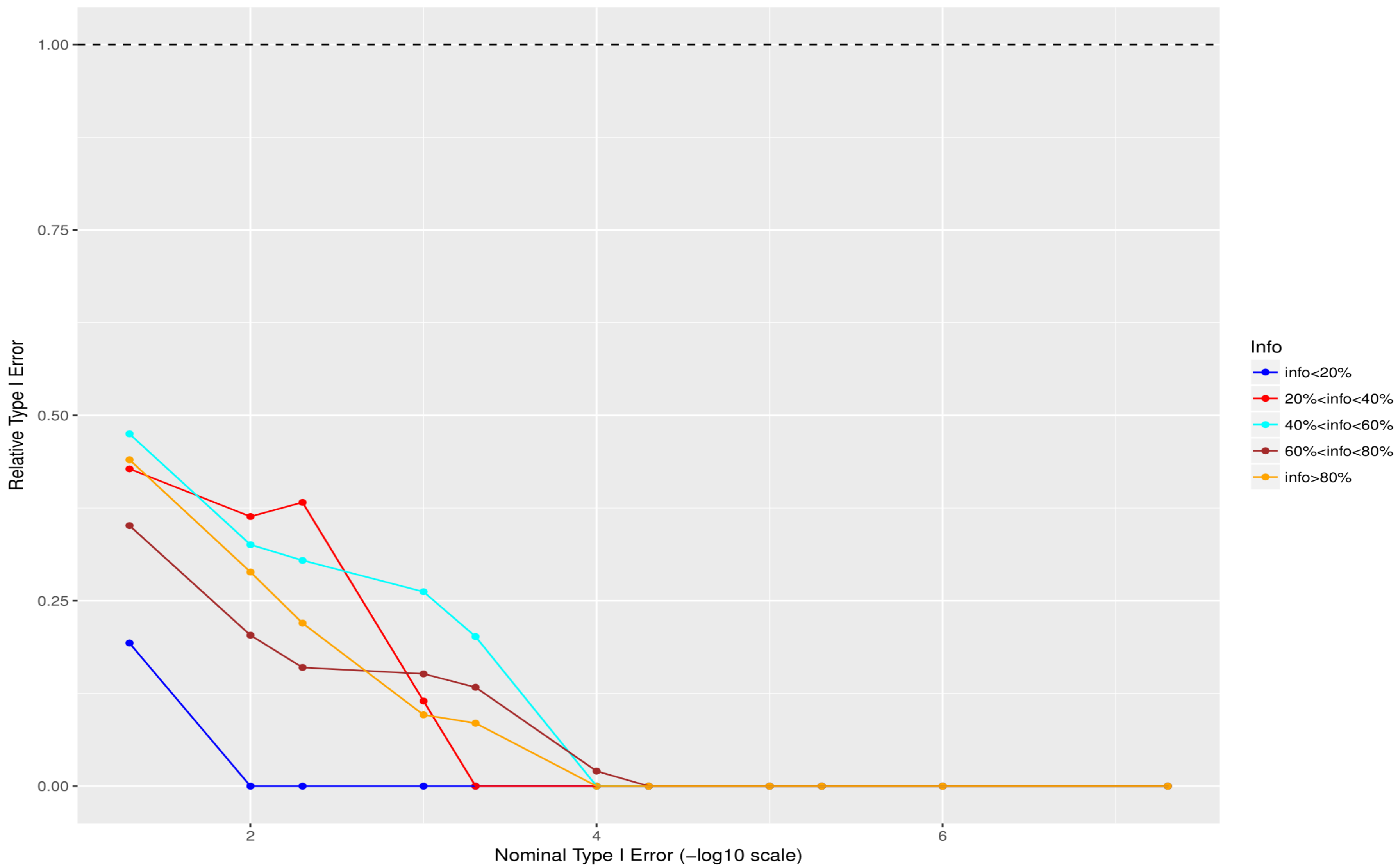


**Fig. S21 Relative size of the test - Cohort 2 for 2%<MAF<5% (the quotient of empirical false positive rate and nominal type I error).**

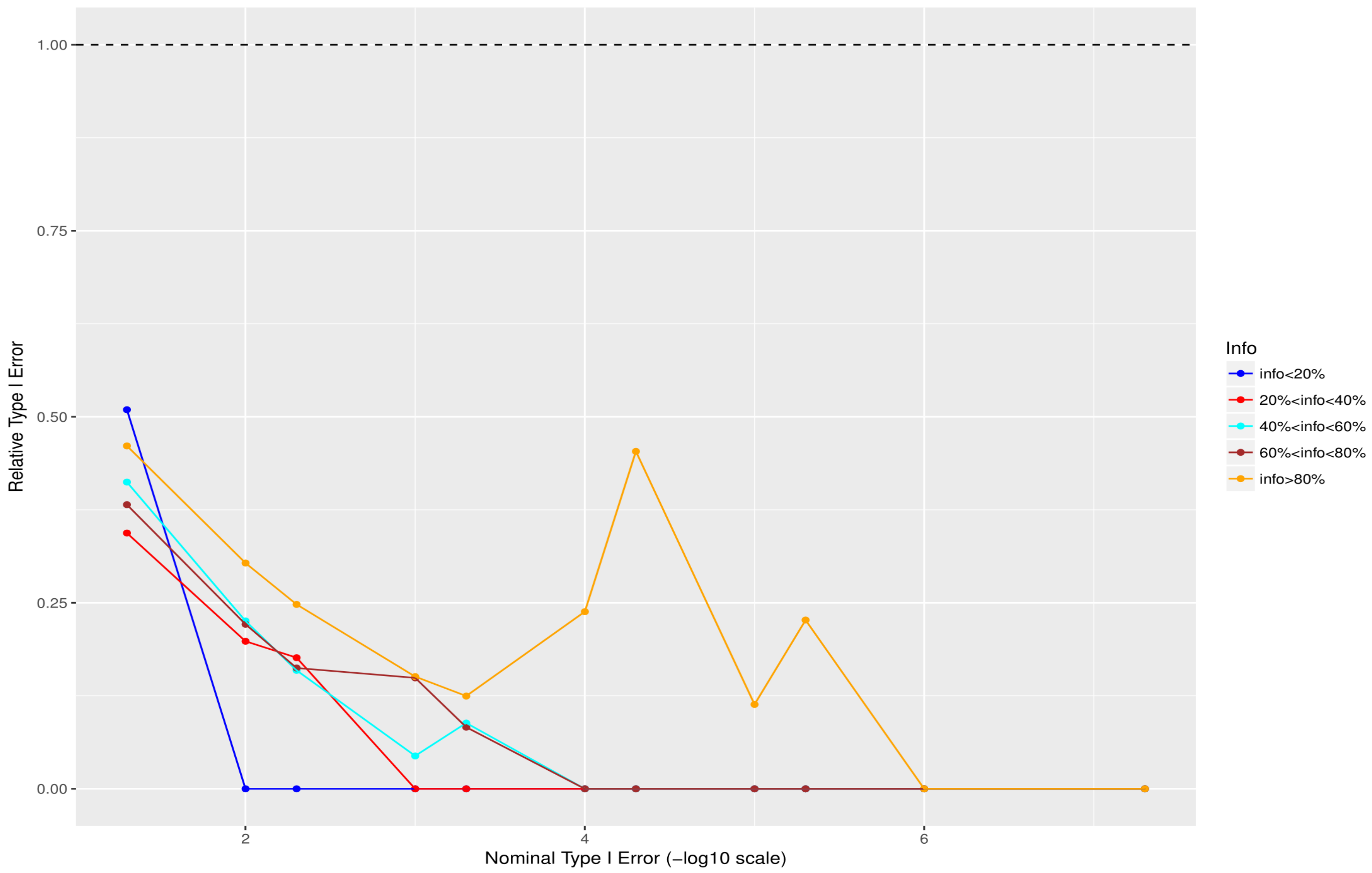


**Fig. S22 Relative size of the test - Cohort 2 for 5%<MAF<10% (the quotient of empirical false positive rate and nominal type I error).**

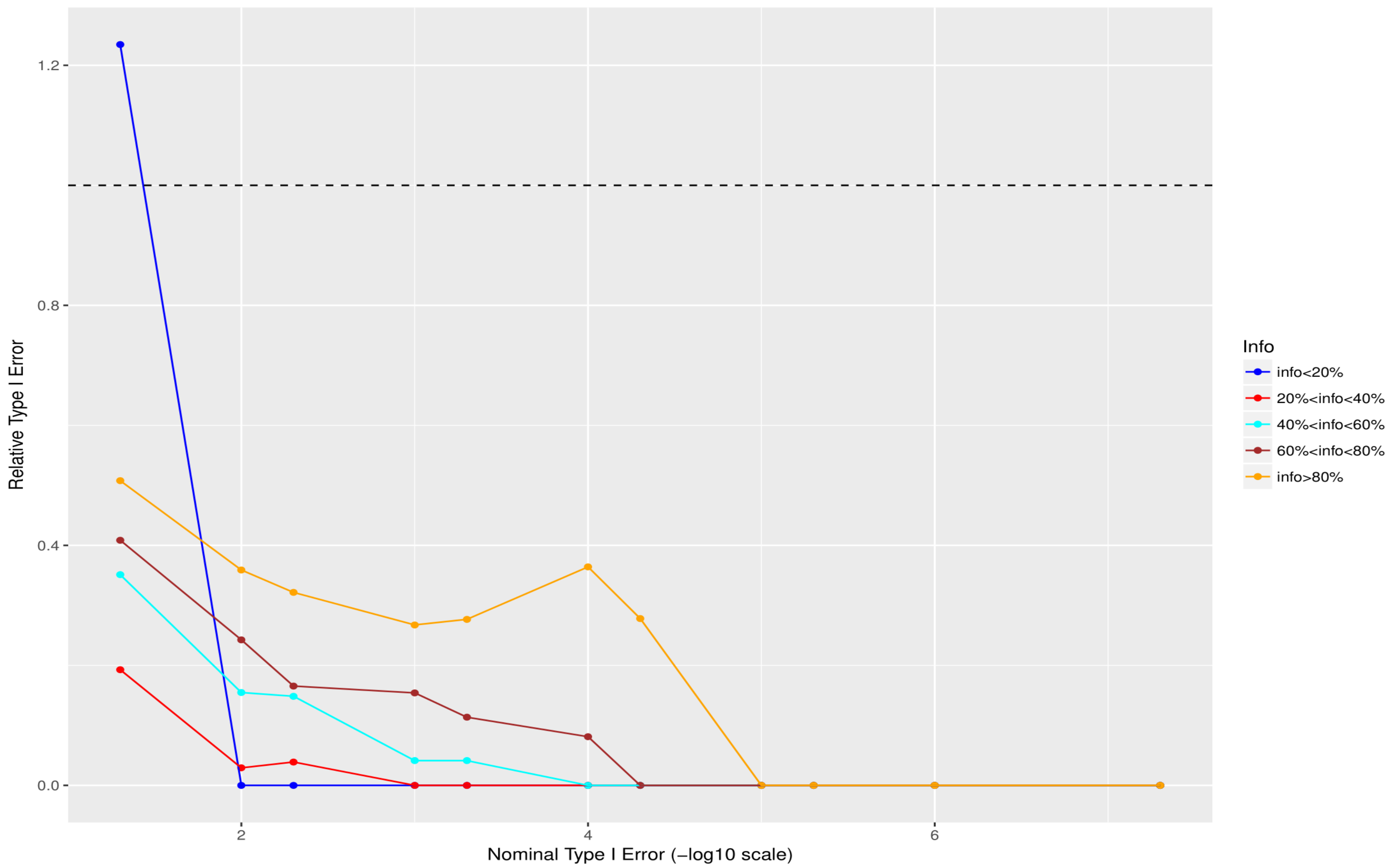


**Fig. S23 Relative size of the test - Cohort 2 for 10%<MAF<50% (the quotient of empirical false positive rate and nominal type I error).**

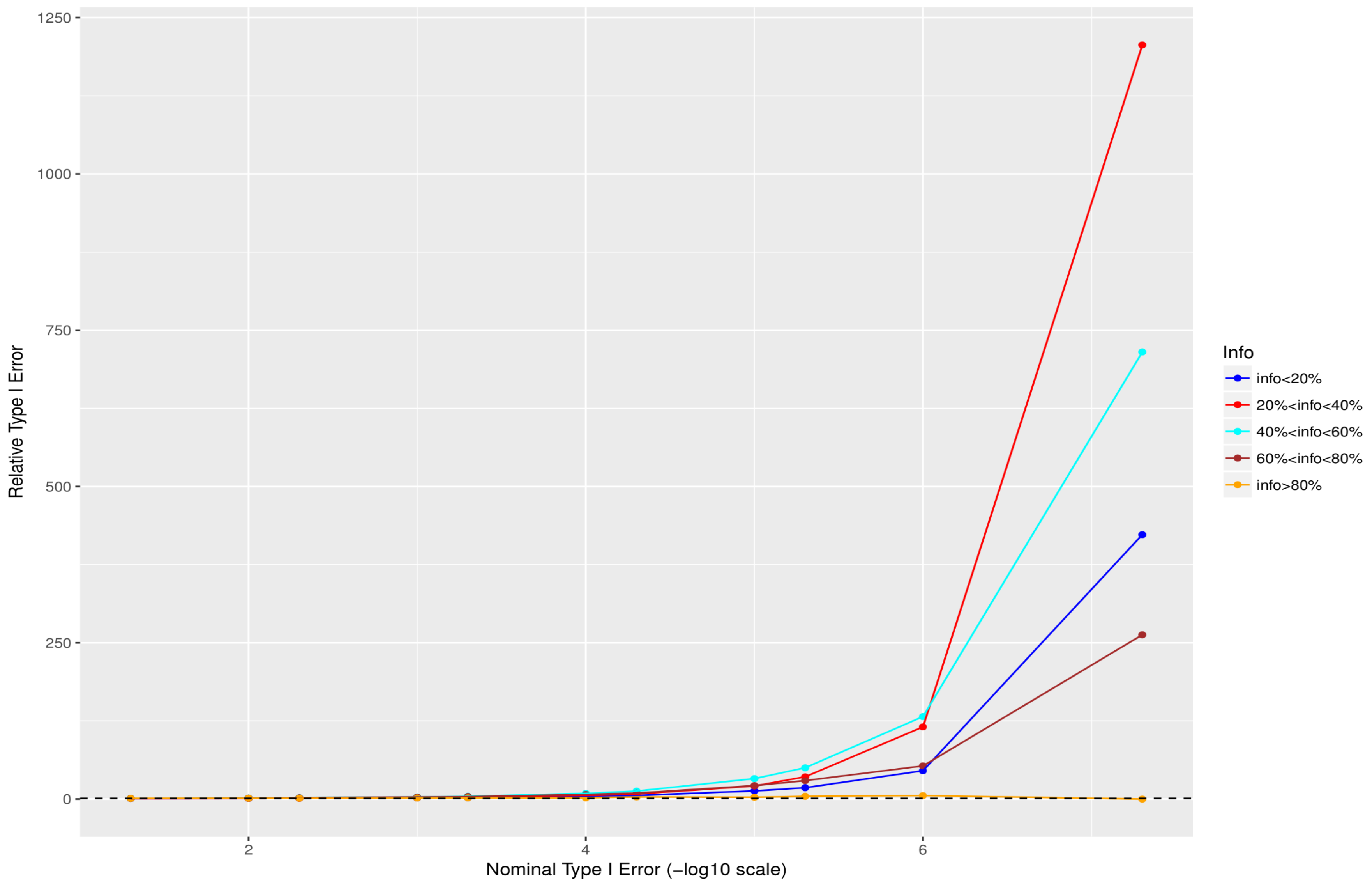


**Fig. S24 Relative size of the test - Cohort 3 for 0.05%<MAF<0.5% (the quotient of empirical false positive rate and nominal type I error).**

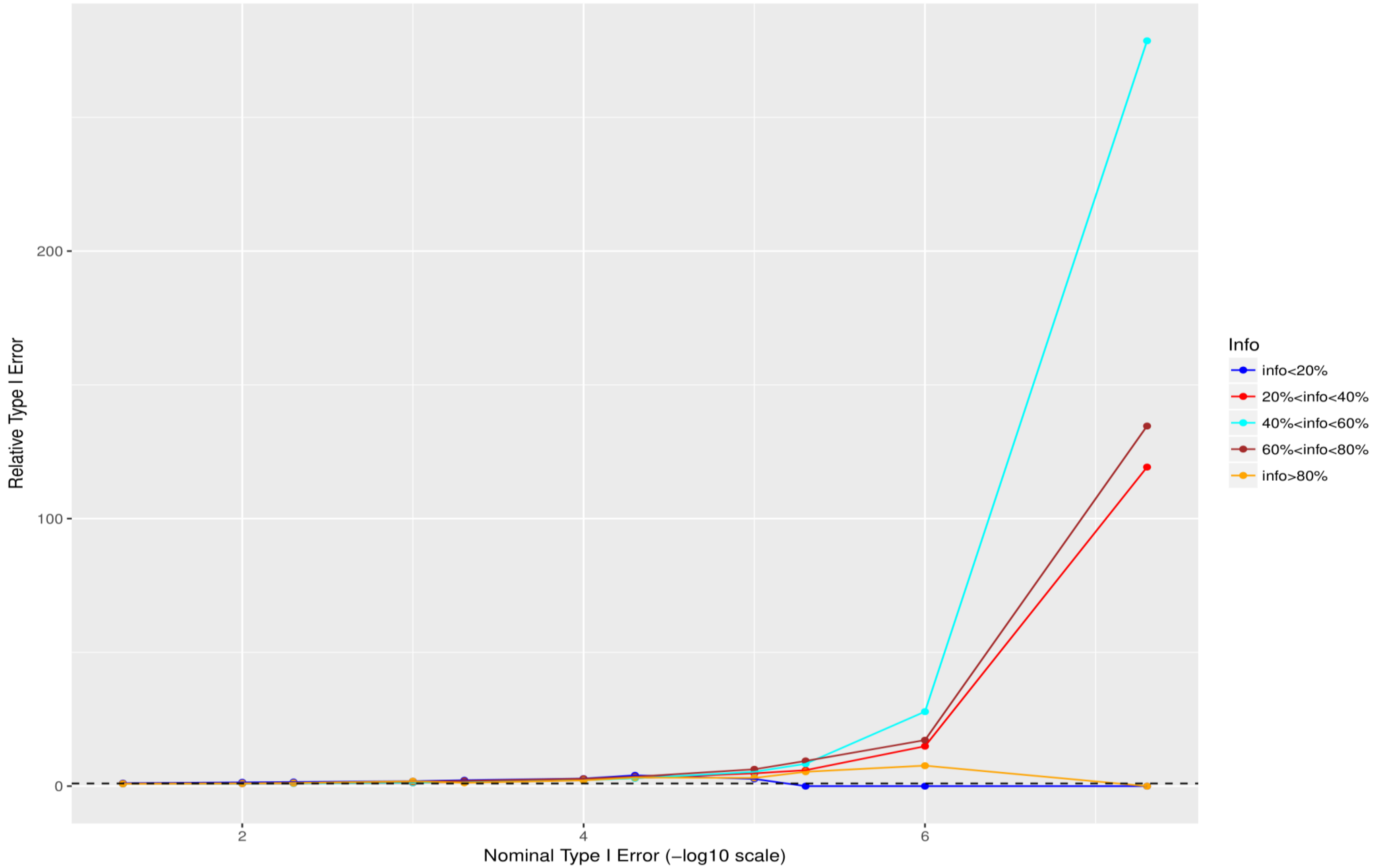


**Fig. S25 Relative size of the test - Cohort 3 for 0. 5%<MAF<1% (the quotient of empirical false positive rate and nominal type I error).**

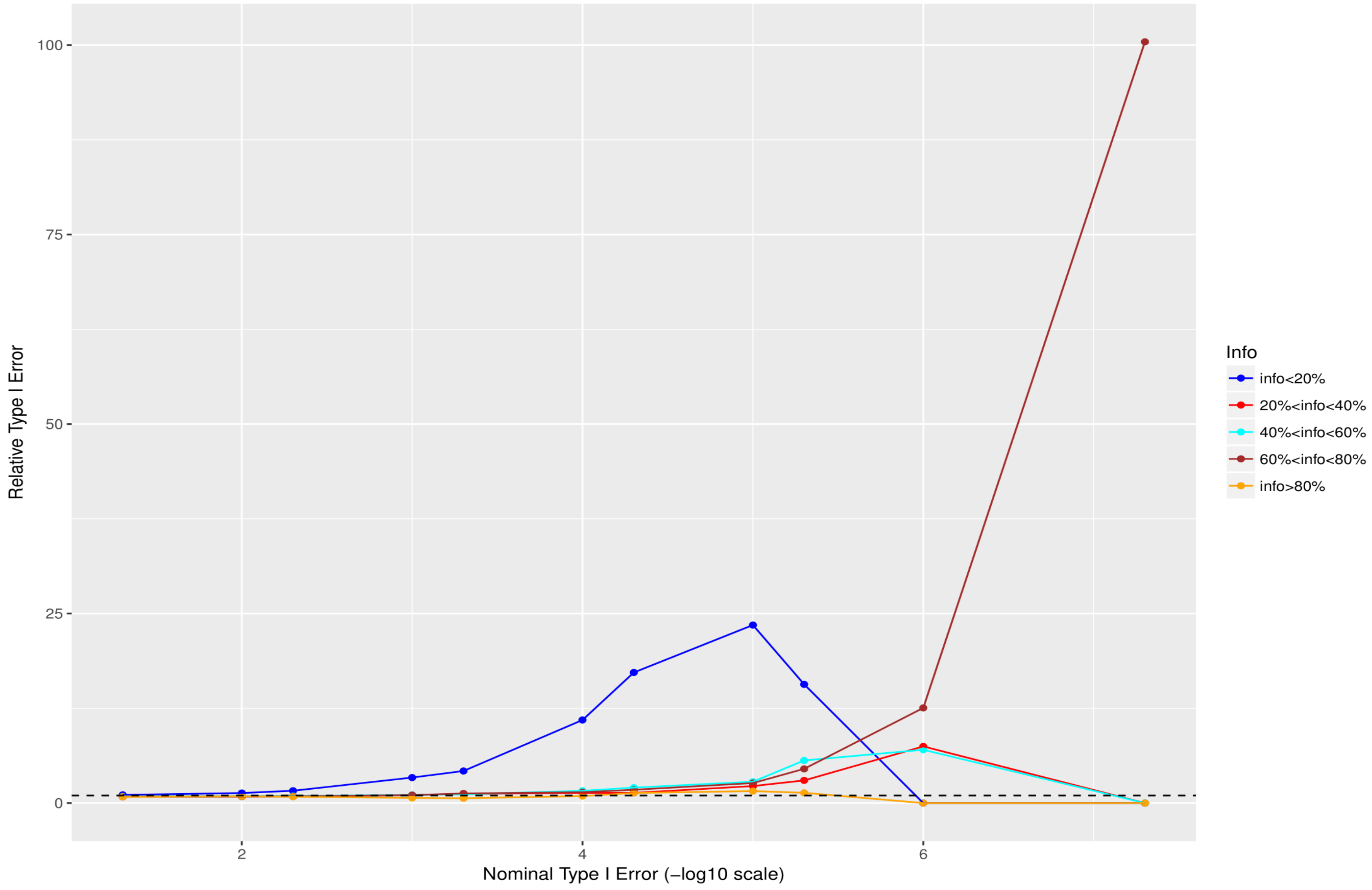


**Fig. S26 Relative size of the test - Cohort 3 for 1%<MAF<2% (the quotient of empirical false positive rate and nominal type I error).**

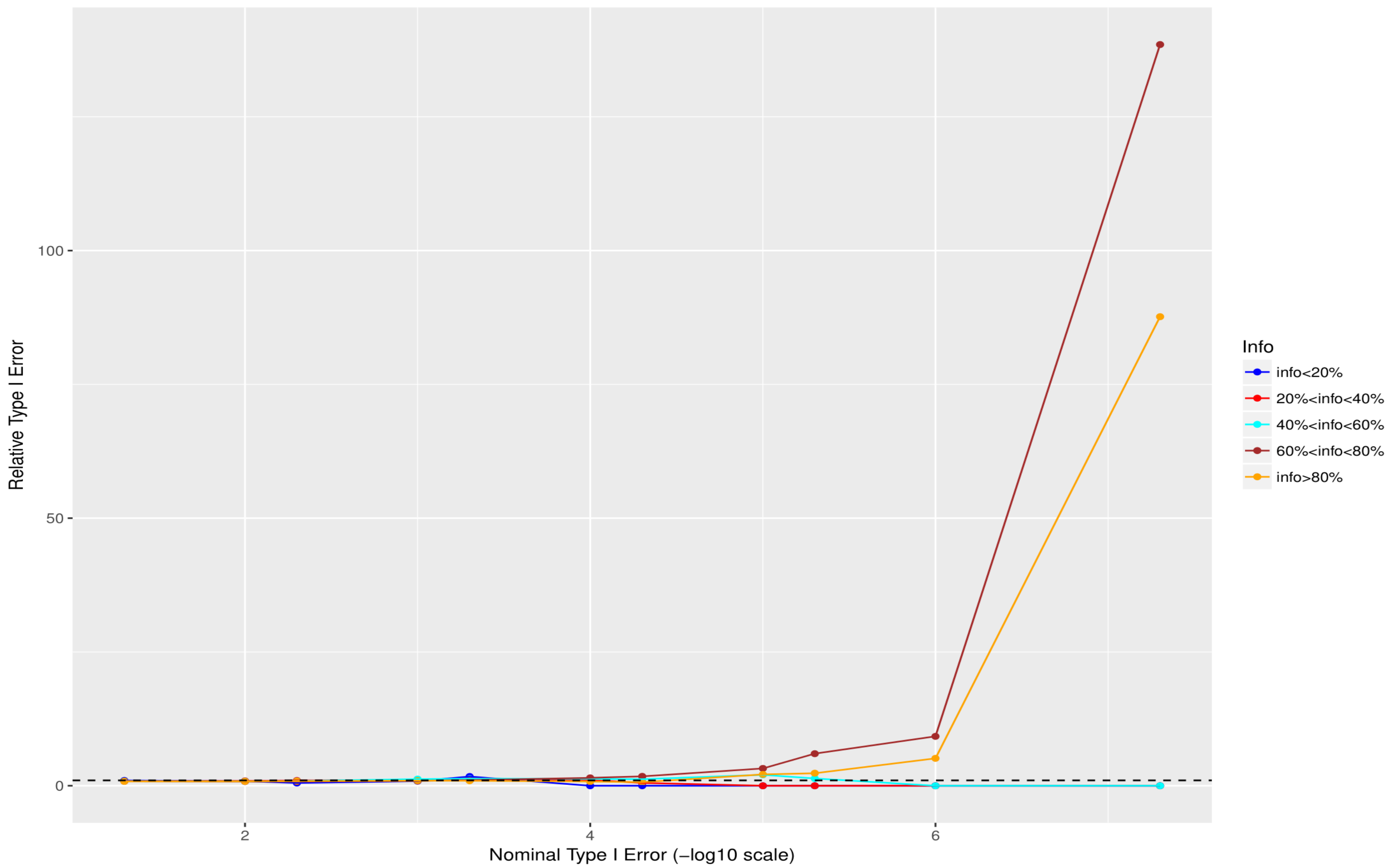


**Fig. S27 Relative size of the test - Cohort 3 for 2%<MAF<5% (the quotient of empirical false positive rate and nominal type I error).**

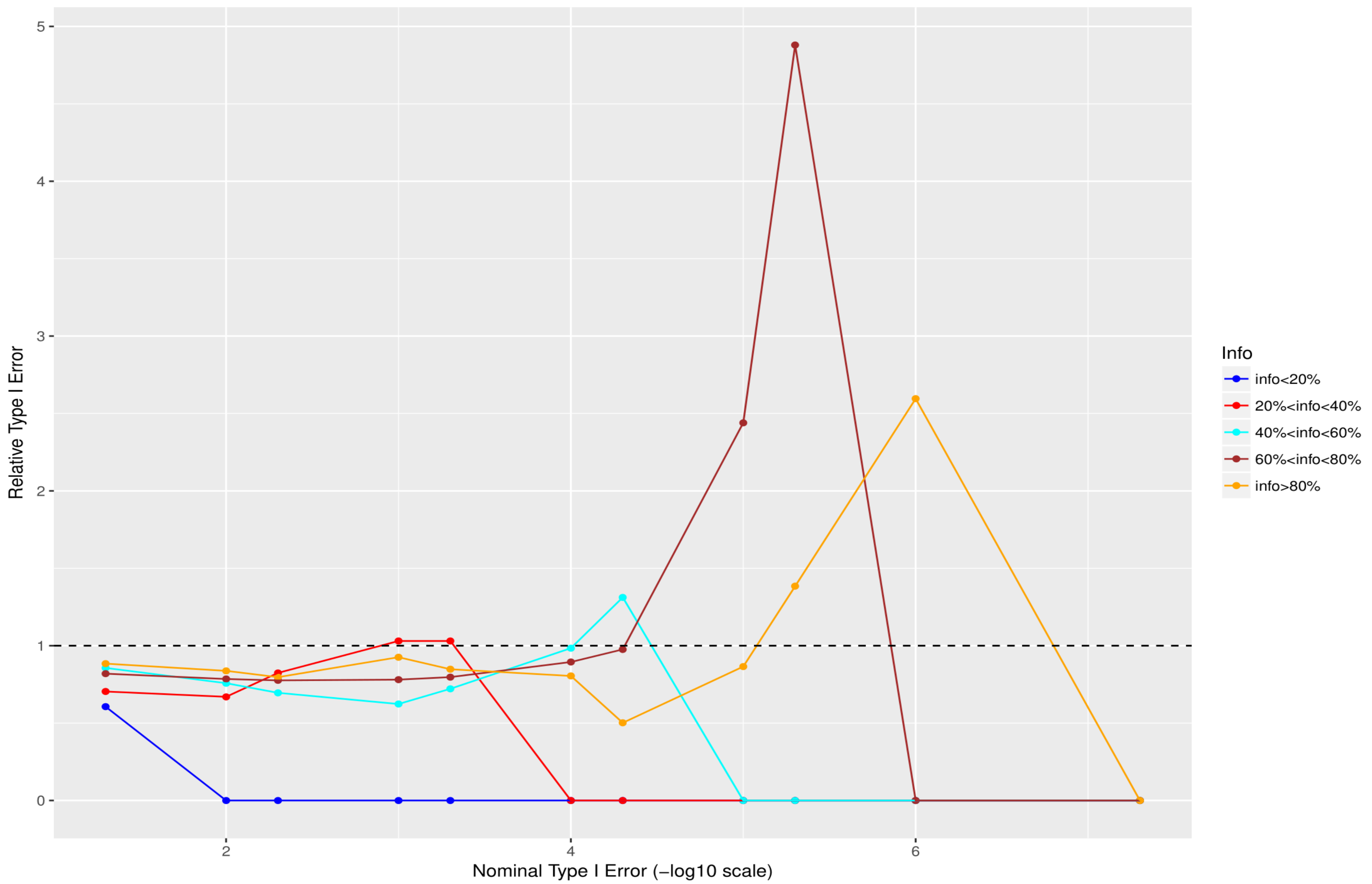


**Fig. S28 Relative size of the test - Cohort 3 for 5%<MAF<10% (the quotient of empirical false positive rate and nominal type I error).**

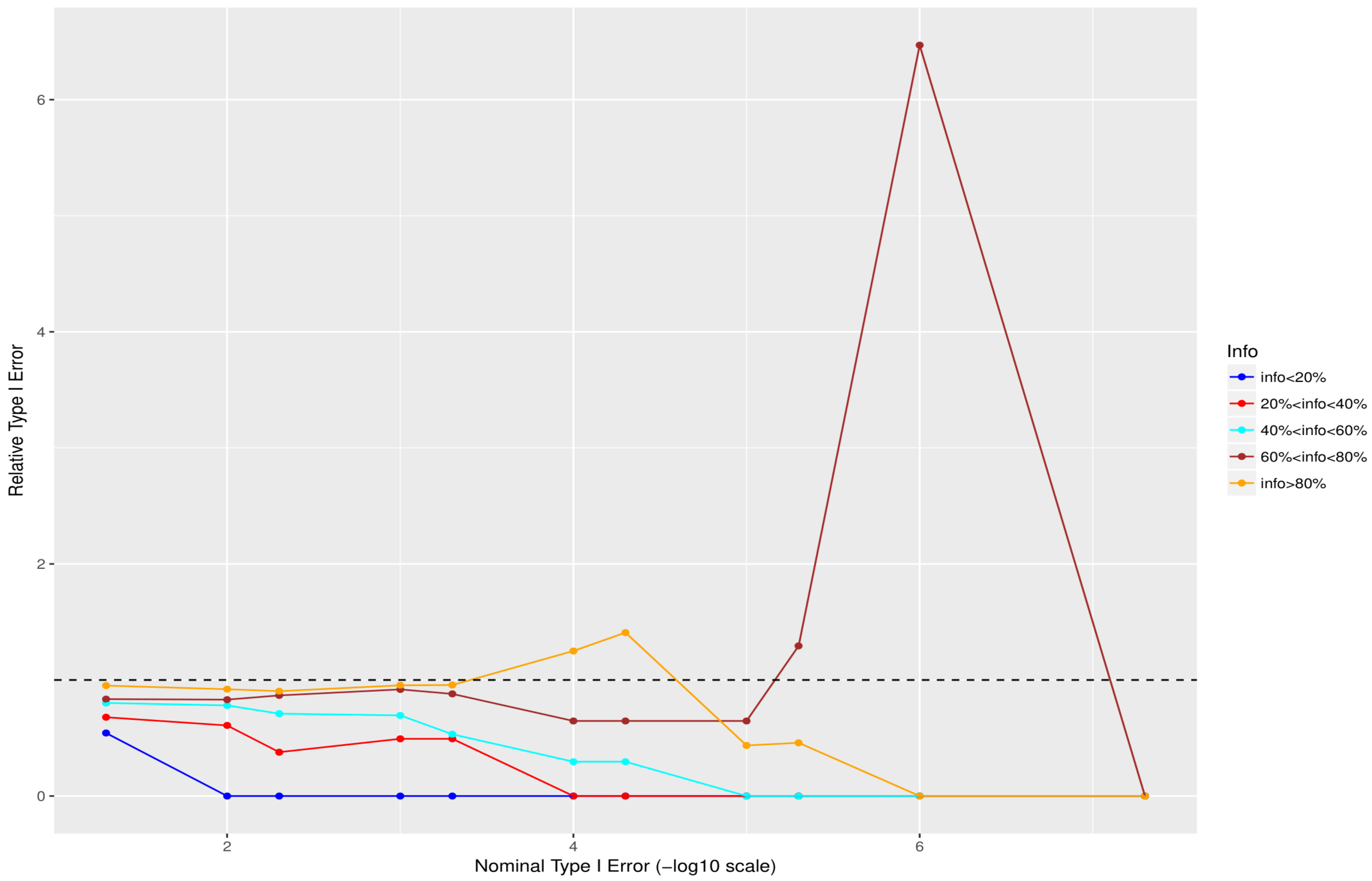


**Fig. S29 Relative size of the test - Cohort 3 for 10%<MAF<50% (the quotient of empirical false positive rate and nominal type I error).**

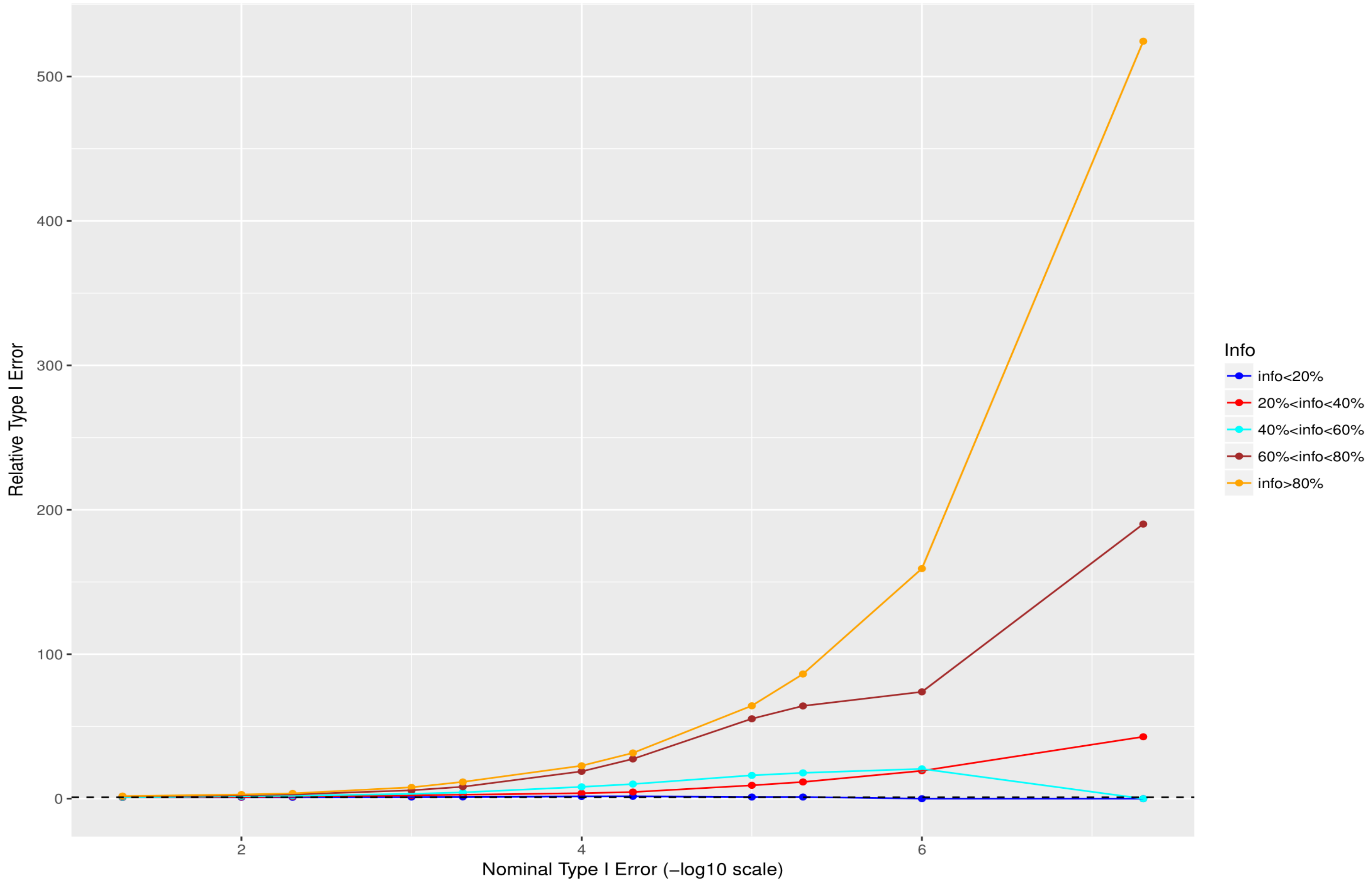


**Fig. S30 Relative size of the test - Cohort 4 for 0.05%<MAF<0.5% (the quotient of empirical false positive rate and nominal type I error).**

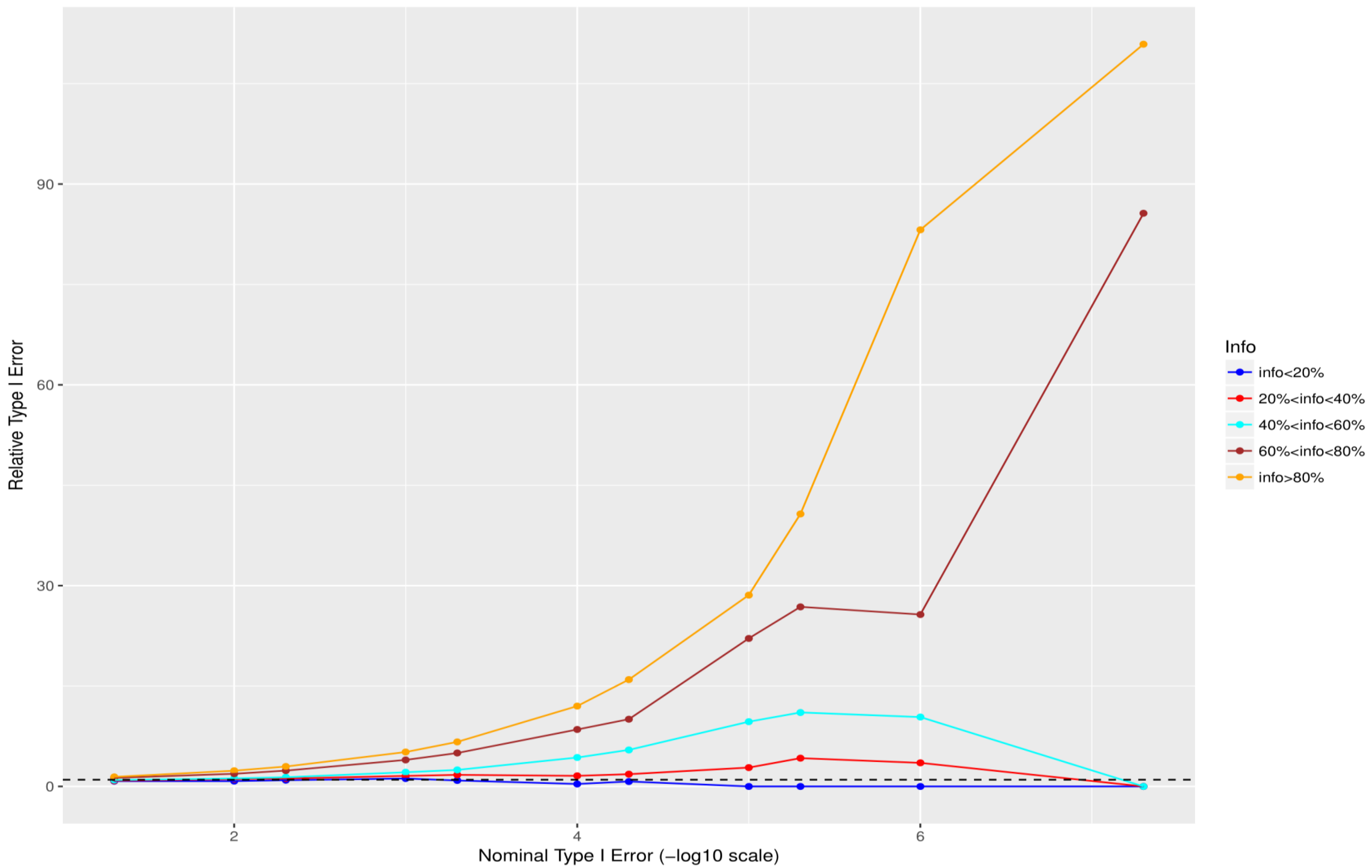


**Fig. S31 Relative size of the test - Cohort 4 for 0. 5%<MAF<1% (the quotient of empirical false positive rate and nominal type I error).**

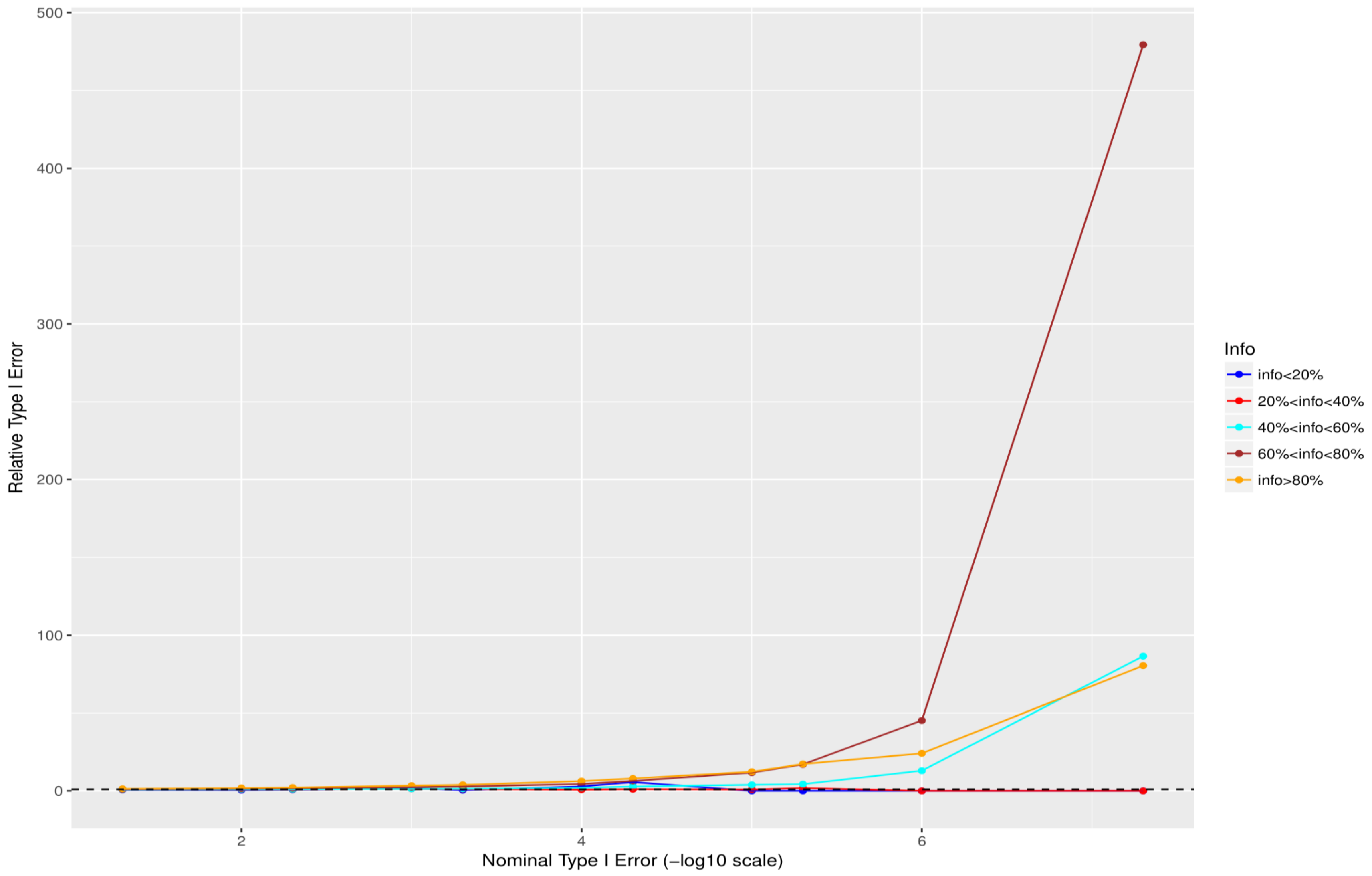


**Fig. S32 Relative size of the test - Cohort 4 for 1%<MAF<2% (the quotient of empirical false positive rate and nominal type I error).**

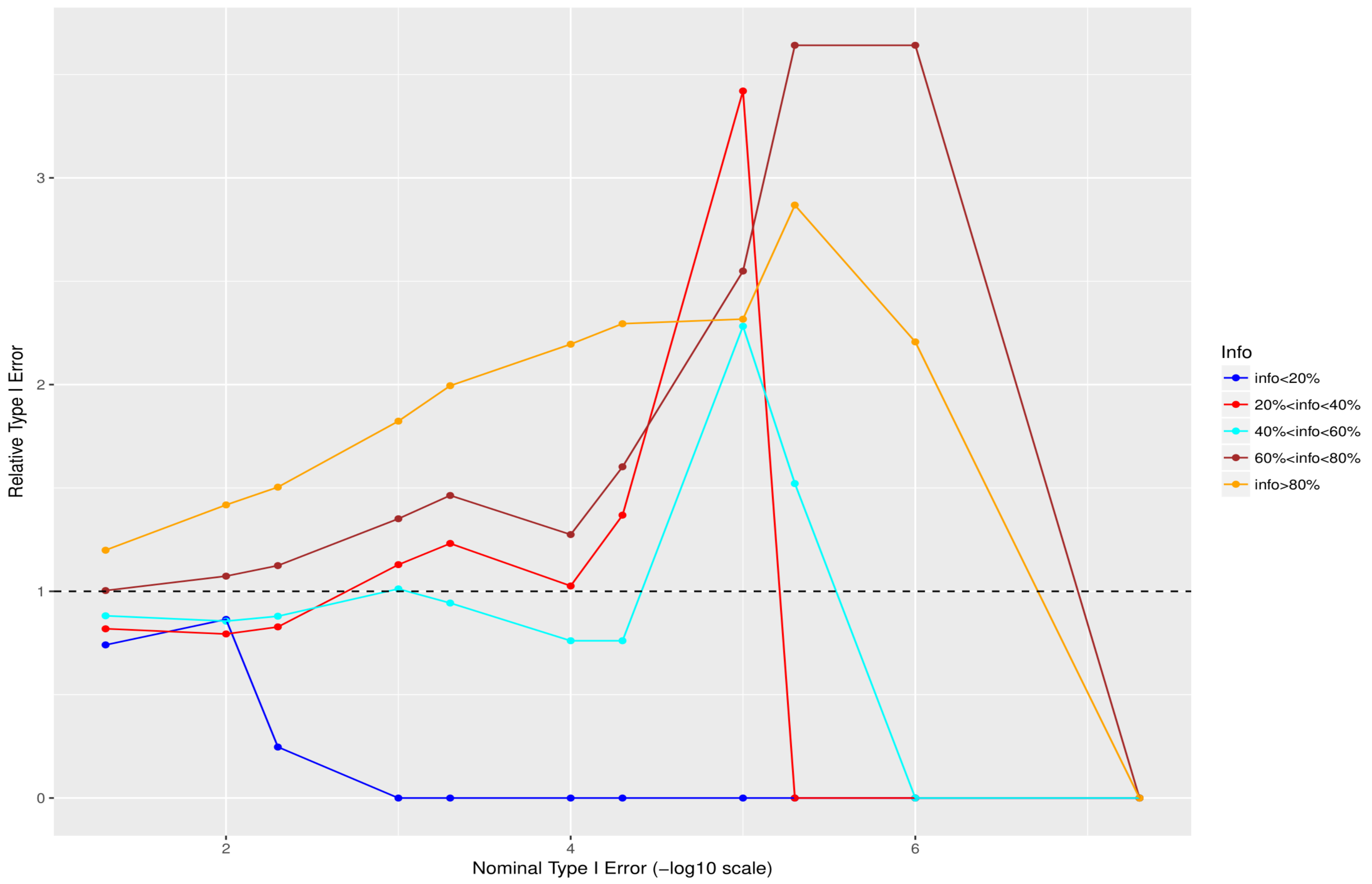


**Fig. S33 Relative size of the test - Cohort 4 for 2%<MAF<5% (the quotient of empirical false positive rate and nominal type I error).**

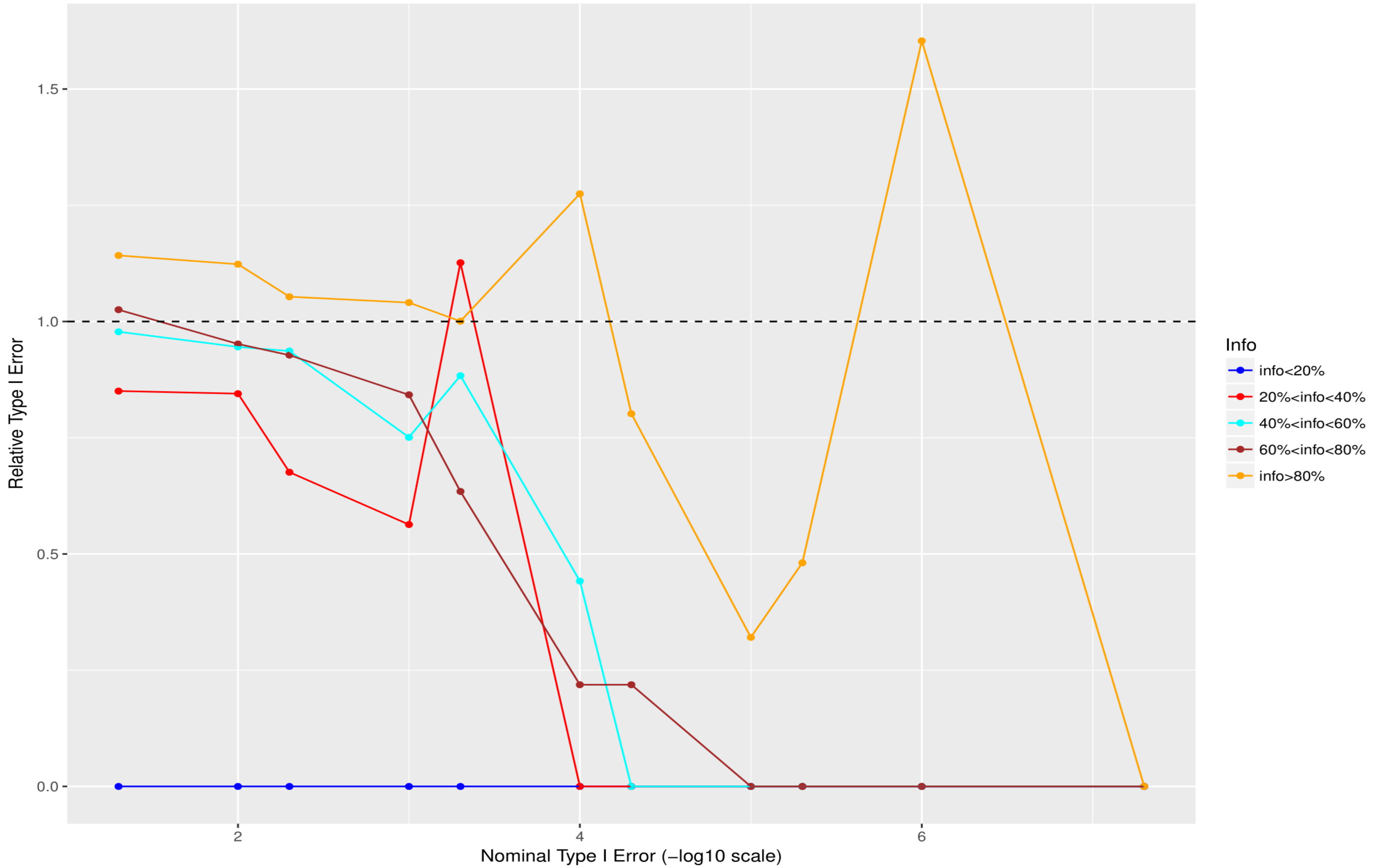


**Fig. S34 Relative size of the test - Cohort 4 for 5%<MAF<10% (the quotient of empirical false positive rate and nominal type I error).**

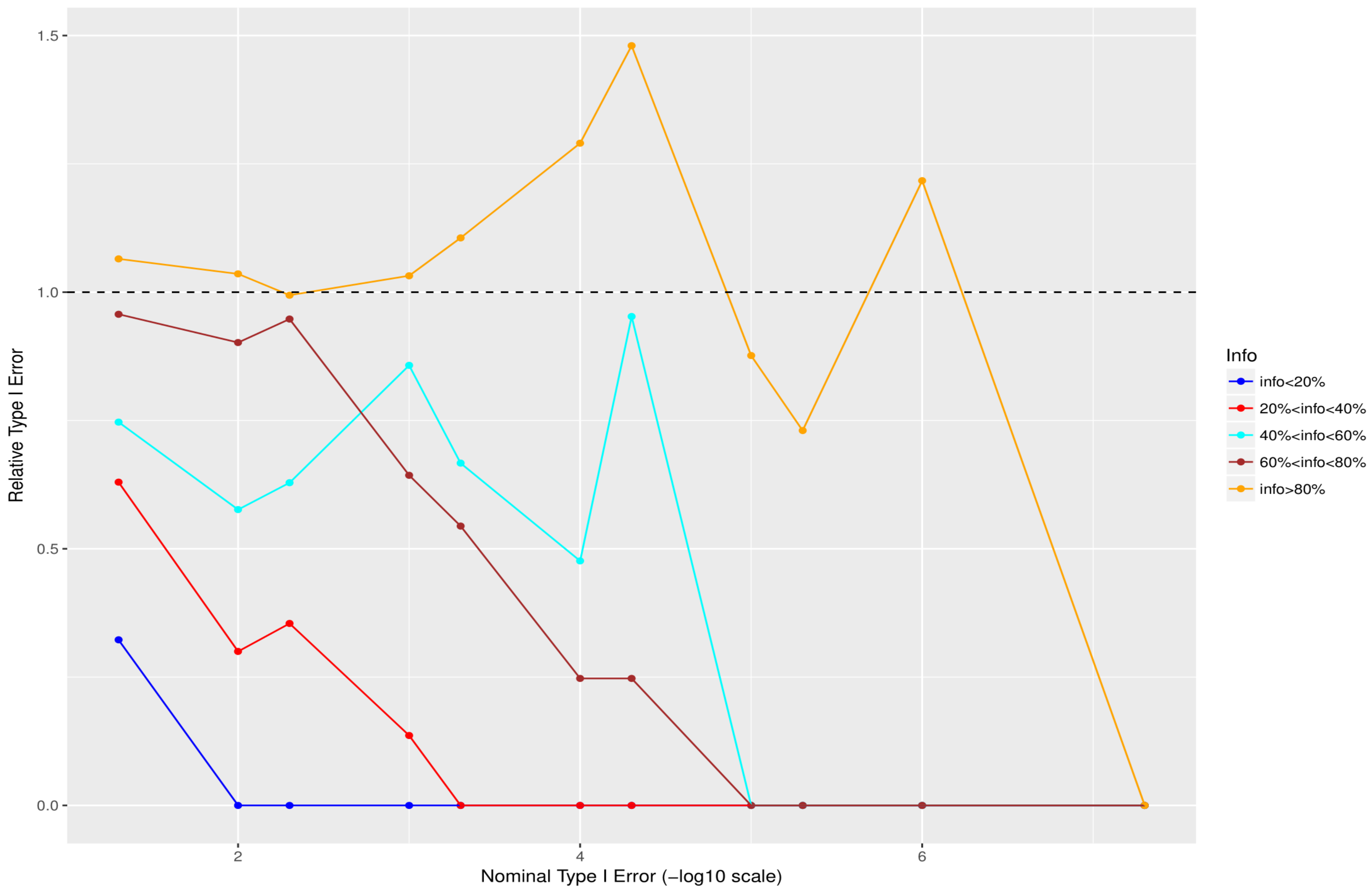


**Fig. S35 Relative size of the test - Cohort 4 for 10%<MAF<50% (the quotient of empirical false positive rate and nominal type I error).**

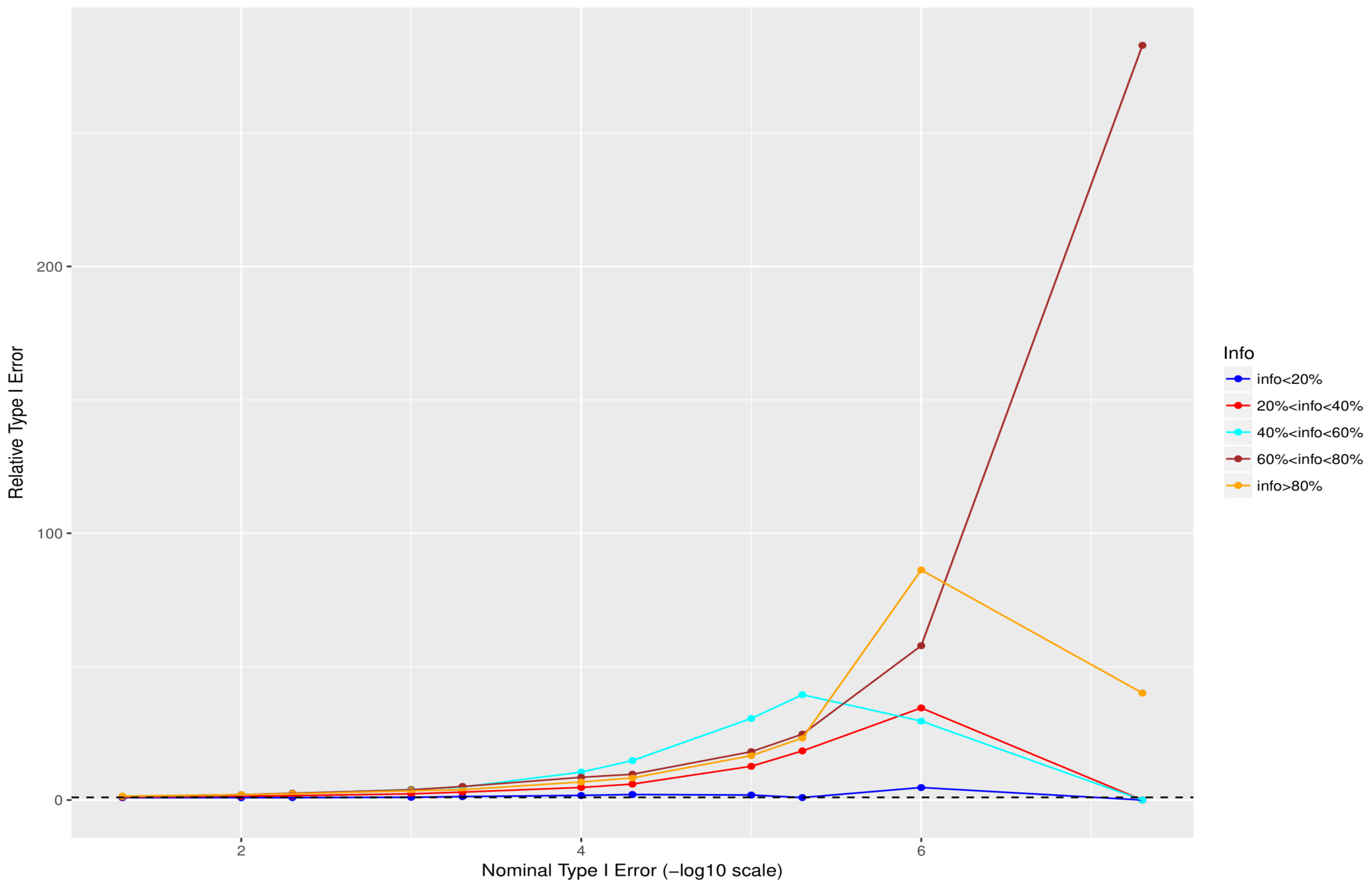


**Fig. S36 Relative size of the test - Cohort 5 for 0.05%<MAF<0.5% (the quotient of empirical false positive rate and nominal type I error).**

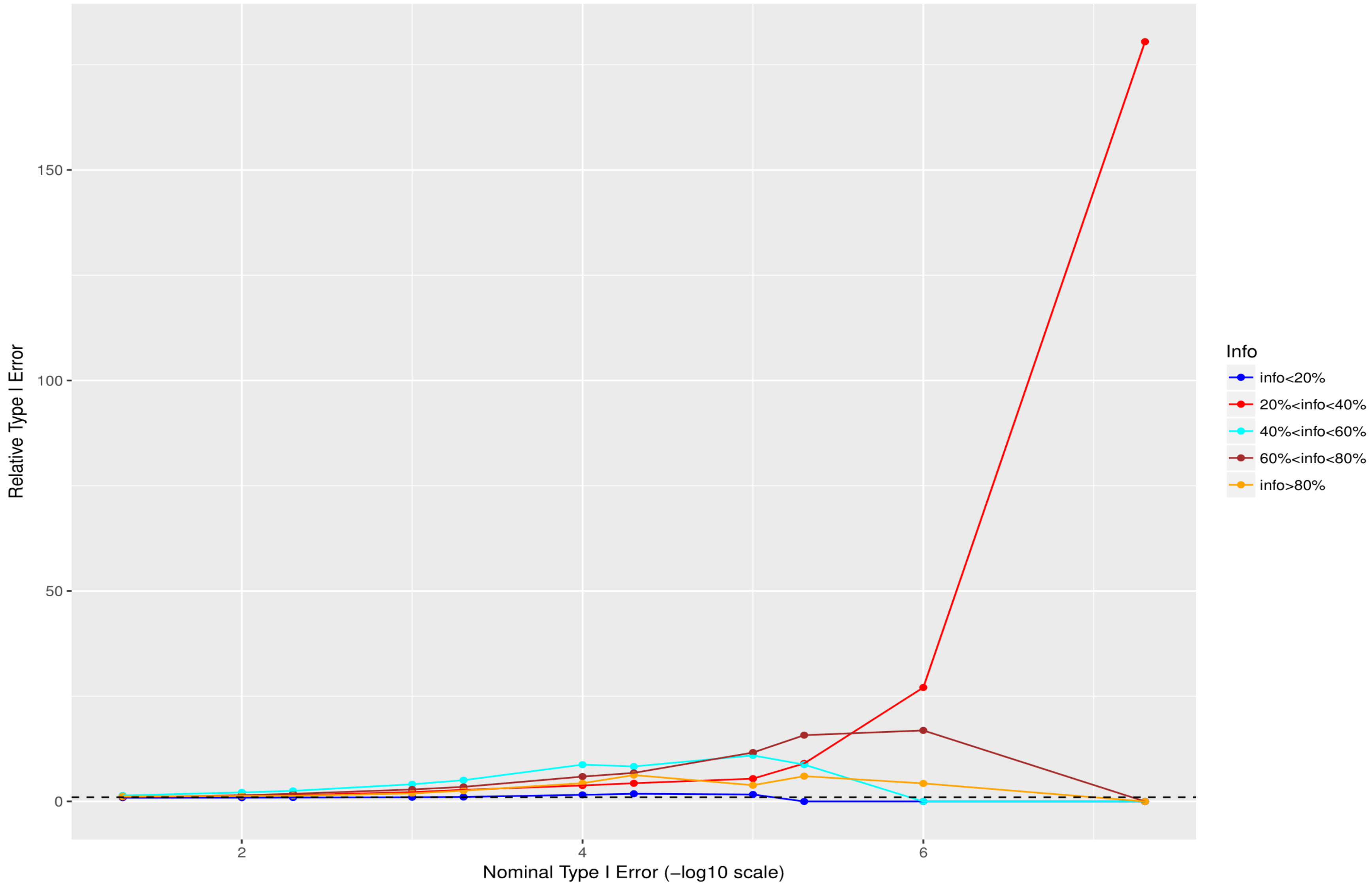


**Fig. S37 Relative size of the test - Cohort 5 for 0. 5%<MAF<1% (the quotient of empirical false positive rate and nominal type I error).**

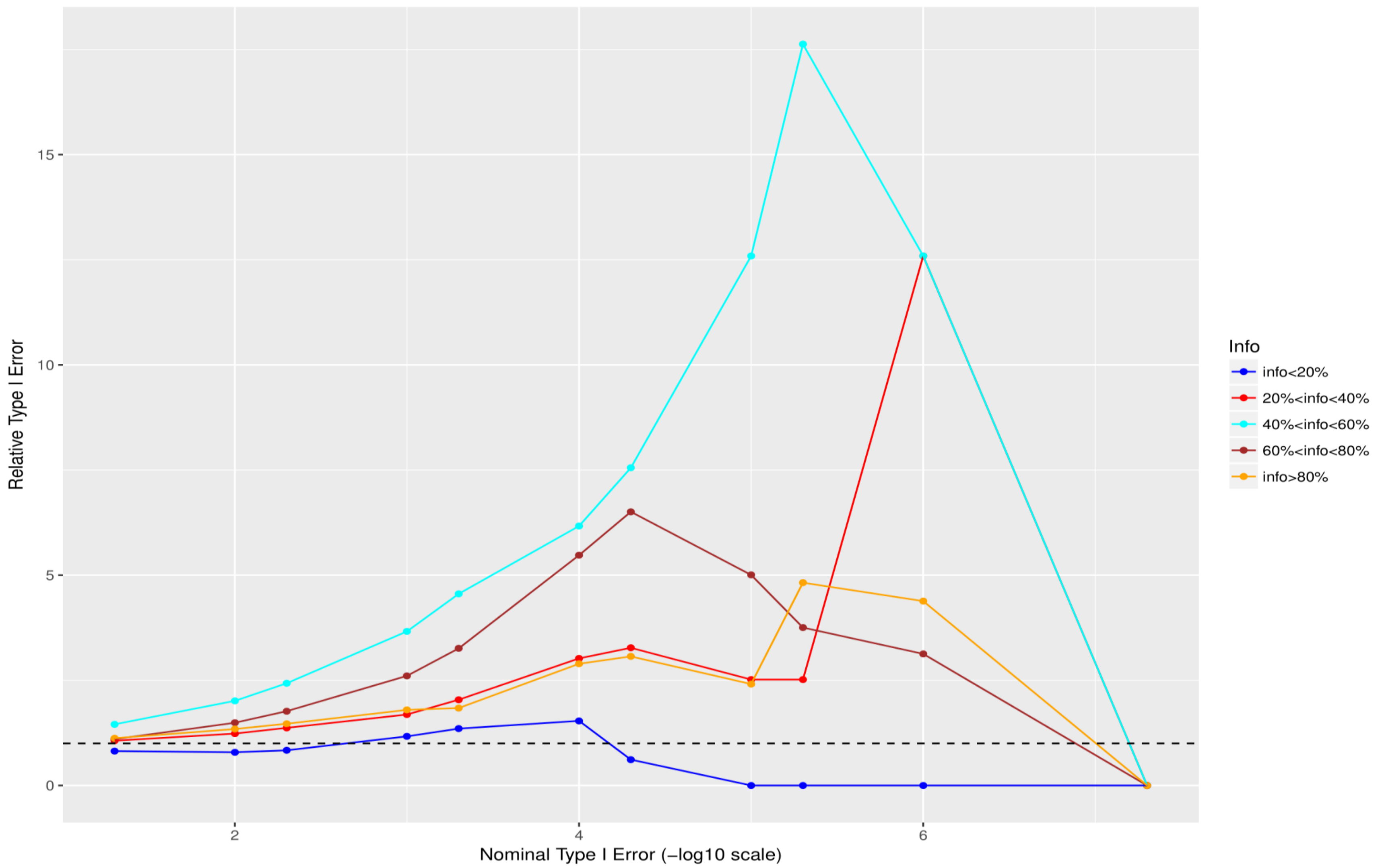


**Fig. S38 Relative size of the test - Cohort 5 for 1%<MAF<2% (the quotient of empirical false positive rate and nominal type I error).**

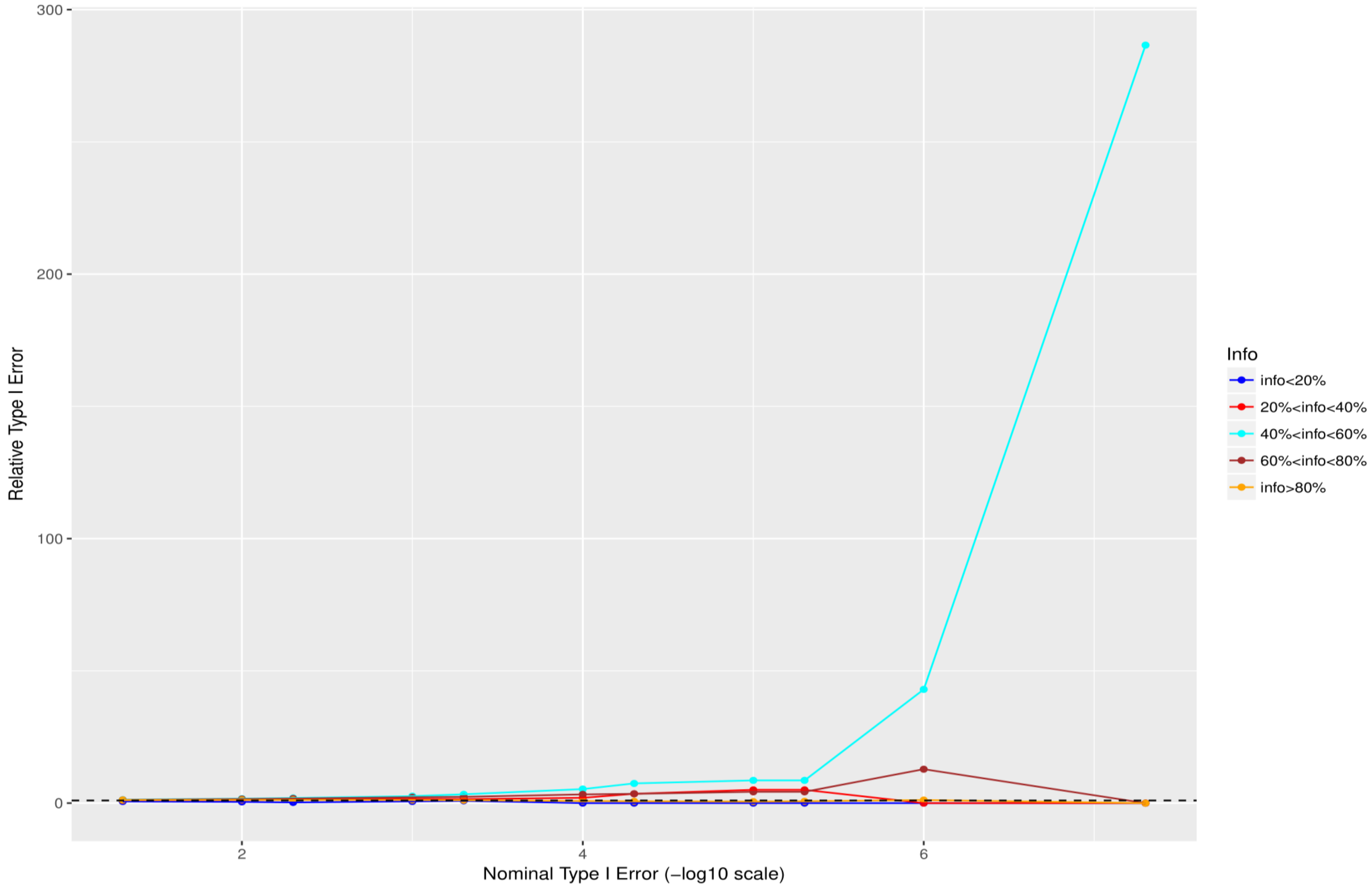


**Fig. S39 Relative size of the test - Cohort 5 for 2%<MAF<5% (the quotient of empirical false positive rate and nominal type I error).**

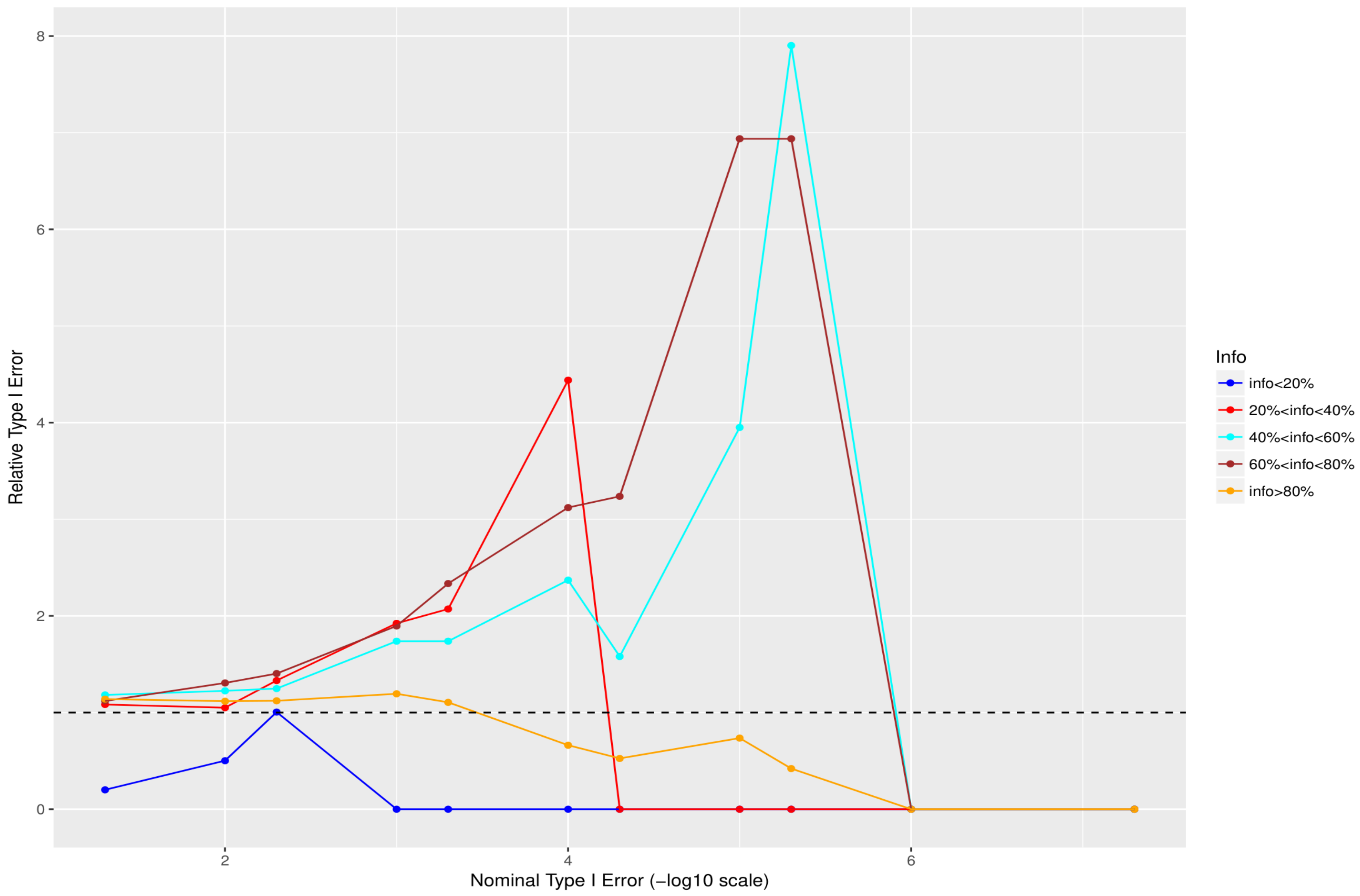


**Fig. S40 Relative size of the test - Cohort 5 for 5%<MAF<10% (the quotient of empirical false positive rate and nominal type I error).**

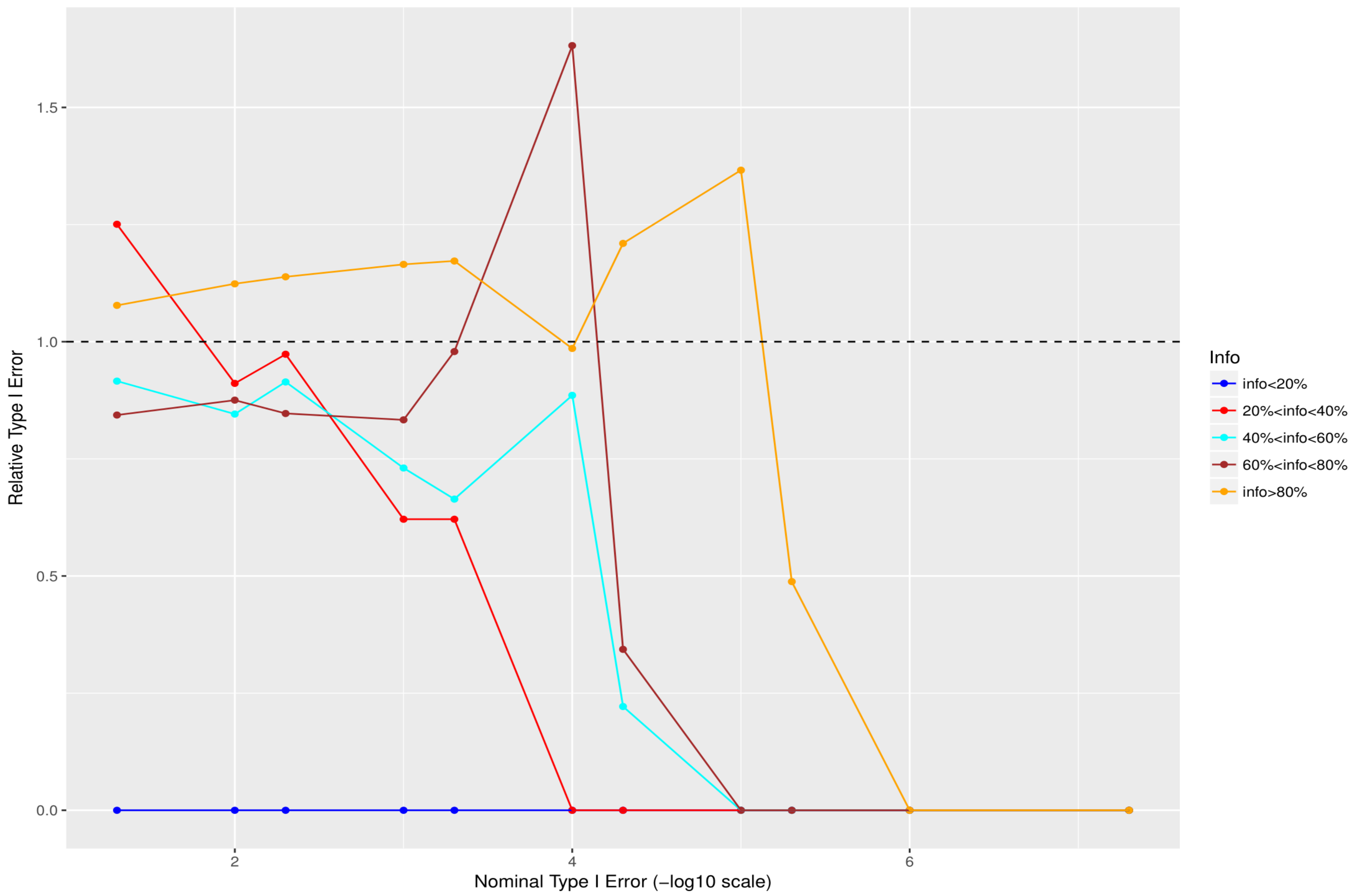


**Fig. S41 Relative size of the test - Cohort 5 for 10%<MAF<50% (the quotient of empirical false positive rate and nominal type I error).**

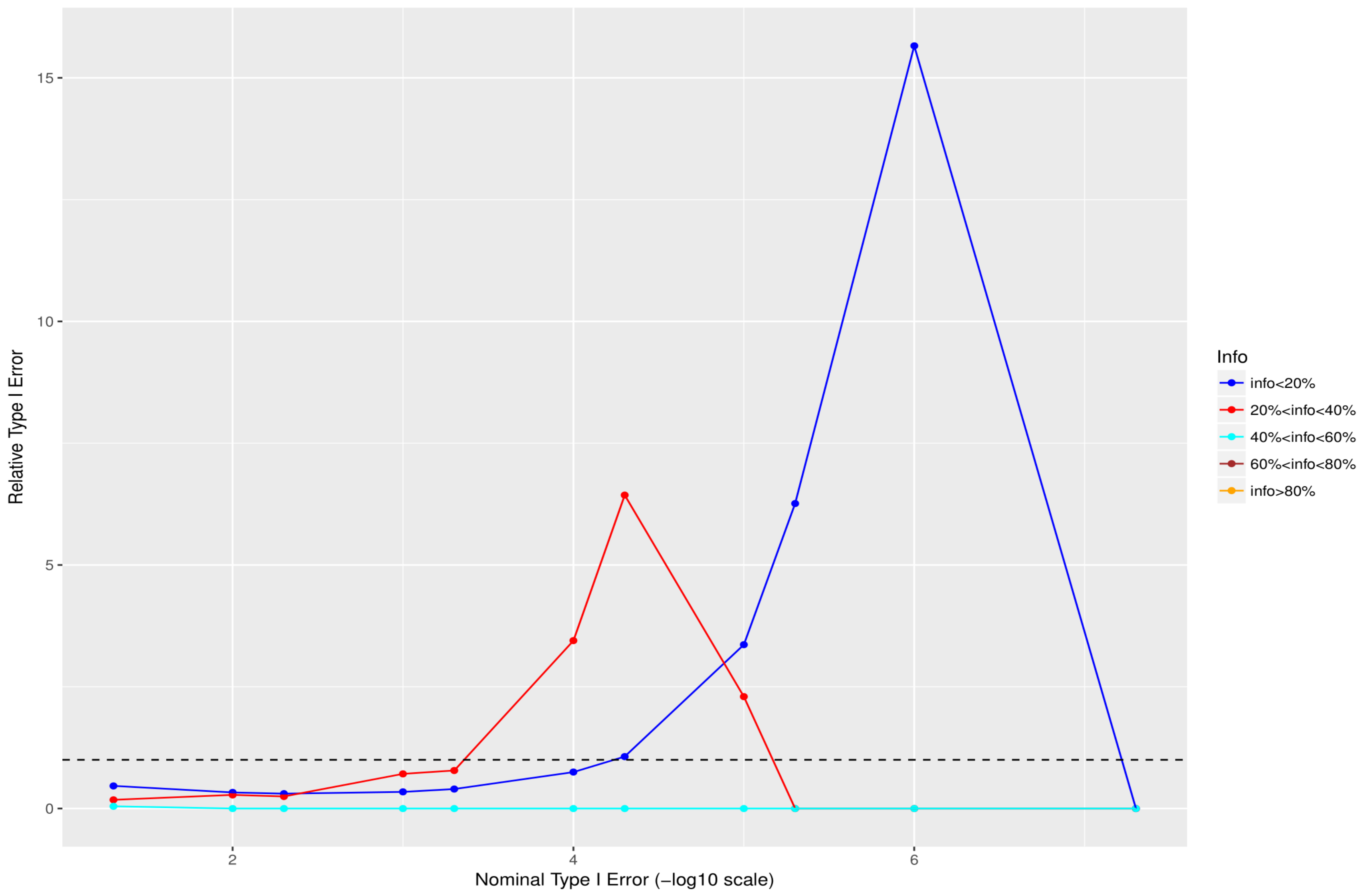


**Fig. S42 Relative size of the test - nullified data for MAF<0.05% (the quotient of empirical false positive rate and nominal type I error).**

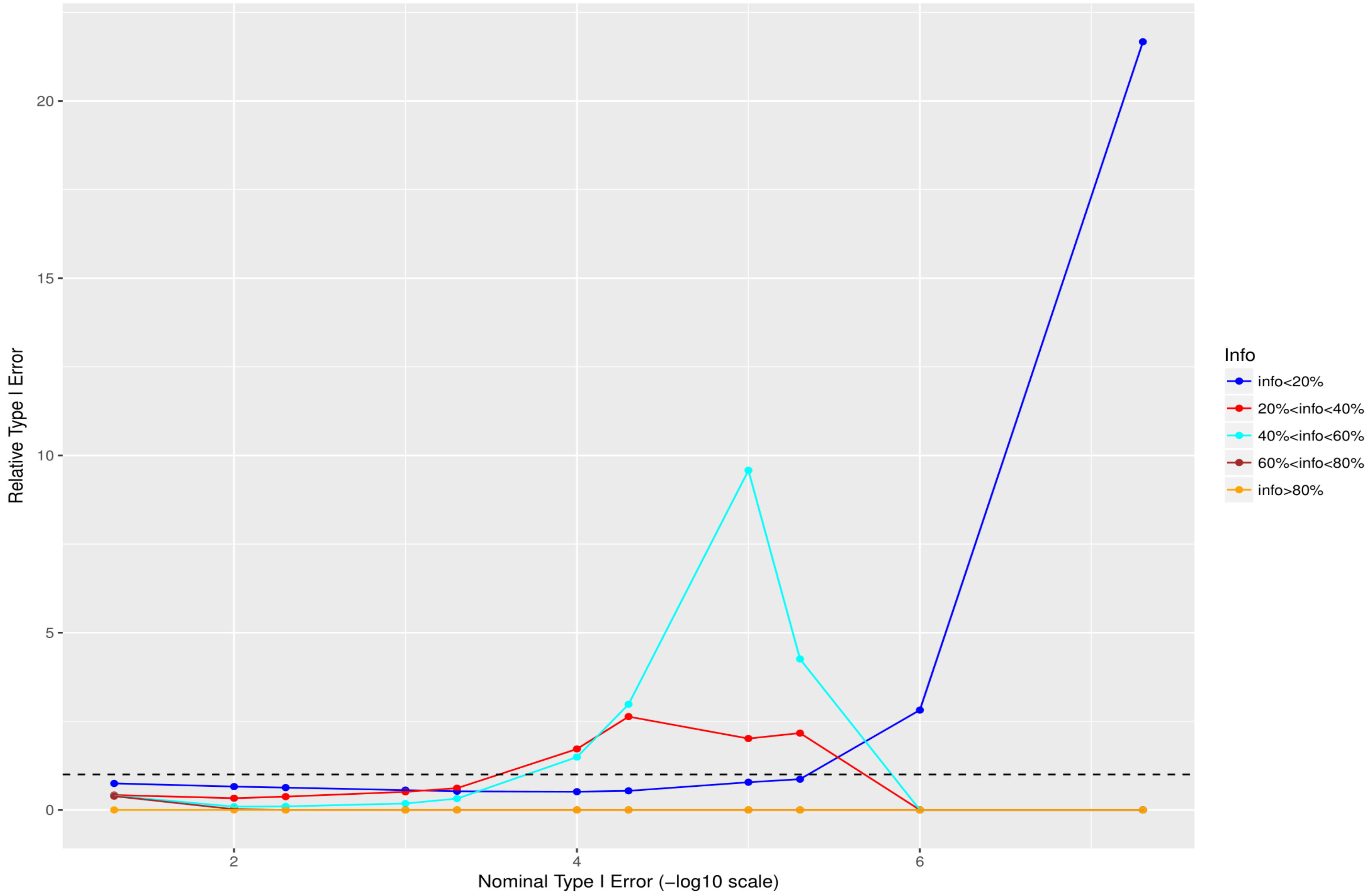


**Fig. S43 Relative size of the test - nullified data for 0.05%<MAF<0.5% (the quotient of empirical false positive rate and nominal type I error).**

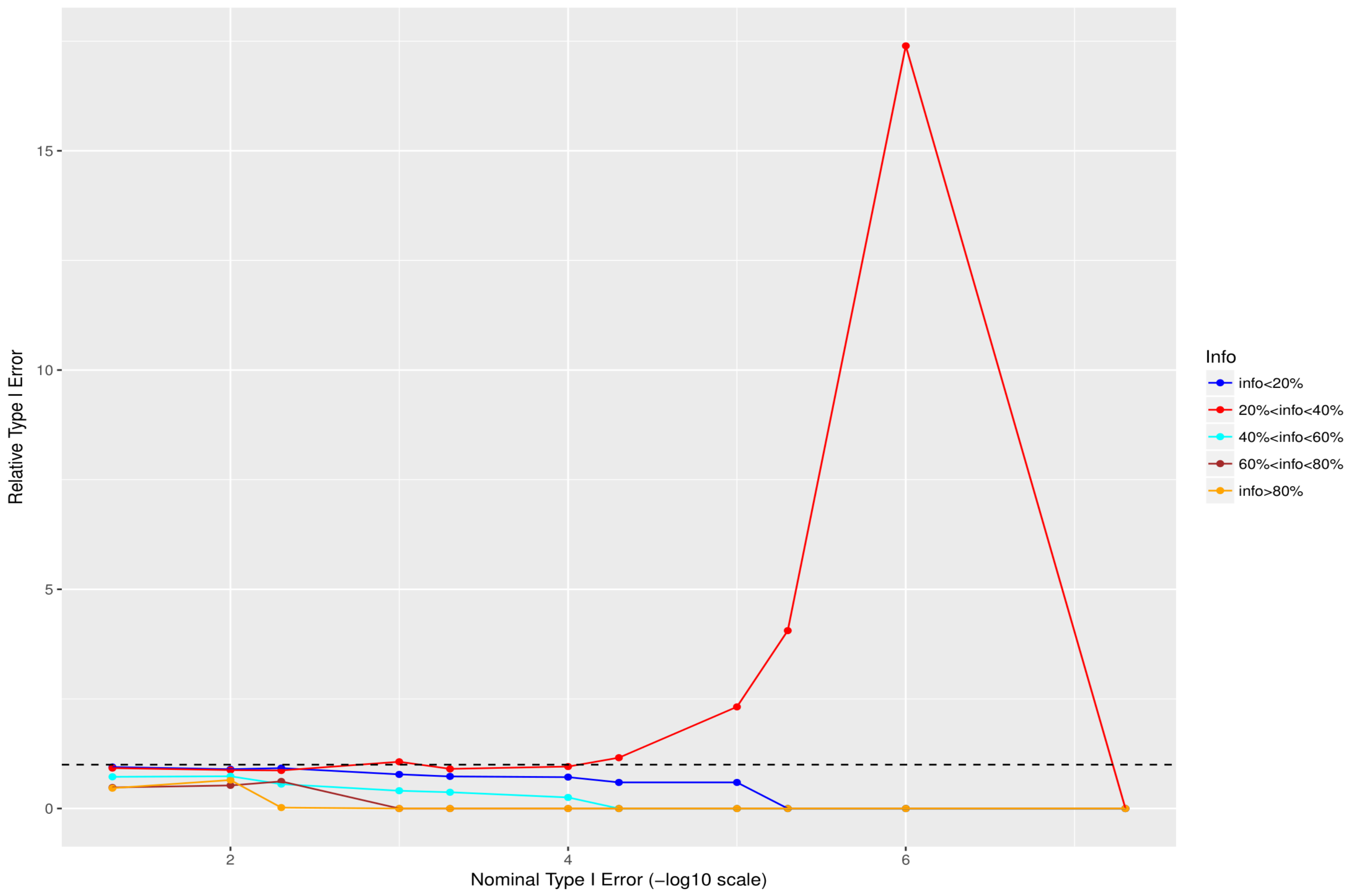


**Fig. S44 Relative size of the test - nullified data for 0.5%<MAF<1% (the quotient of empirical false positive rate and nominal type I error).**

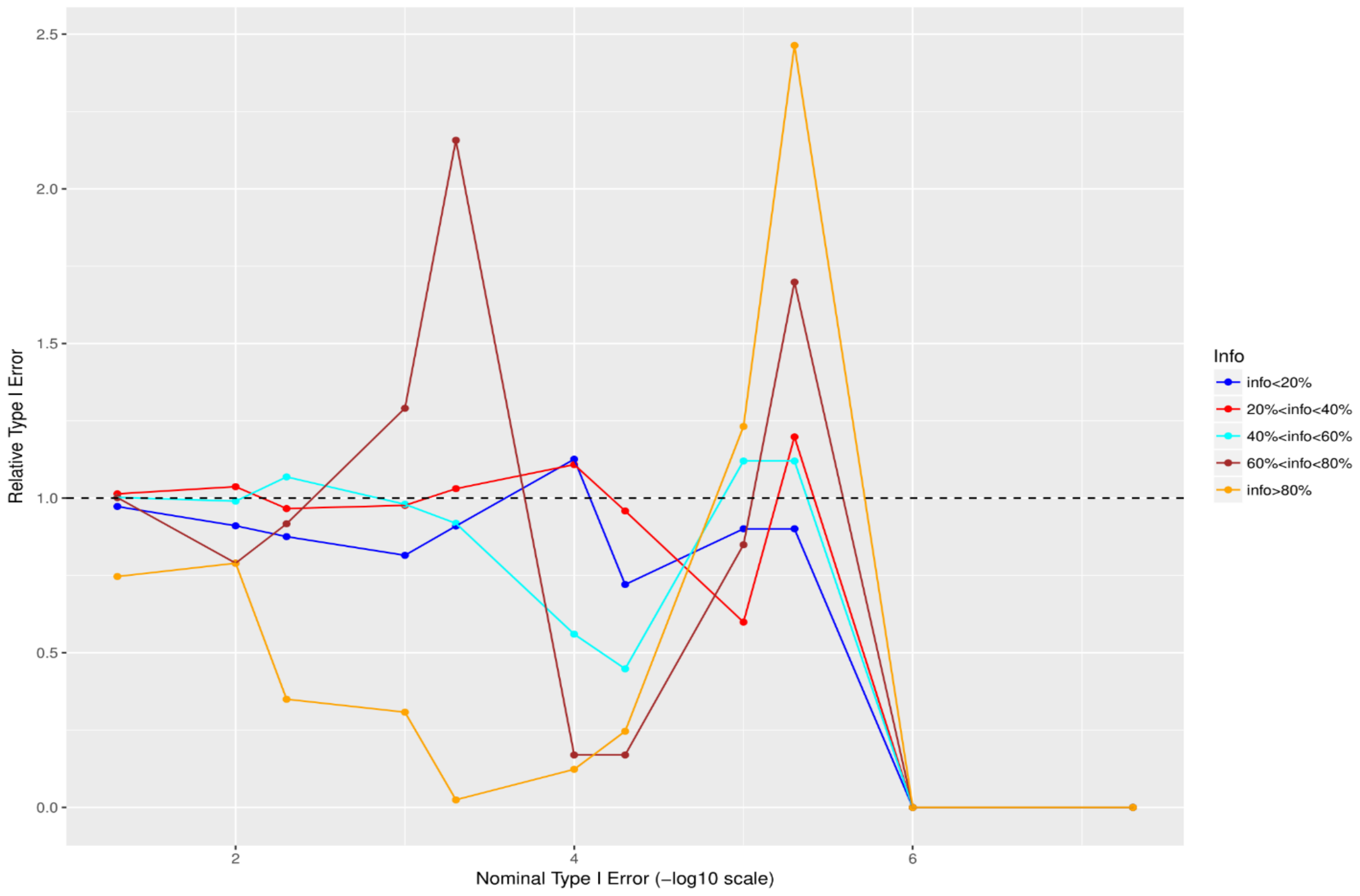


**Fig. S45 Relative size of the test - nullified data for 1%<MAF<2% (the quotient of empirical false positive rate and nominal type I error).**

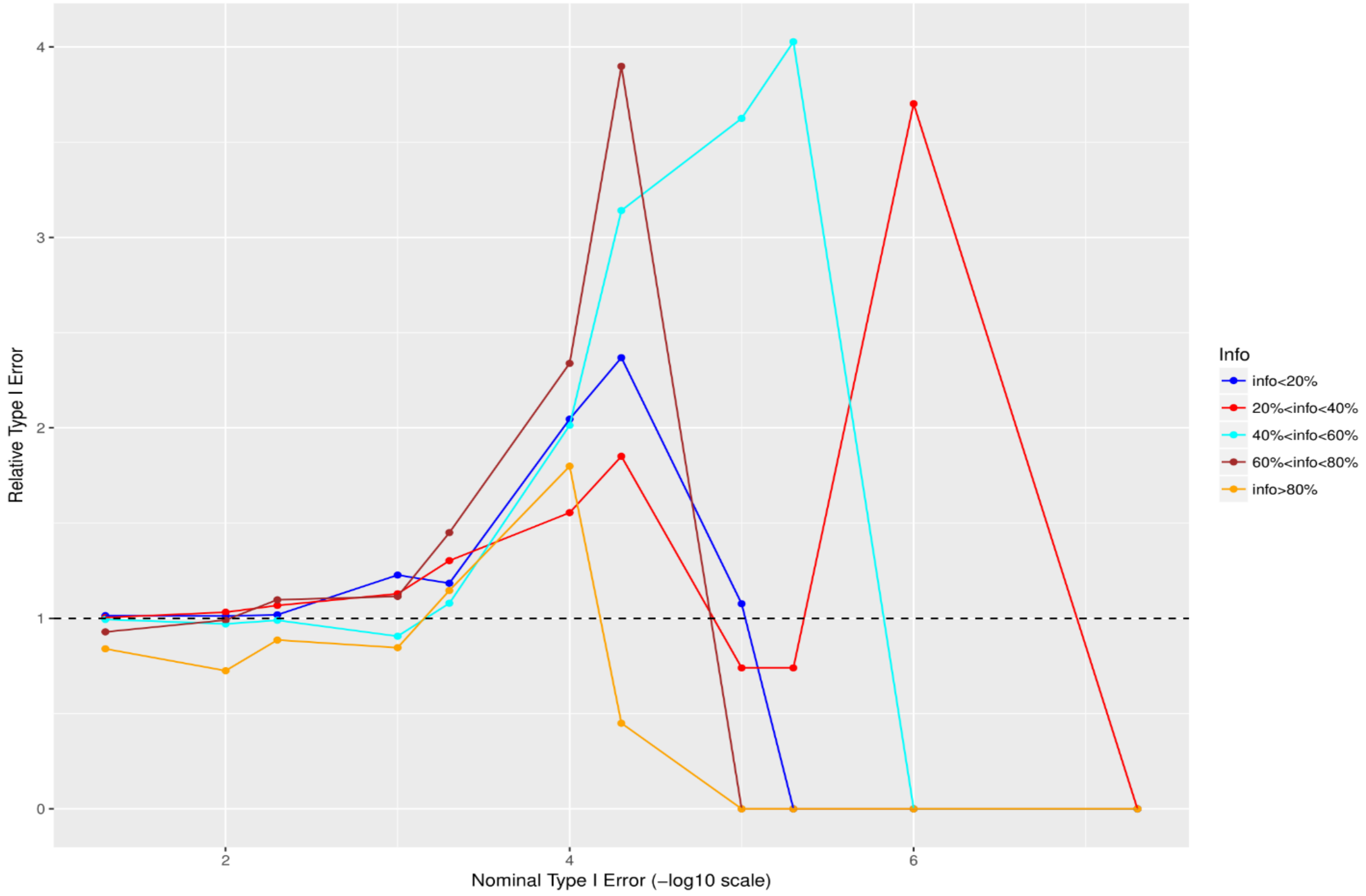


**Fig. S46 Relative size of the test - nullified data for 2%<MAF<5% (the quotient of empirical false positive rate and nominal type I error).**

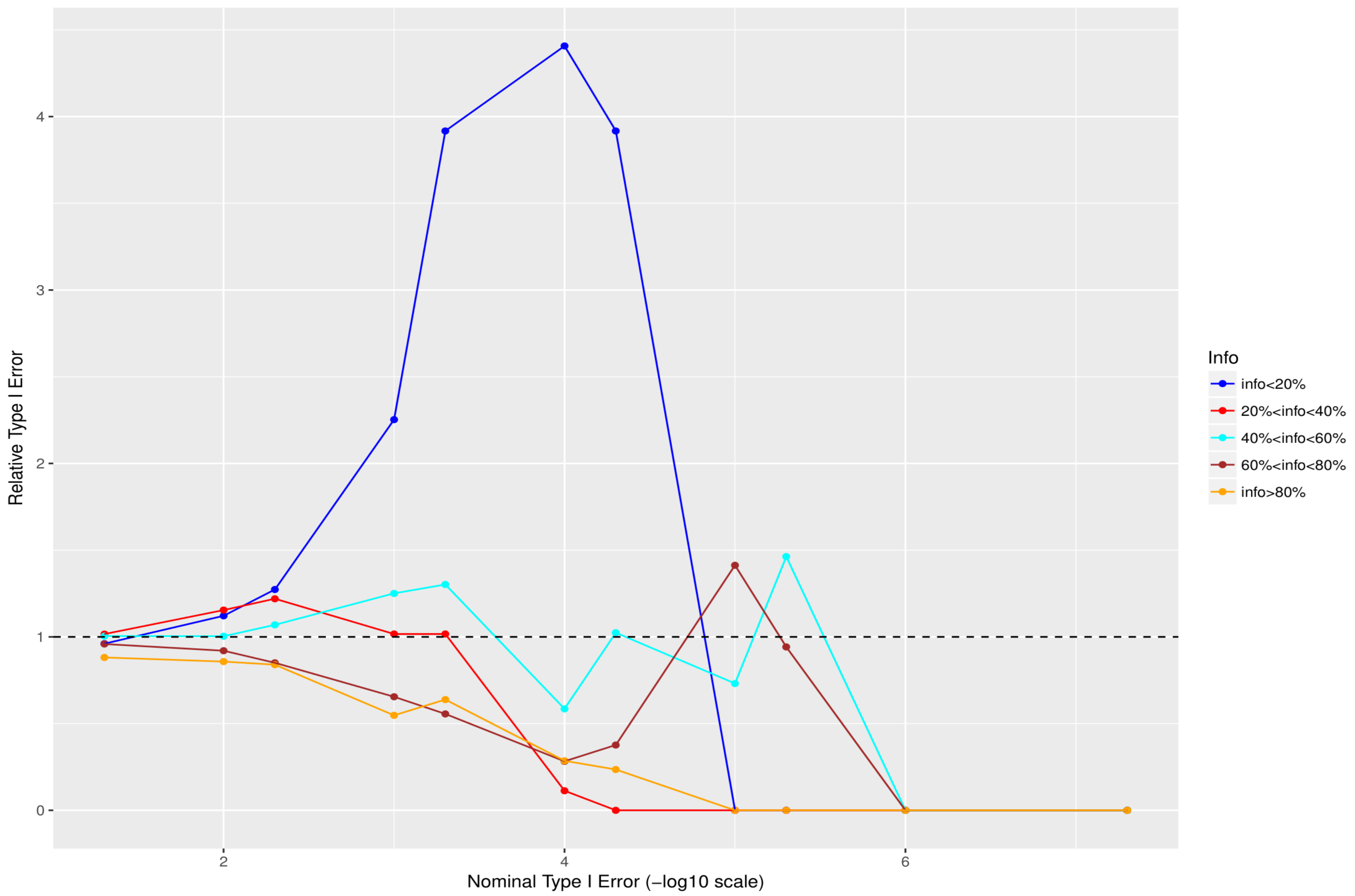


**Fig. S47 Relative size of the test - nullified data for 5%<MAF<10% (the quotient of empirical false positive rate and nominal type I error).**

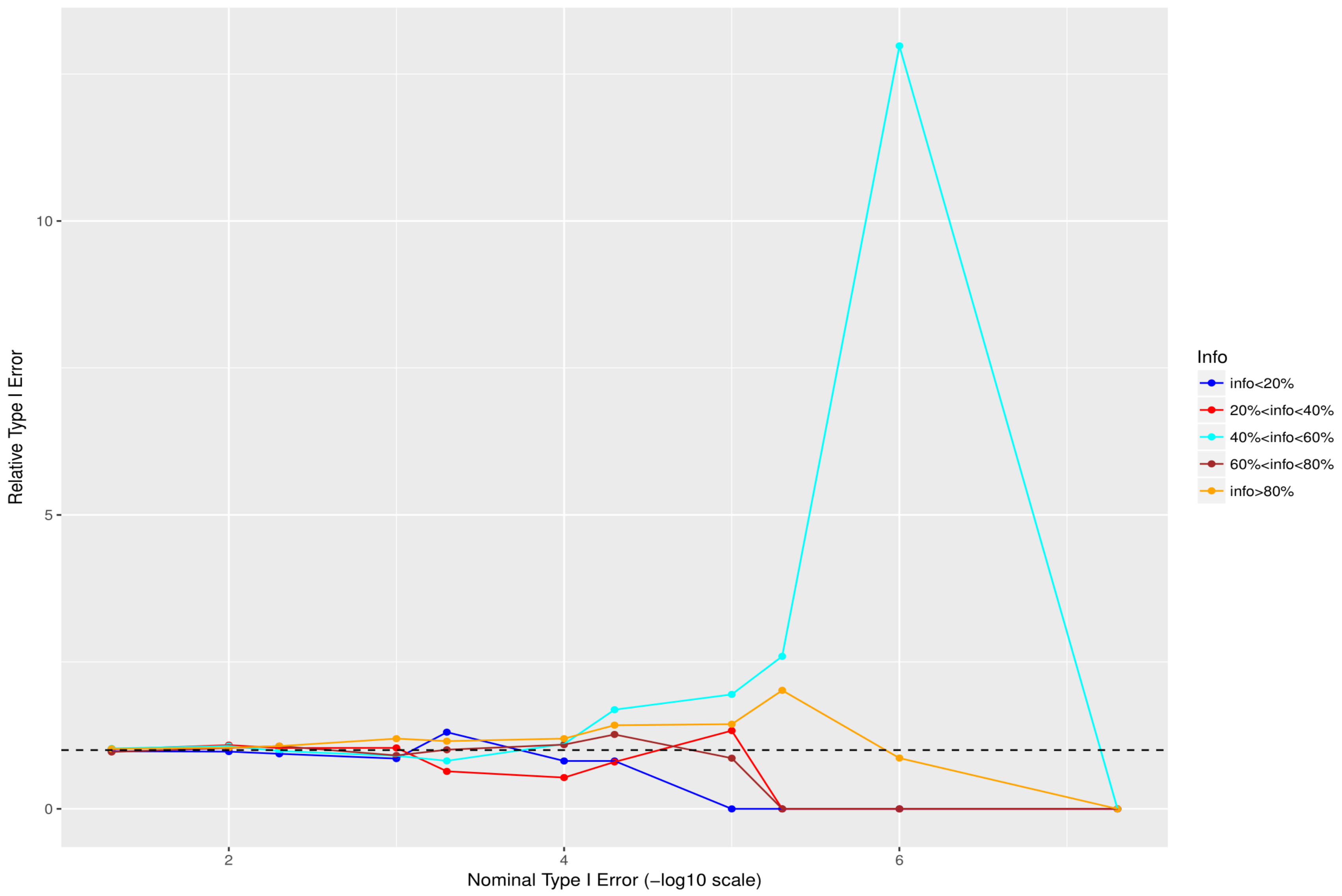


**Fig. S48 Relative size of the test - nullified data for 10%<MAF<50% (the quotient of empirical false positive rate and nominal type I error).**

### S3. Practical Applications.

In this section, we obtained signals by applying DISTMIX2 to association summary statistics as ADHD, AUT, ED, BIP, MDD and SCZ. We construct Manhattans plots for all chromosomes general and for specific chromosomes which we had new signals (Fig. S49-S59). Additionally, in order to investigate the potential risk of genomic inflation, we conducted Q-Q plots for the following three scenarios (i) all the SNPs, (ii) rare SNPs and (iii) common SNPS (Fig. S60-S76). Finally, we compare distmix2 with ARDISS [1]. Since ARDISS software does not provide minor allele frequency information (MAF) and info estimation we subset the imputed signals according to distimix2 for $maf > 0.05$ (Fig. S77-S83).

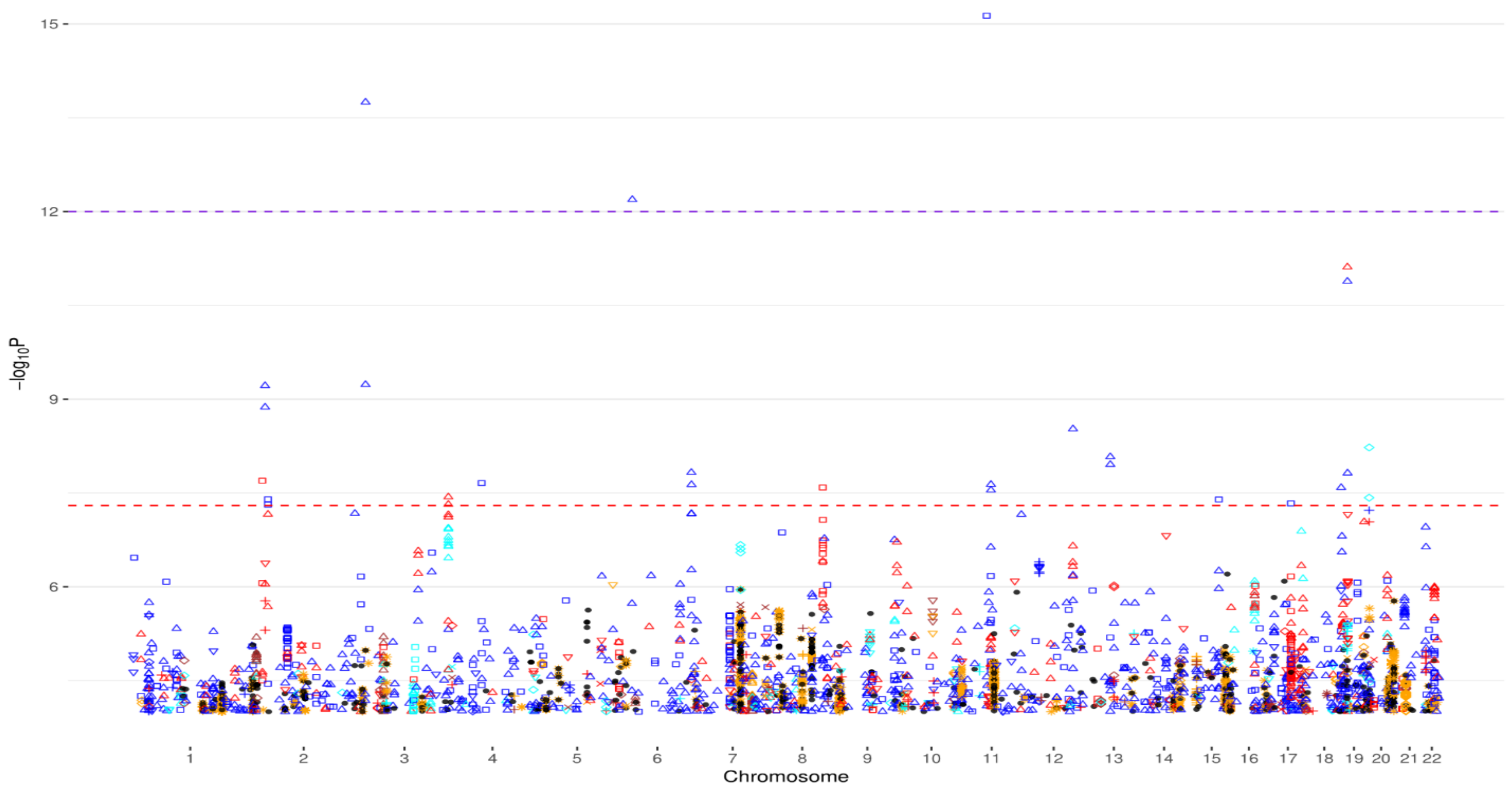


**Fig. S49 Manhattan plot of chromosomes 1-22 for ADHD. ● denotes reported signals and the remain symbols and colors denote DISTMIX2 imputed signals. Among imputed signals blue denotes info<0.2, red denotes 0.2<info<0.4, cyan denotes 0.4<info<0.6, brown denotes 0.6<info<0.8, orange denotes info>0.8, □ denotes MAF <0.05%, △ denotes 0.05%<MAF<0.5%, ▽ denotes 0.5%<MAF<1%, + denotes 1%<MAF<2%, ◊ denotes 2%<MAF<5%, $x$ denotes 5%<MAF<10% and ⁕ denotes 10%<MAF<50%. The red line is the default genome-wide threshold of $p = 5 * 10^{-8}$, which is applicable common SNPs with moderate to large Info values. The purple line at $p = 10^{-12}$ is the threshold to be used for rare/very rare variants.**

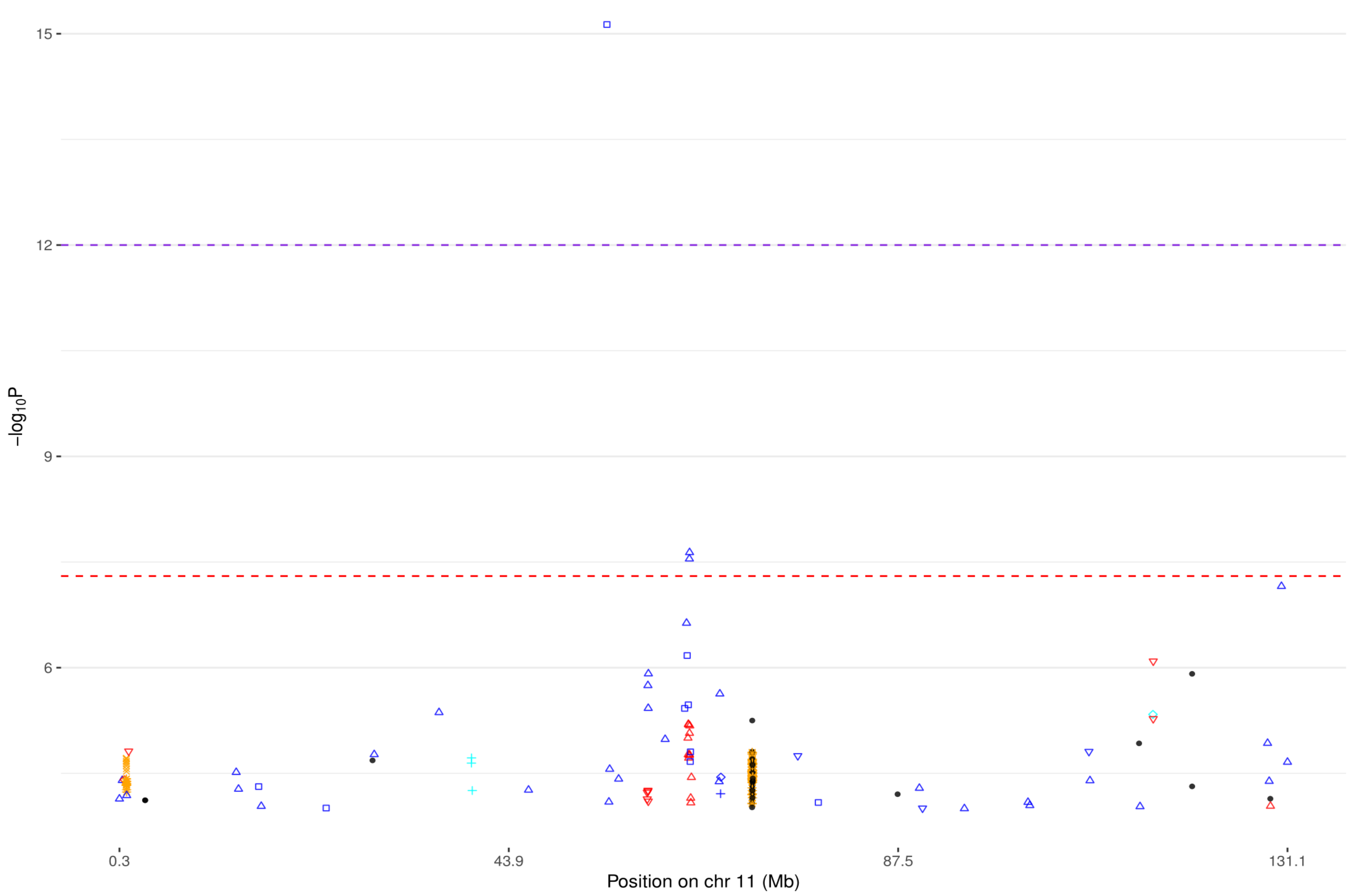


**Fig. S50 Manhattan plot for chromosome 11 for ADHD (see Fig. S49 for background).**

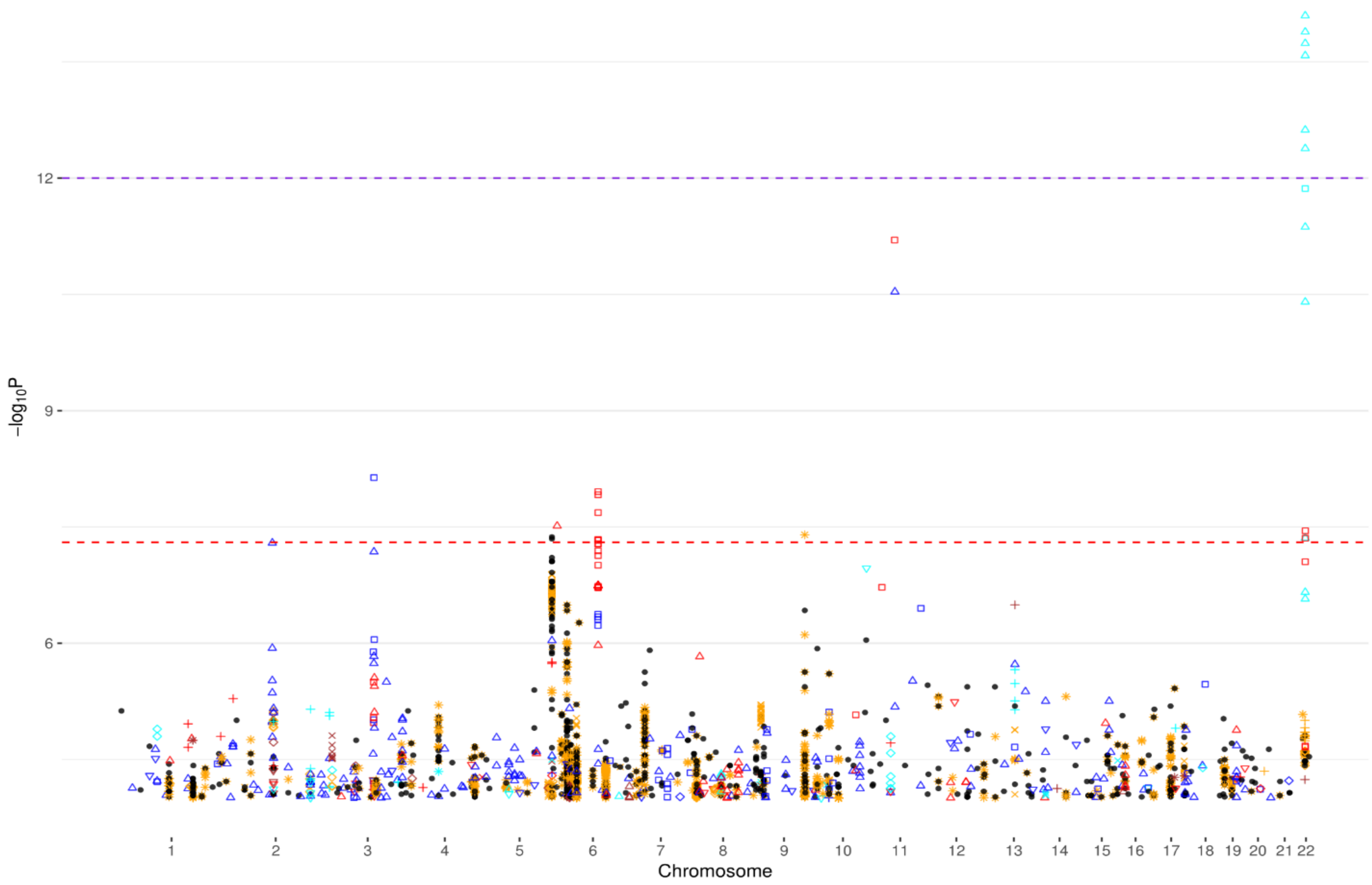


**Fig. S51 Manhattan plot of chromosomes 1-22 for AUT (see Fig. S49 for background).**

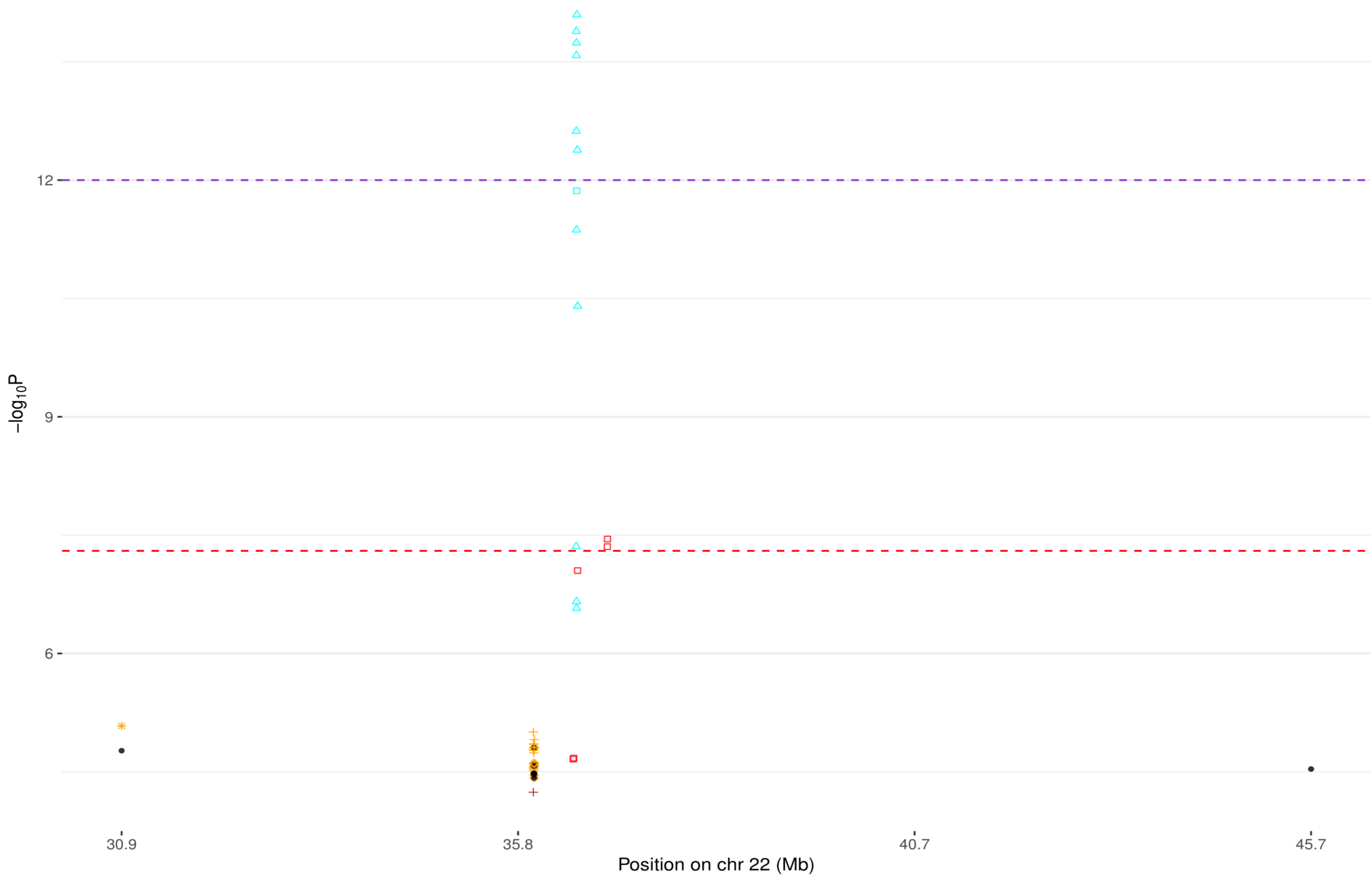


**Fig. S52 Manhattan plot of chromosome 22 for AUT (see Fig. S49 for background).**

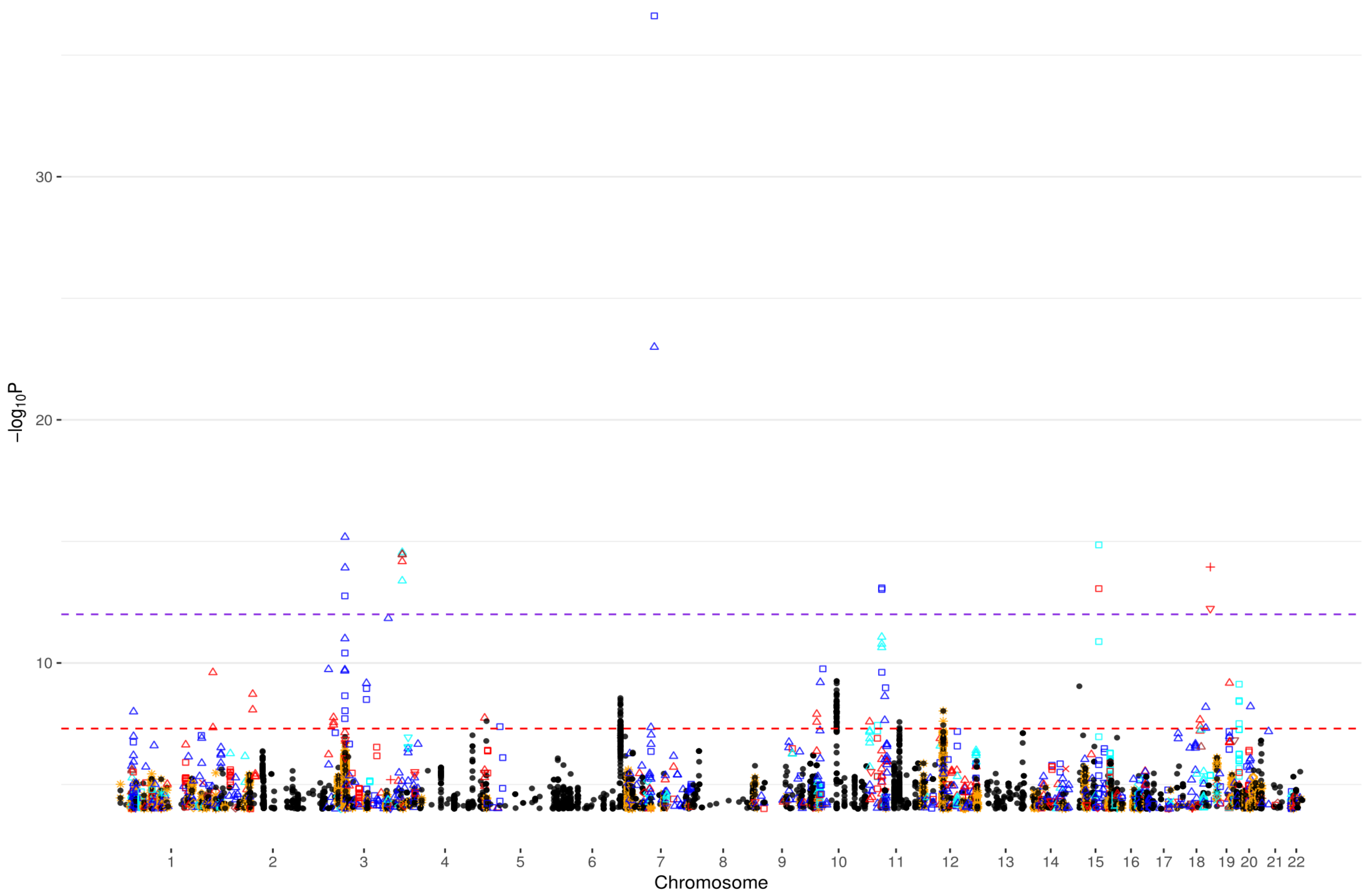


**Fig. S53 Manhattan plot for chromosome 1-22 for BIP (see Fig. S49 for background).**

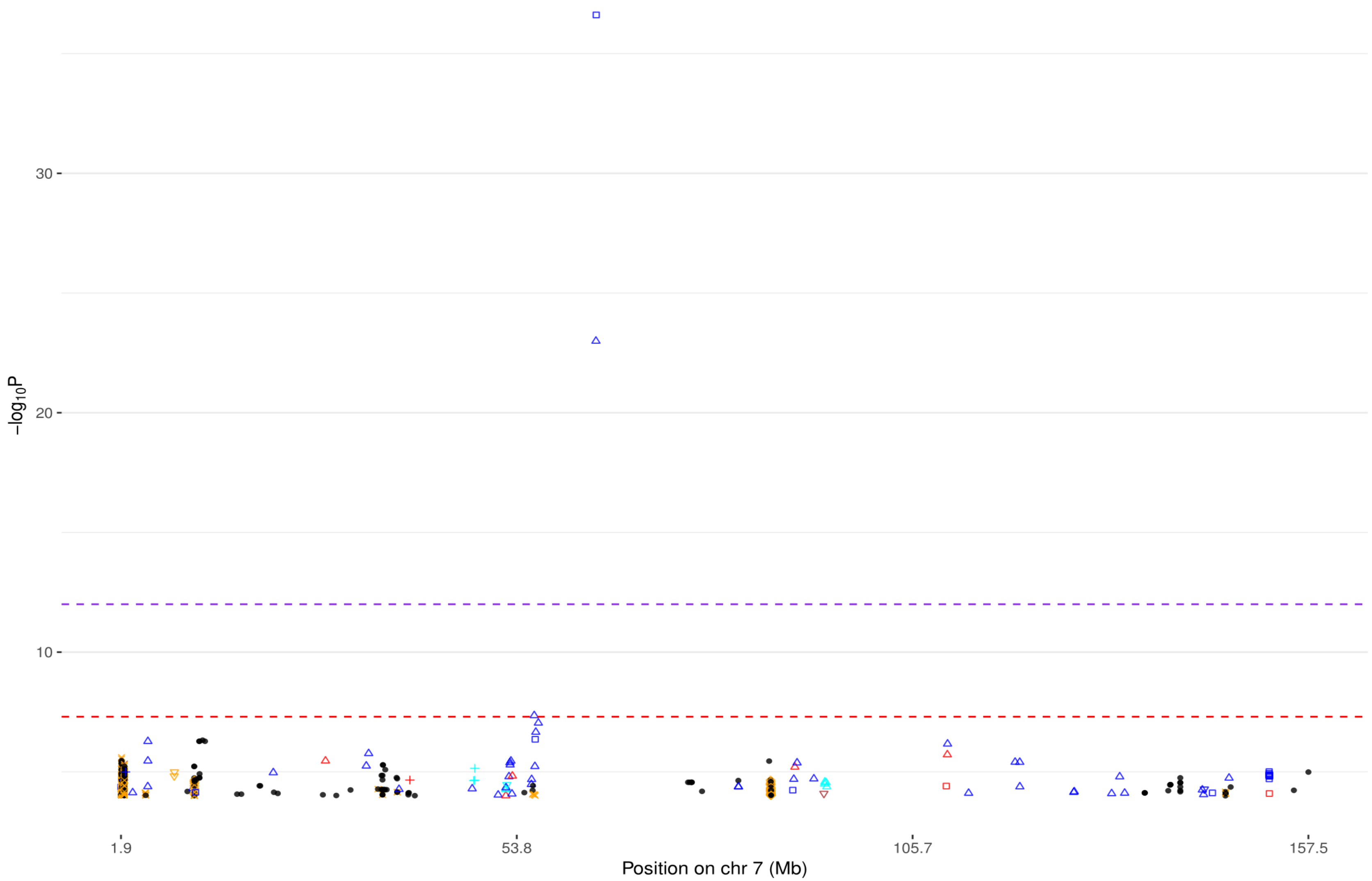


**Fig. S54 Manhattan plot for chromosome 7 for BIP (see Fig. S49 for background).**

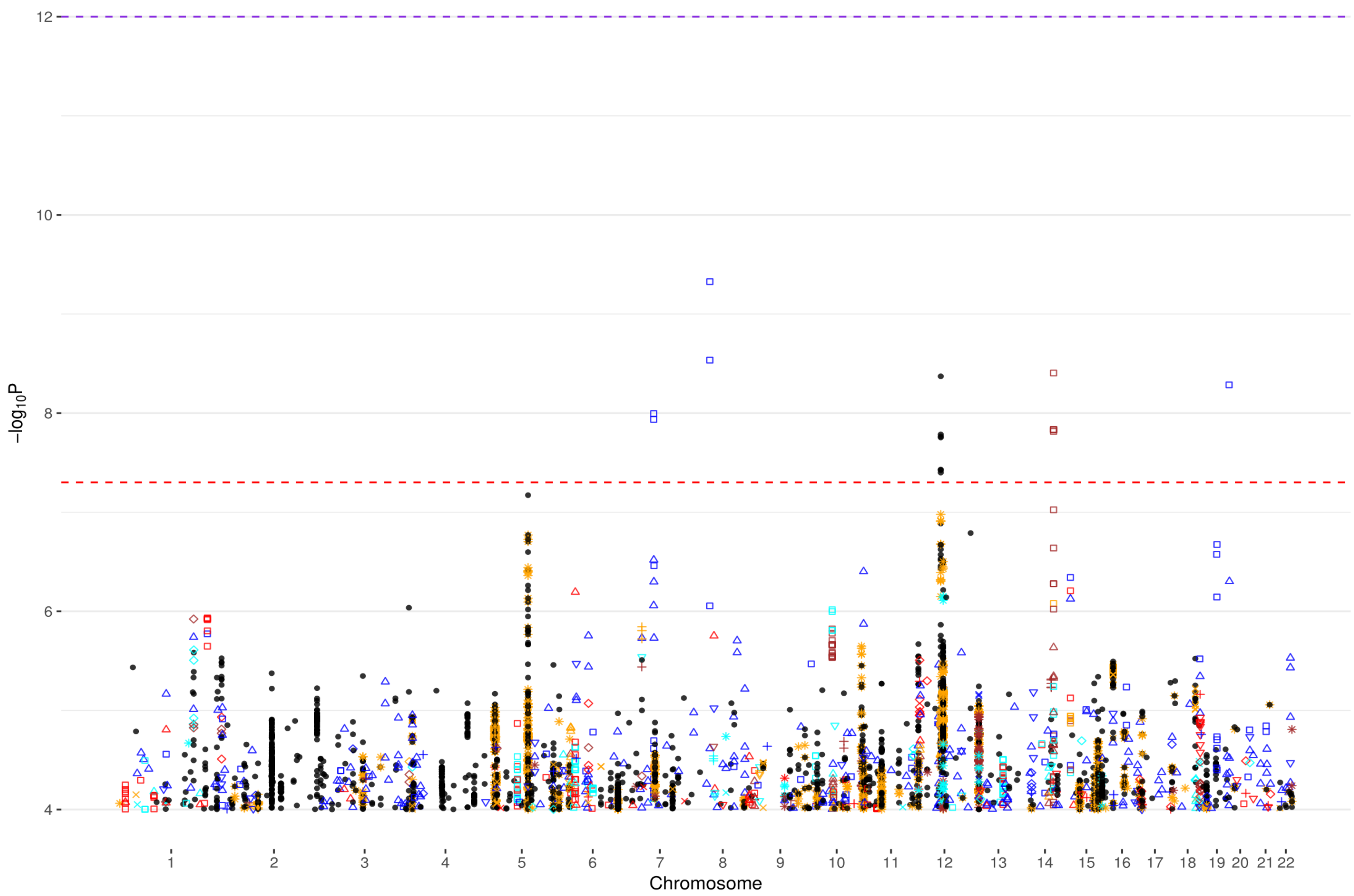


**Fig. S55 Manhattan plot of chromosomes 1-22 for ED (see Fig. S49 for background).**

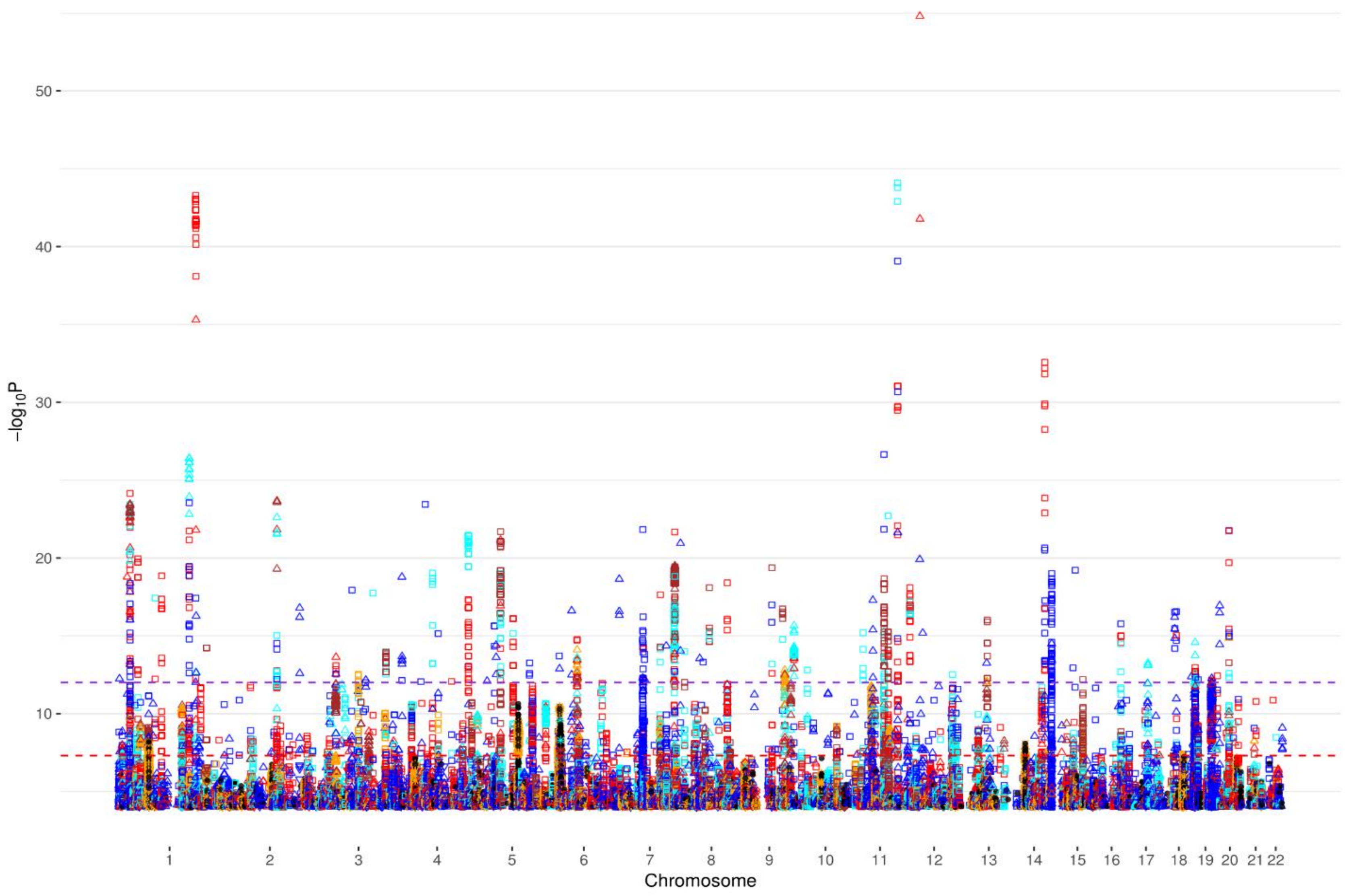


**Fig. S56 Manhattan plot of chromosomes 1-22 for MDD (see Fig. S49 for background).**

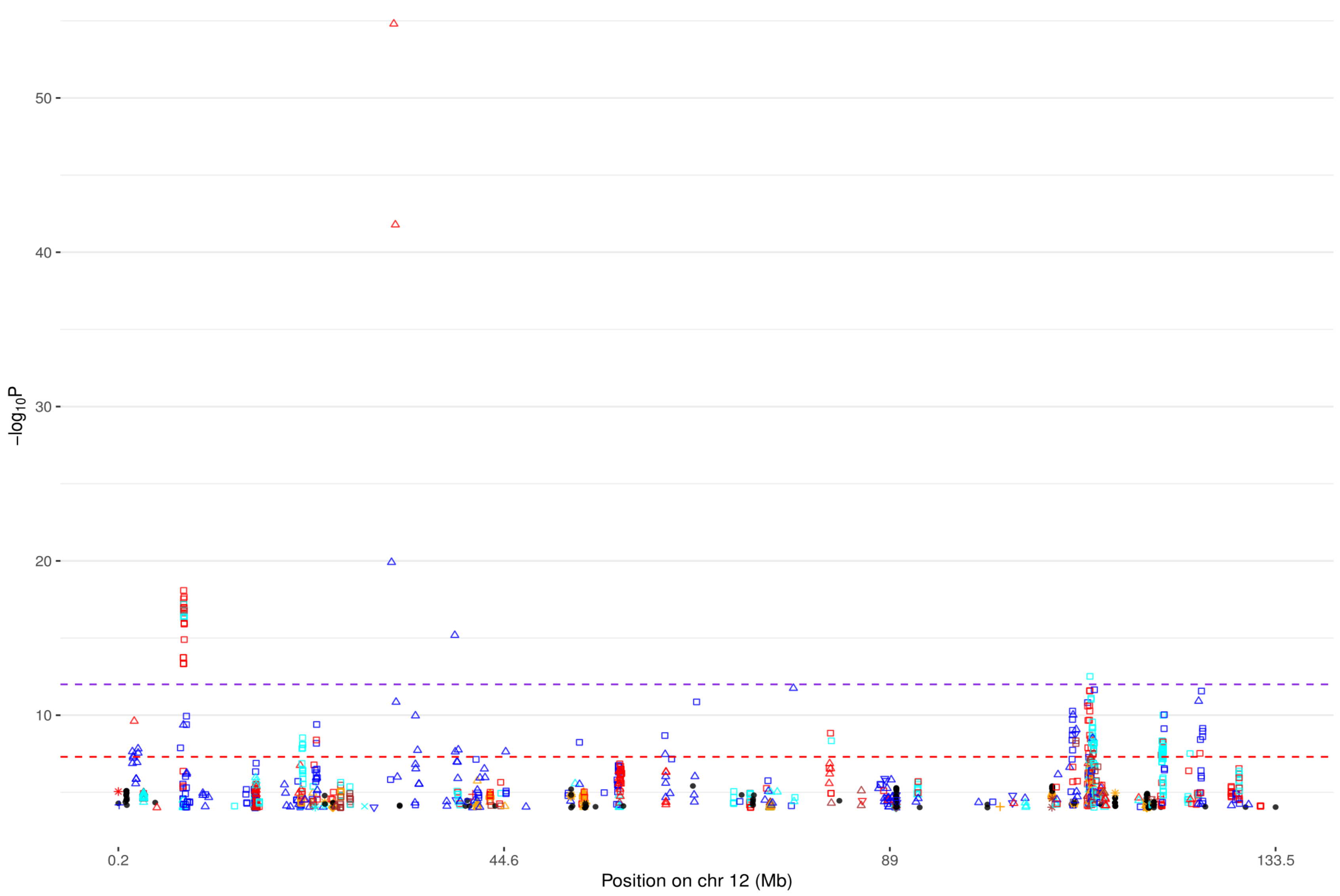


**Fig. S57 Manhattan plot of chromosome 12 for MDD (see Fig. S49 for background).**

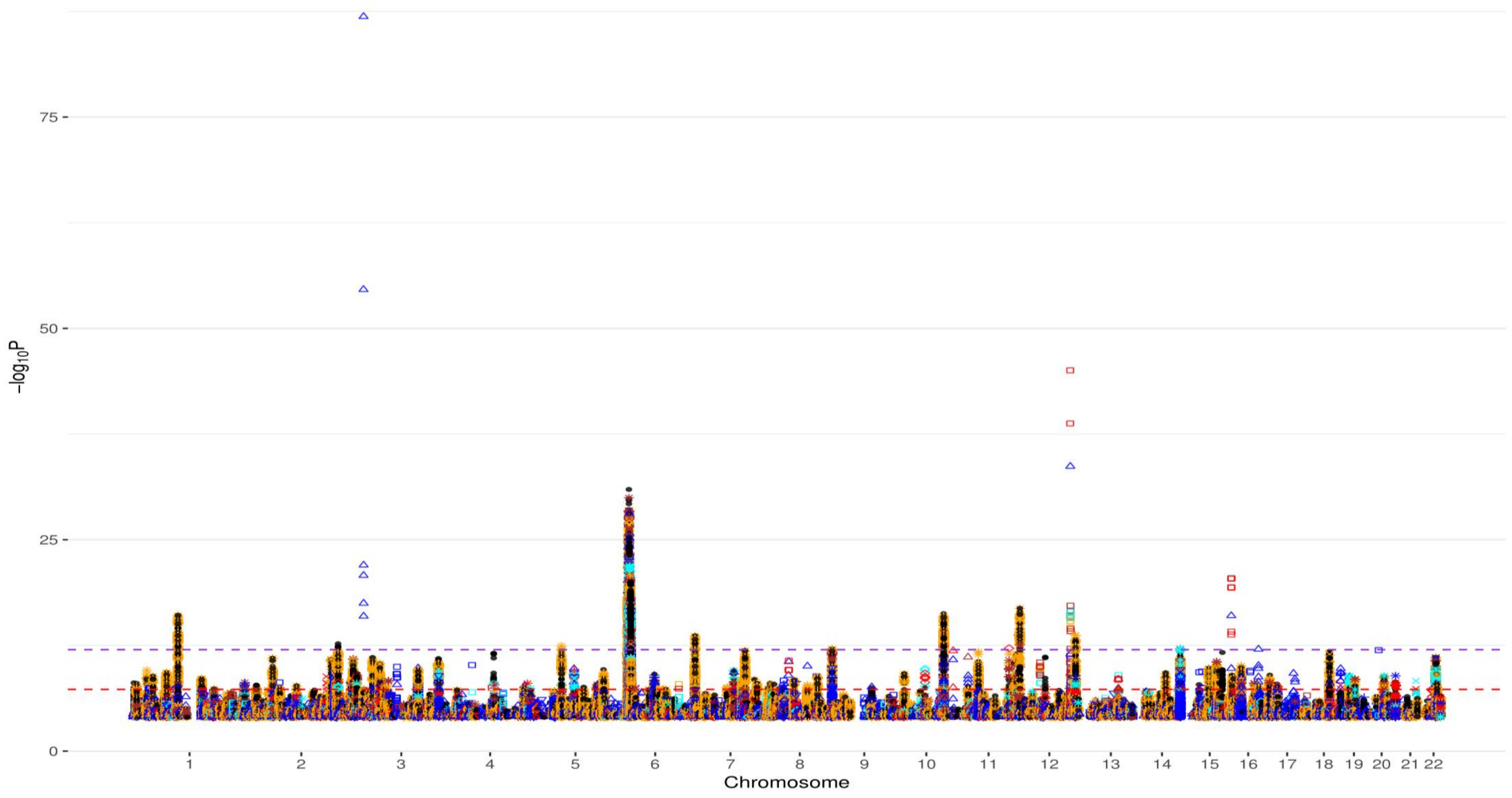


**Fig. S58 Manhattan plot of chromosomes 1-22 for SCZ (see Fig. S49 for background).**

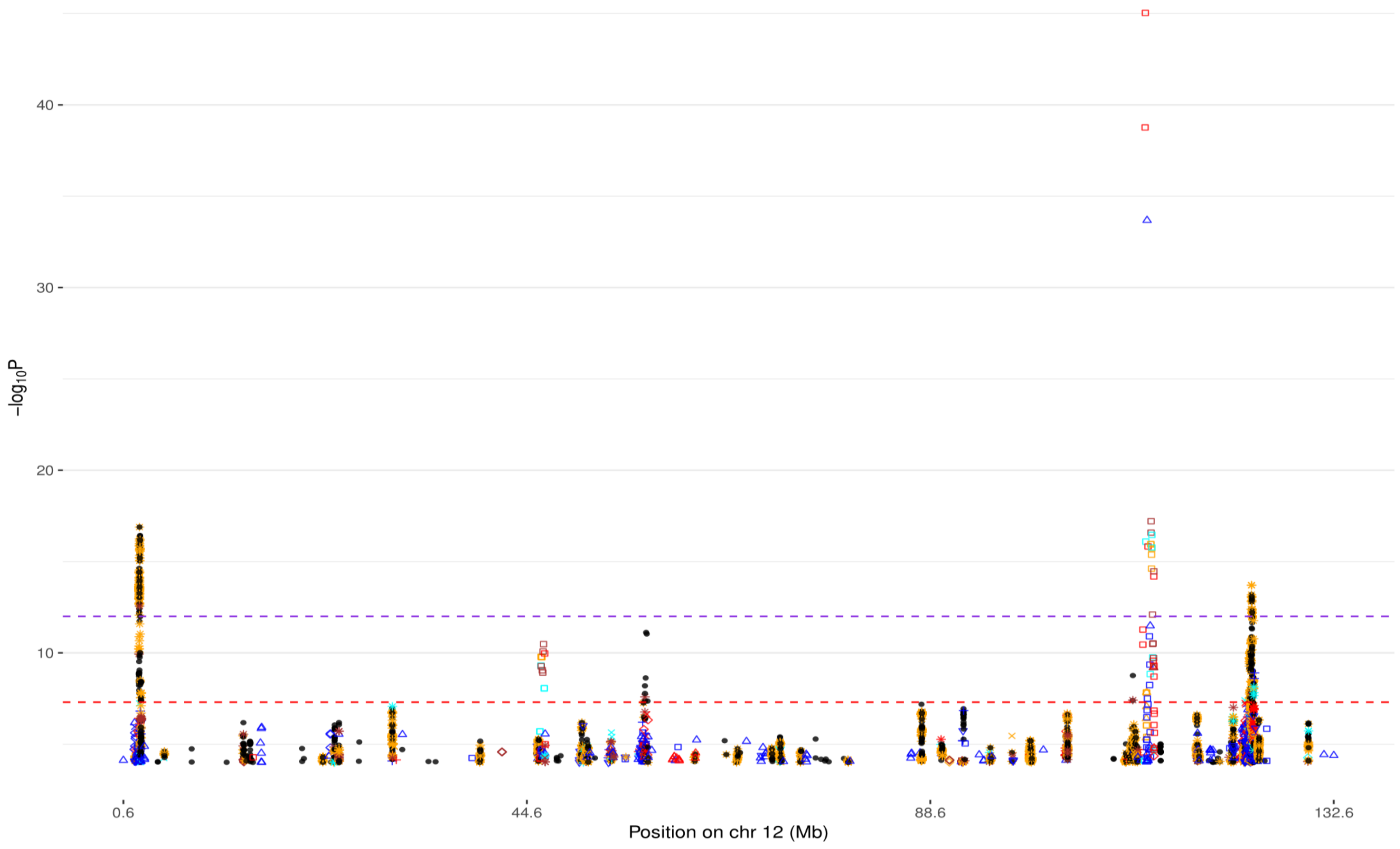


**Fig. S59 Manhattan plot of chromosome 12 for SCZ (see Fig. S49 for background).**

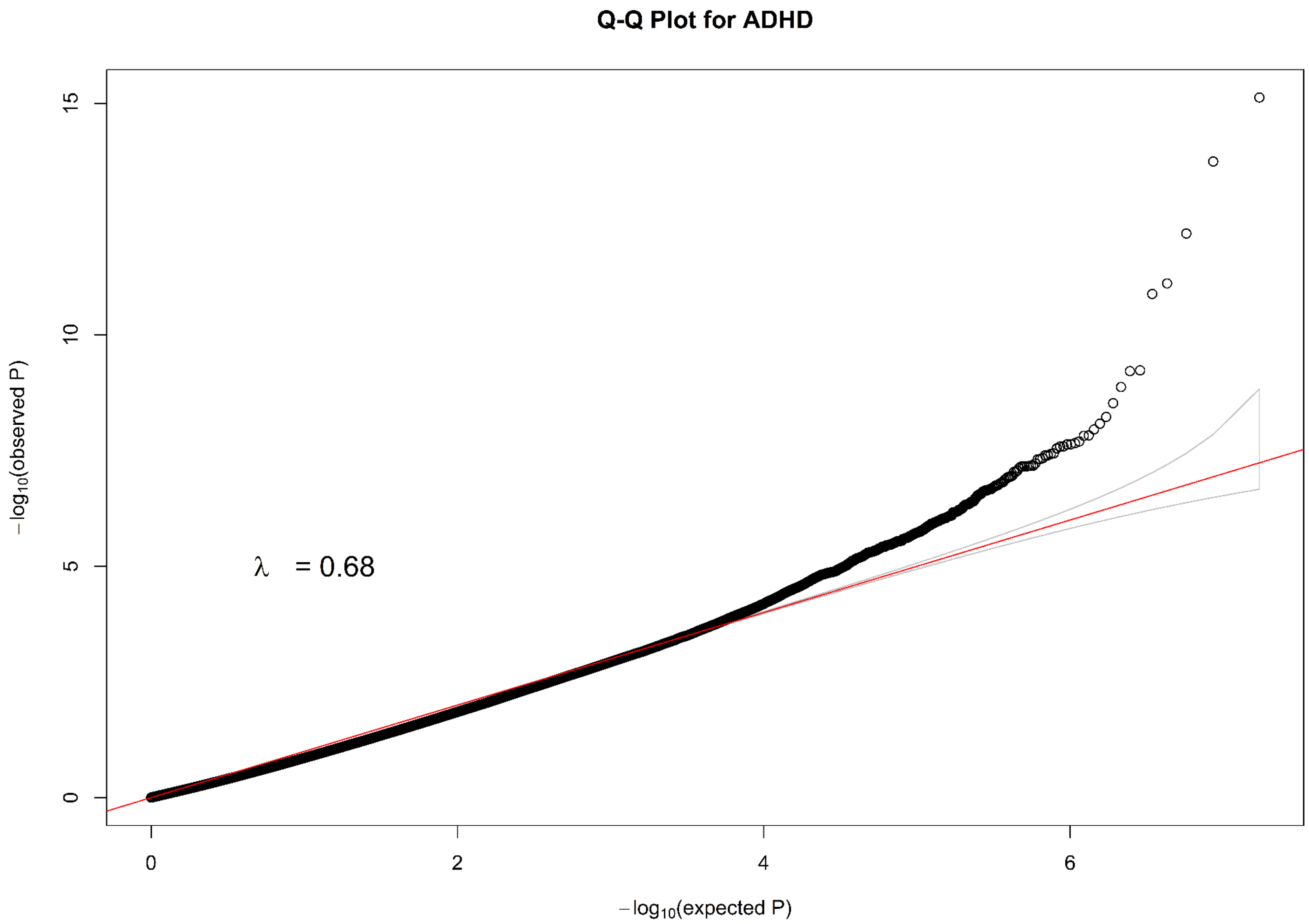


**Fig. S60 Q-Q plot for all SNPs of ADHD.**

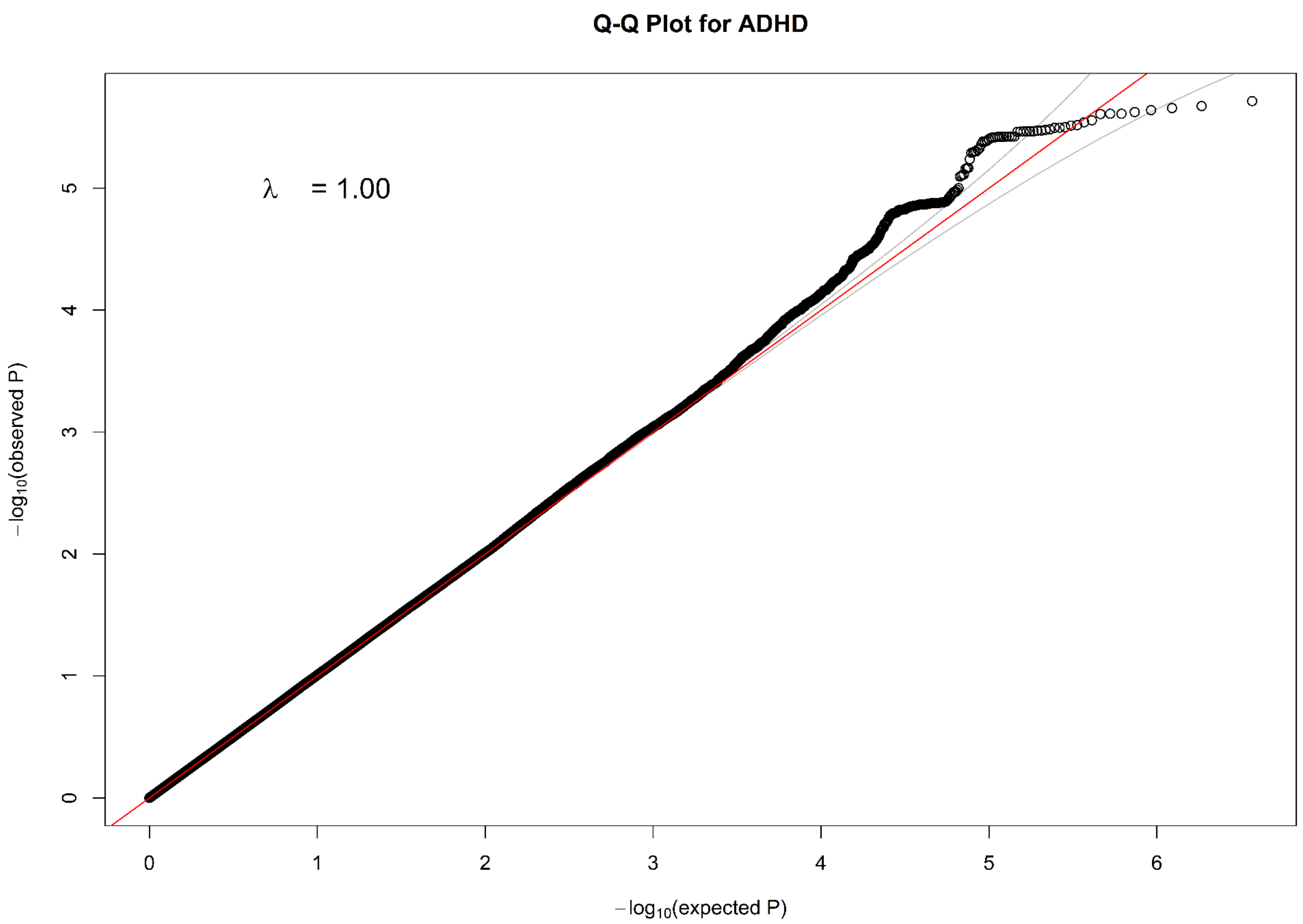


**Fig. S61 Q-Q plot for common SNPs of ADHD.**

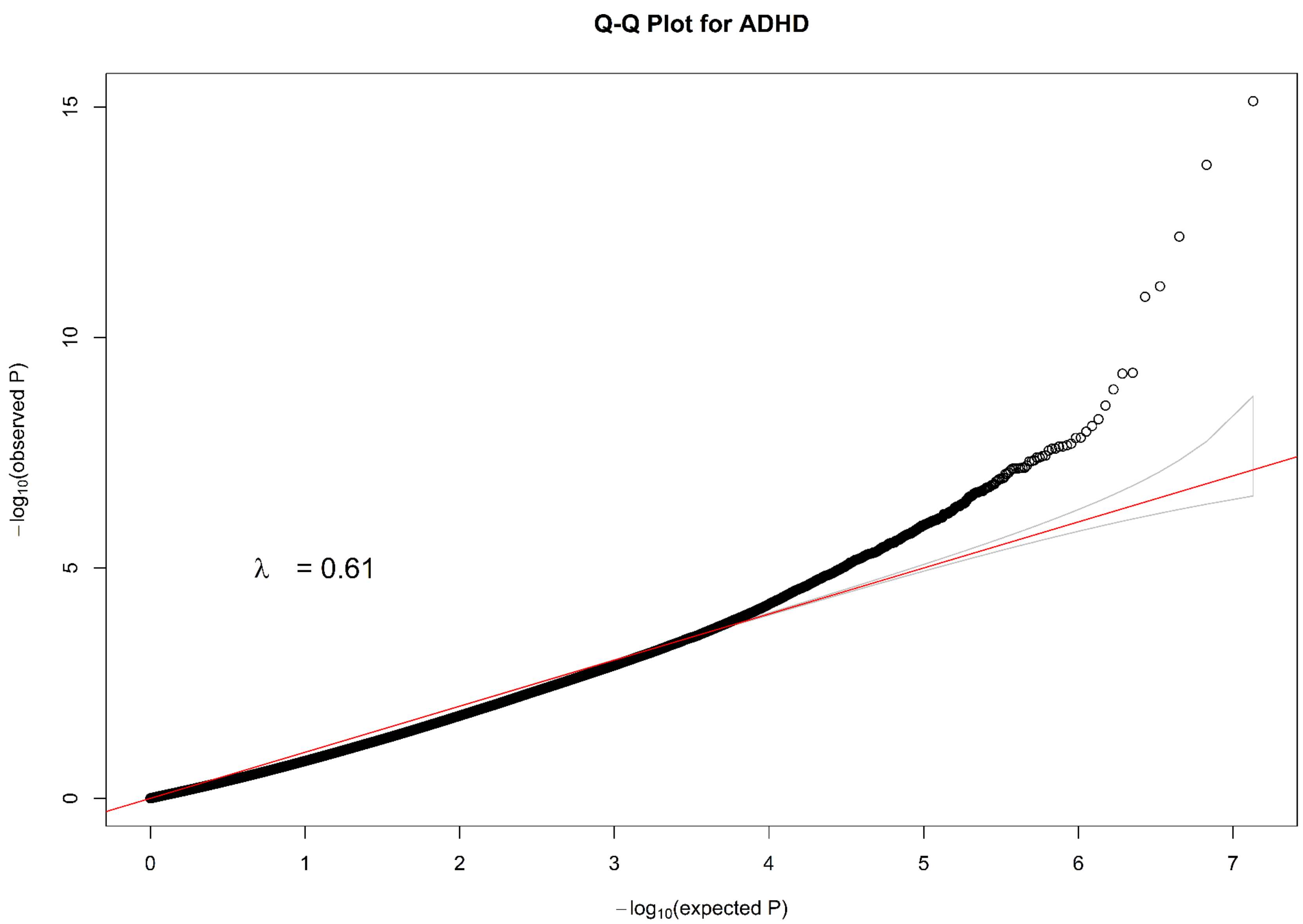


**Fig. S62 Q-Q plot for rare SNPs of ADHD.**

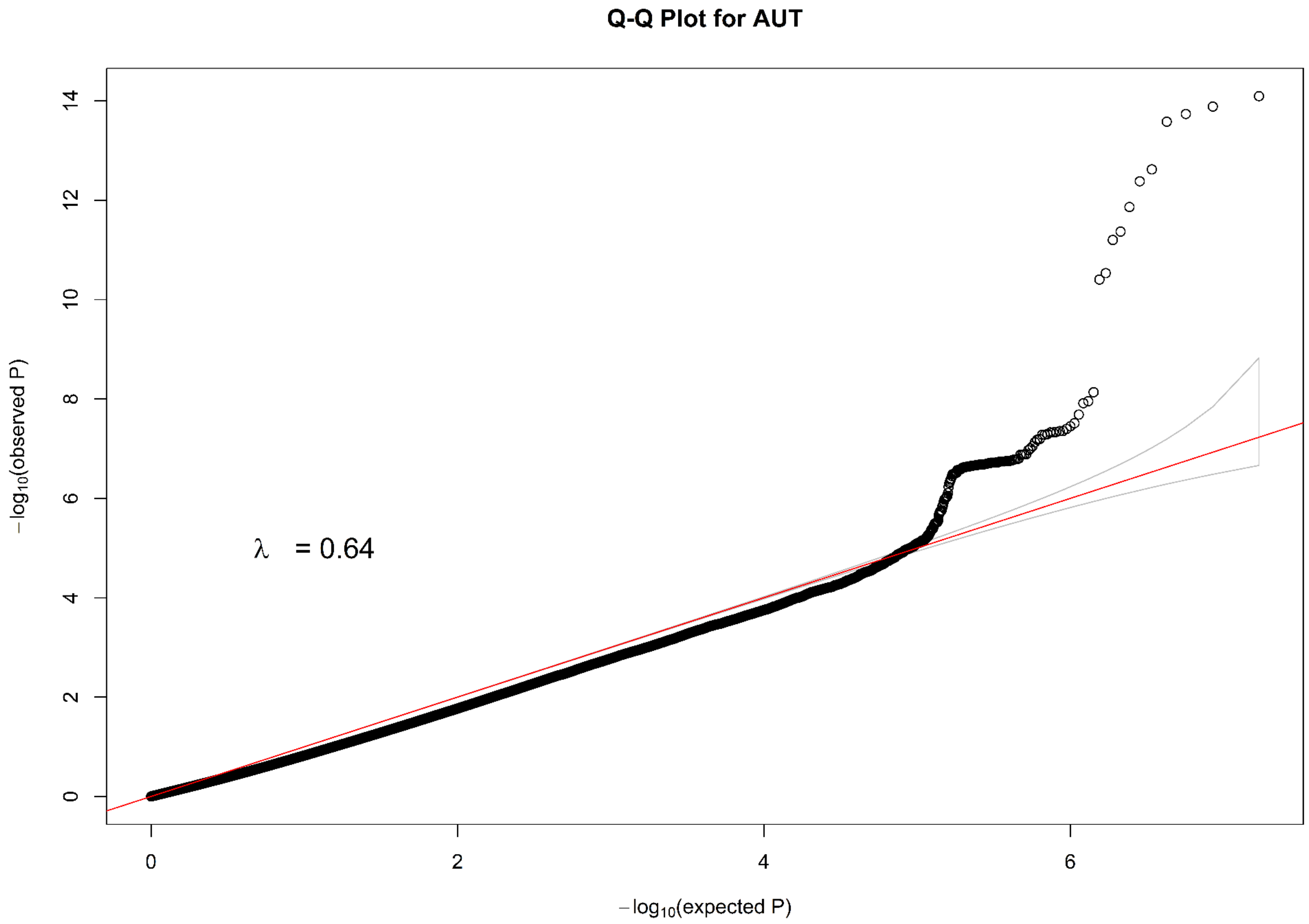


**Fig. S63 Q-Q plot for all SNPs of AUT.**

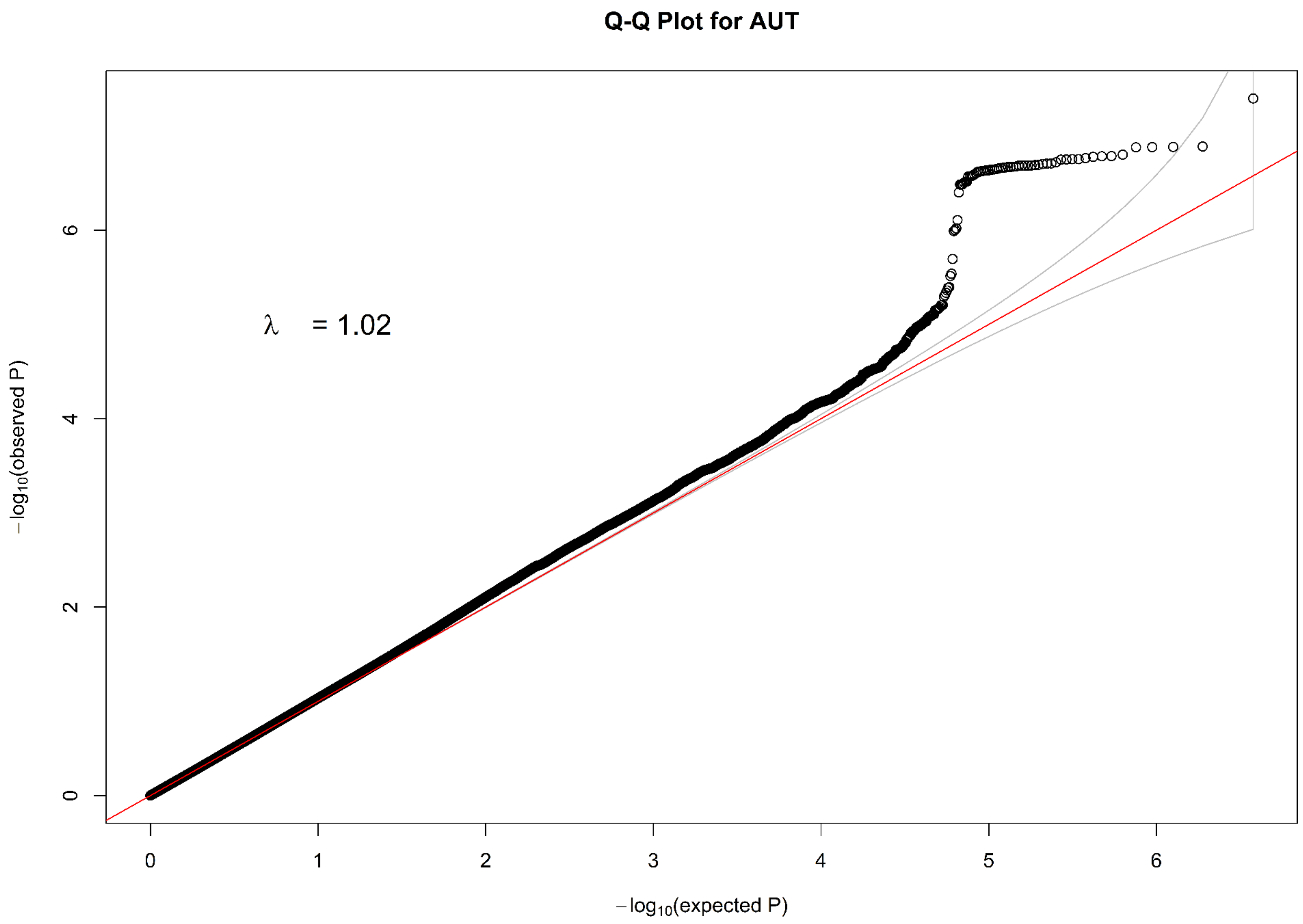


**Fig. S64 Q-Q plot for common SNPs of AUT.**

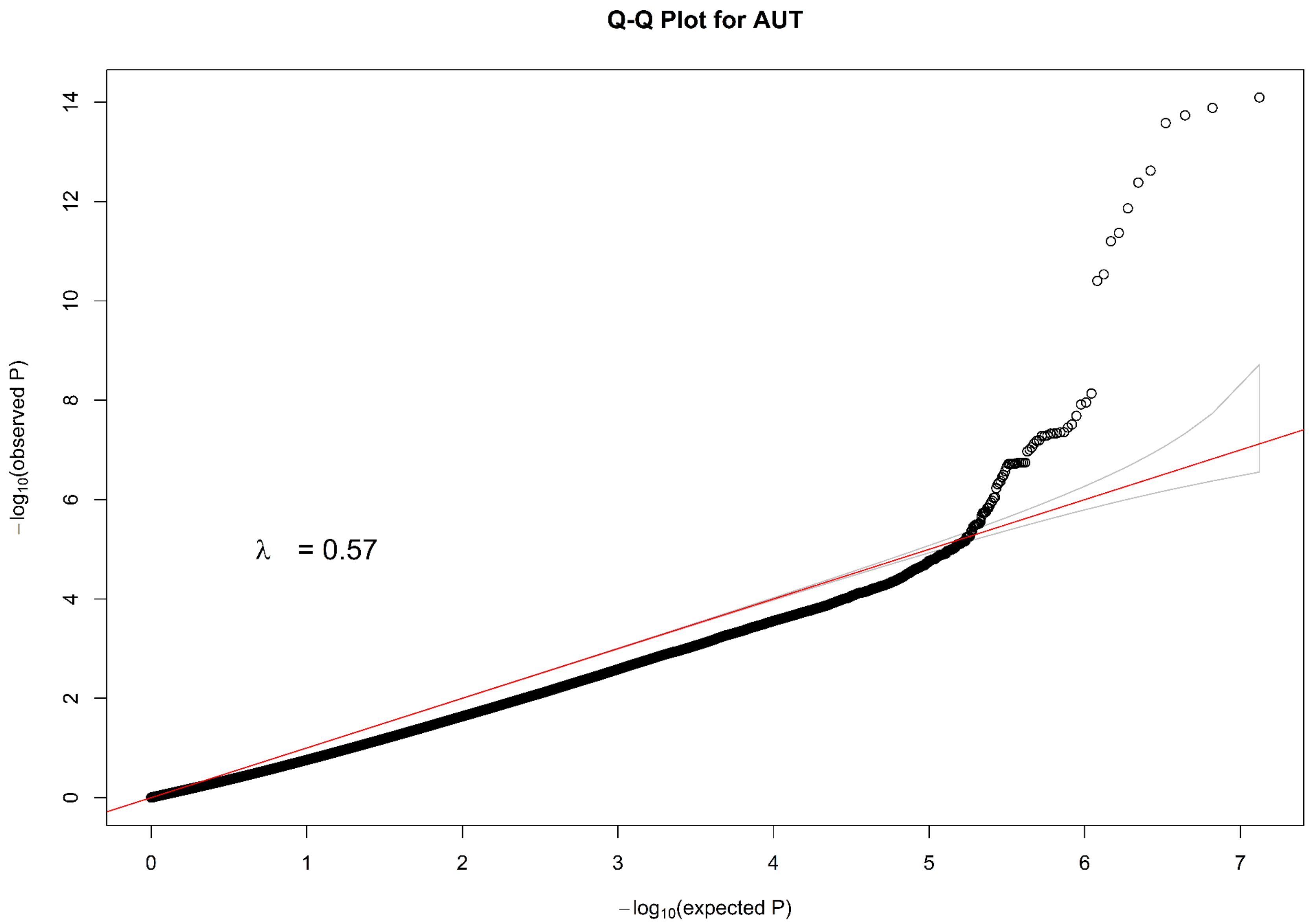


**Fig. S64 Q-Q plot for rare SNPs of AUT.**

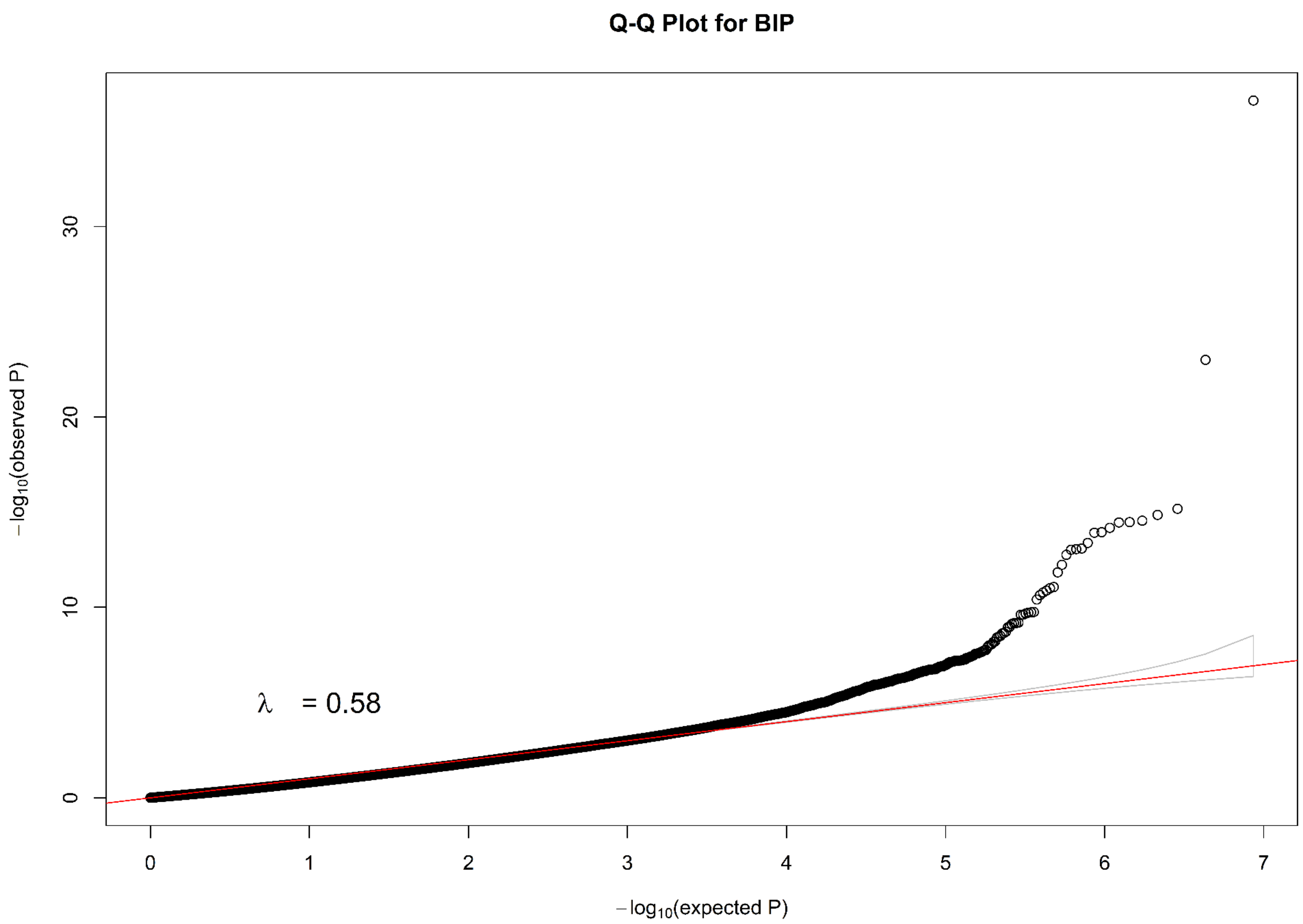


**Fig. S65 Q-Q plot for all SNPs of BIP.**

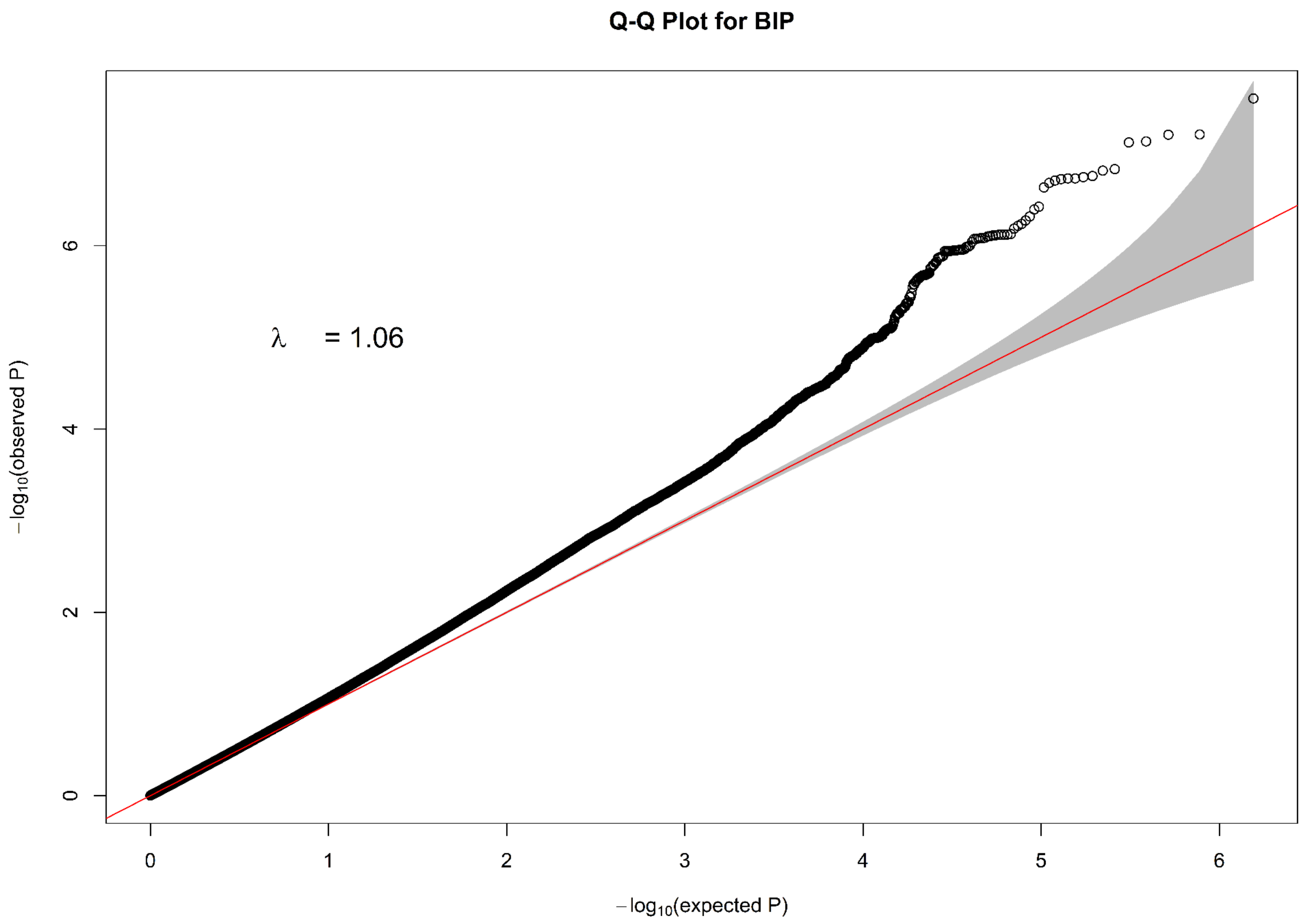


**Fig. S66 Q-Q plot for common SNPs of BIP.**

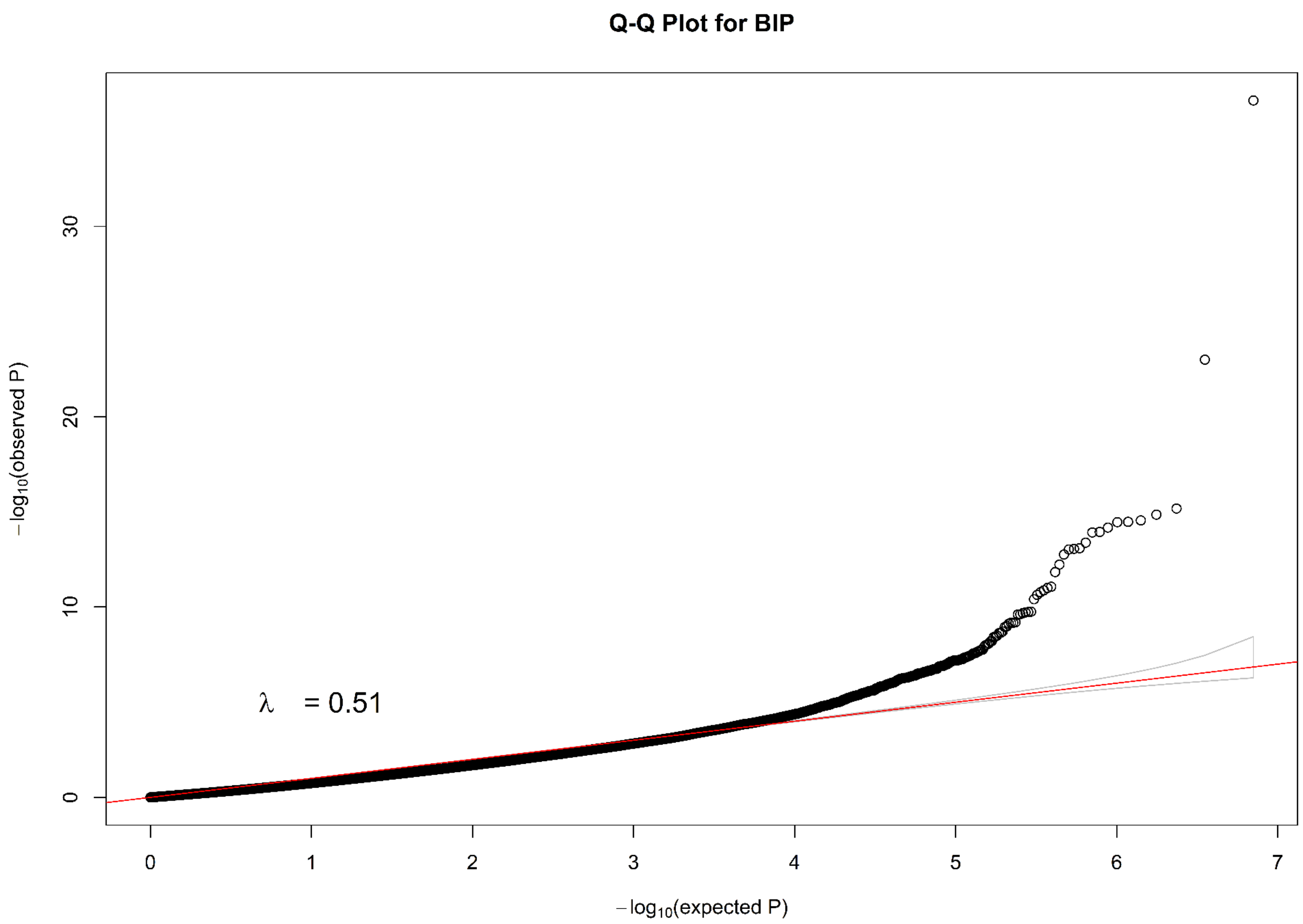


**Fig. S67 Q-Q plot for rare SNPs of BIP.**

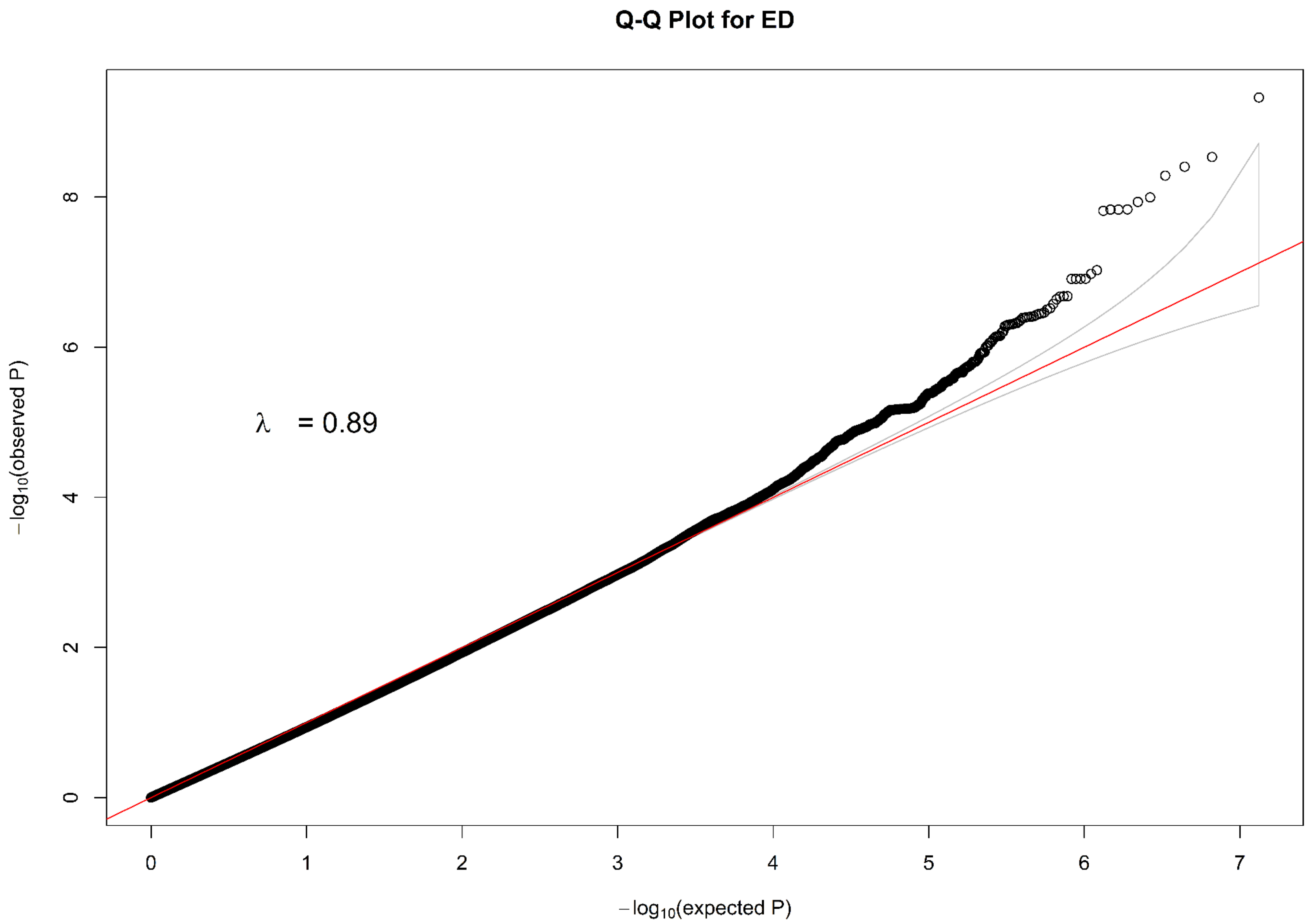


**Fig. S68 Q-Q plot for all SNPs of ED.**

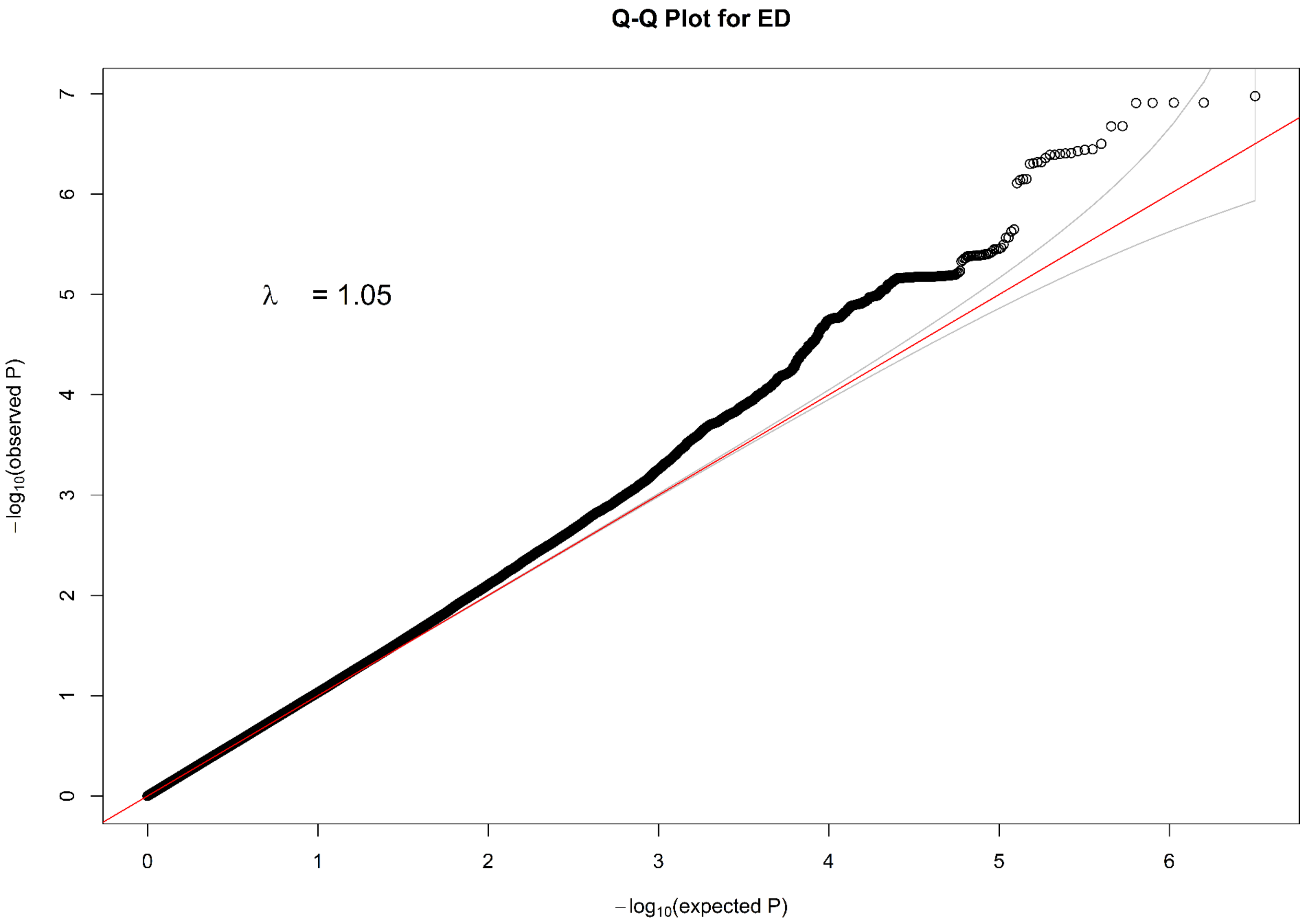


**Fig. S69 Q-Q plot for common SNPs of ED.**

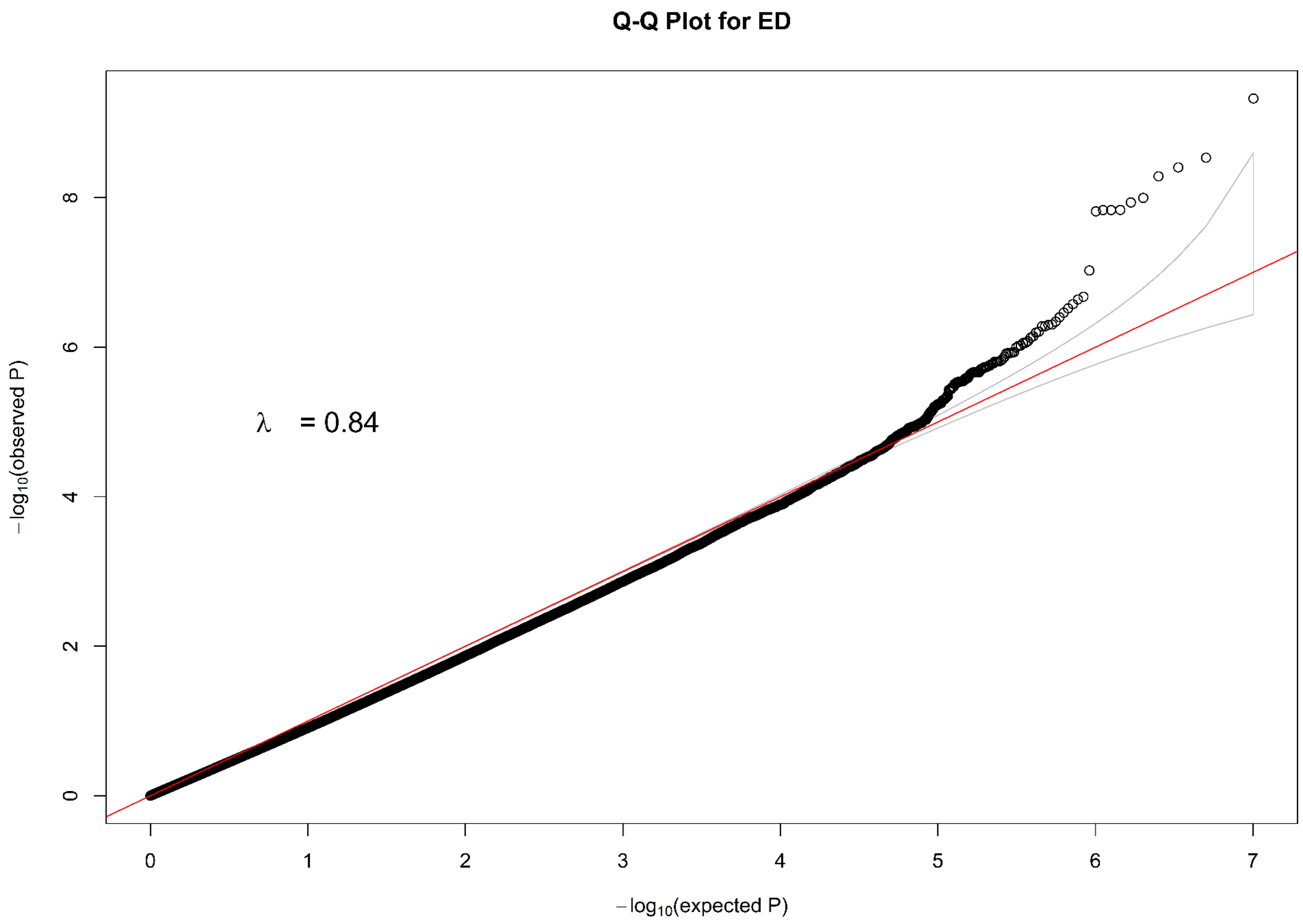


**Fig. S70 Q-Q plot for rare SNPs of ED.**

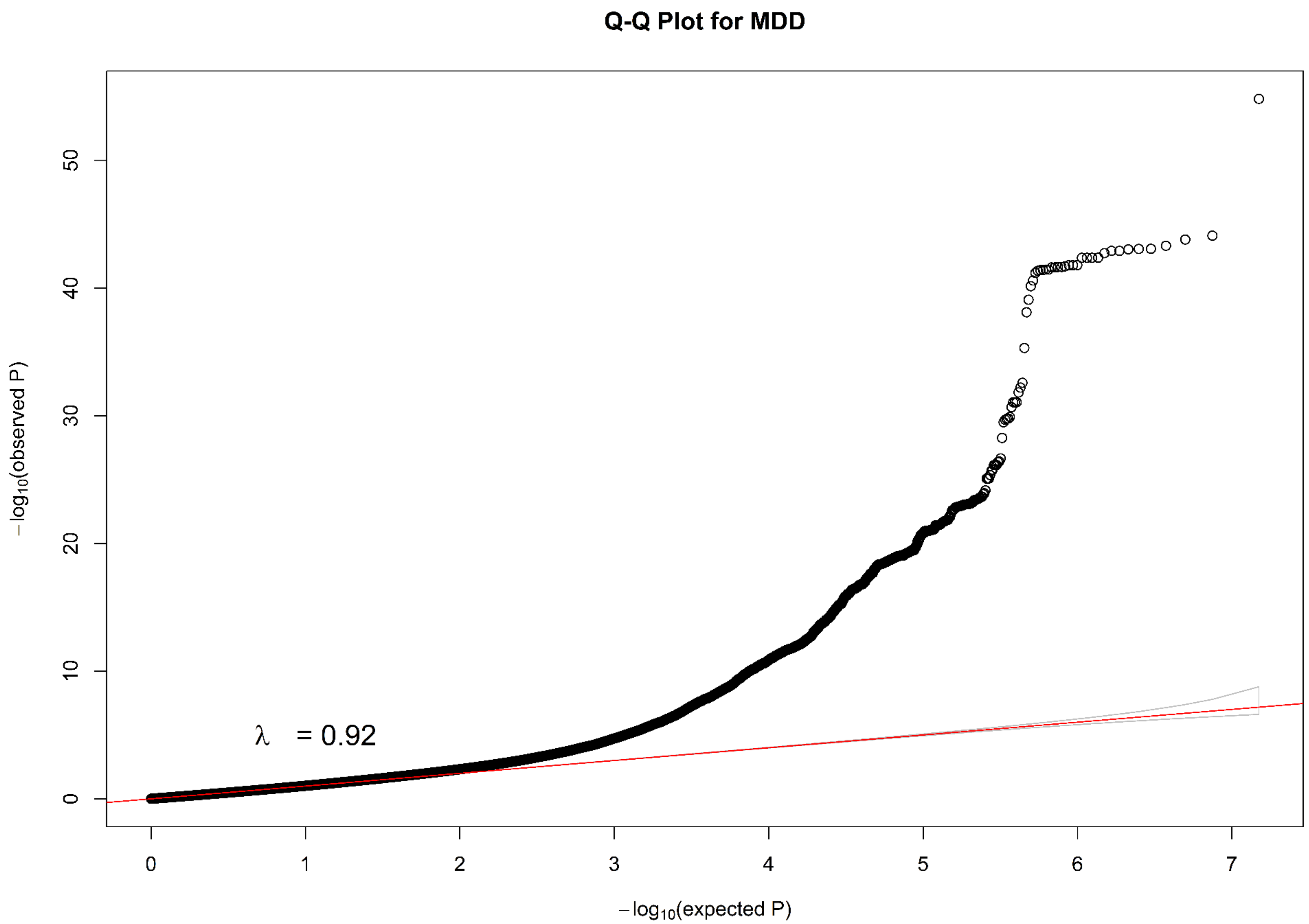


**Fig. S71 Q-Q plot for all SNPs of MDD.**

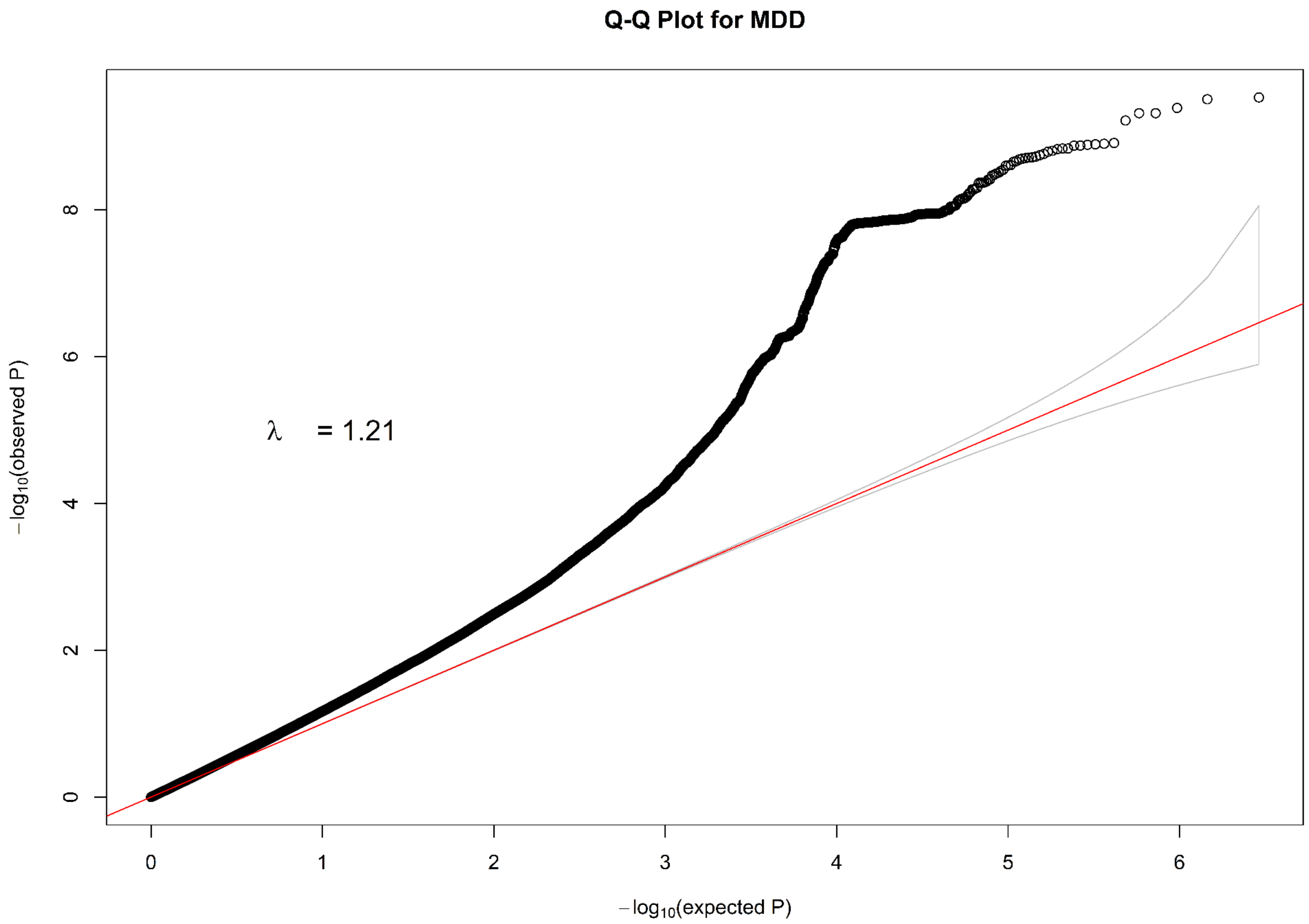


**Fig. S72 Q-Q plot for common SNPs of MDD.**

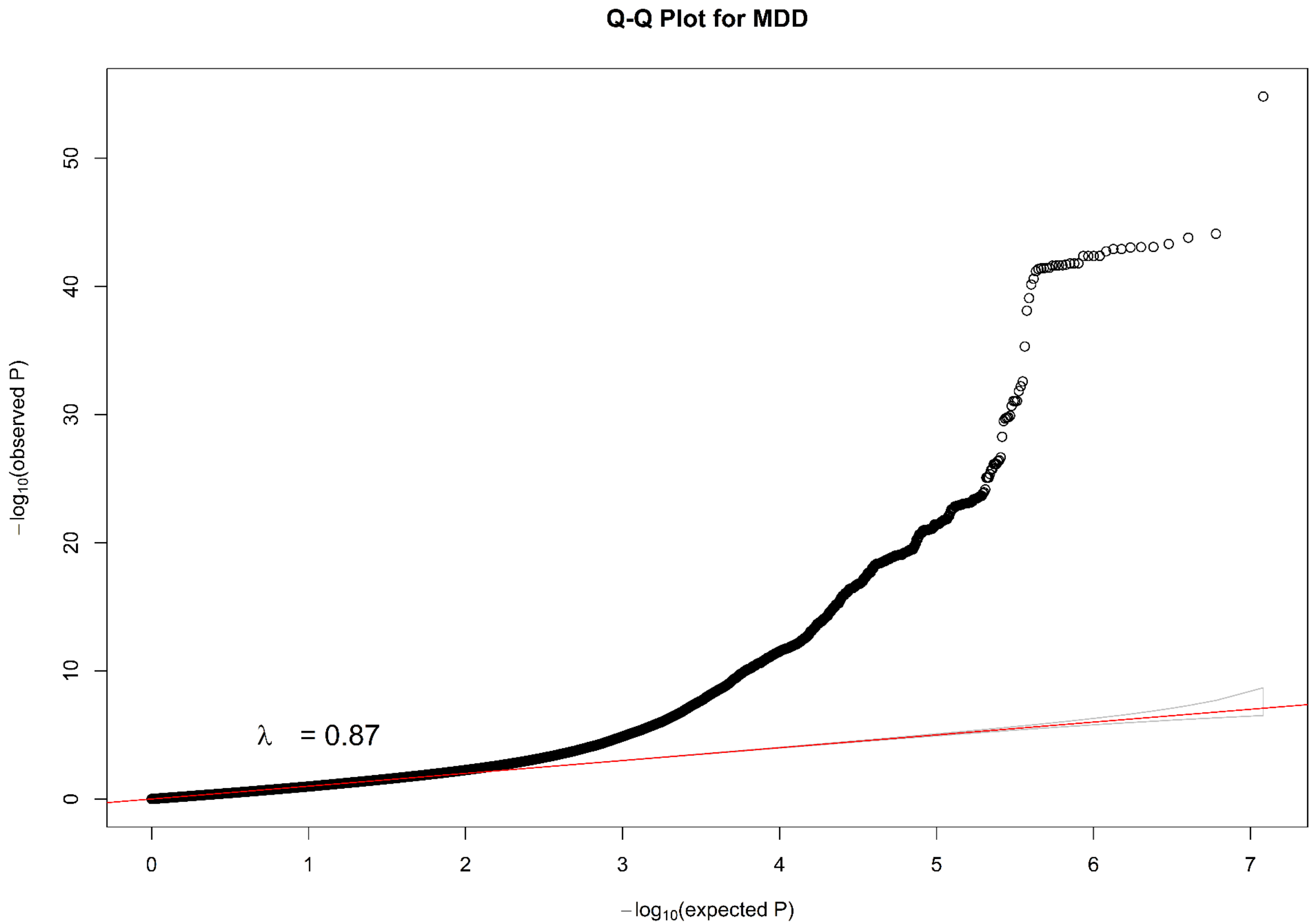

**Fig. S73 Q-Q plot for rare SNPs of MDD.**

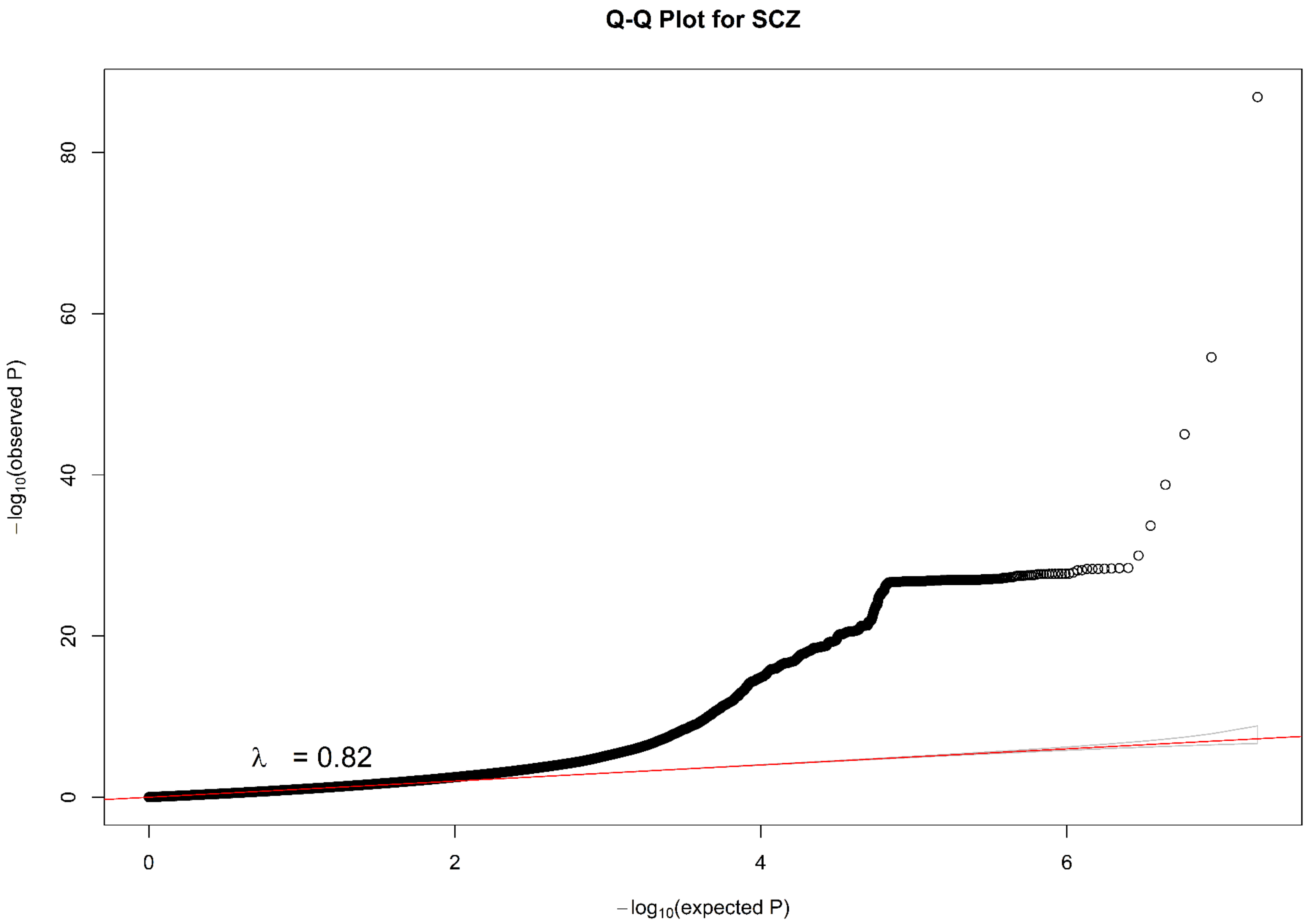


**Fig. S74 Q-Q plot for all SNPs of SCZ.**

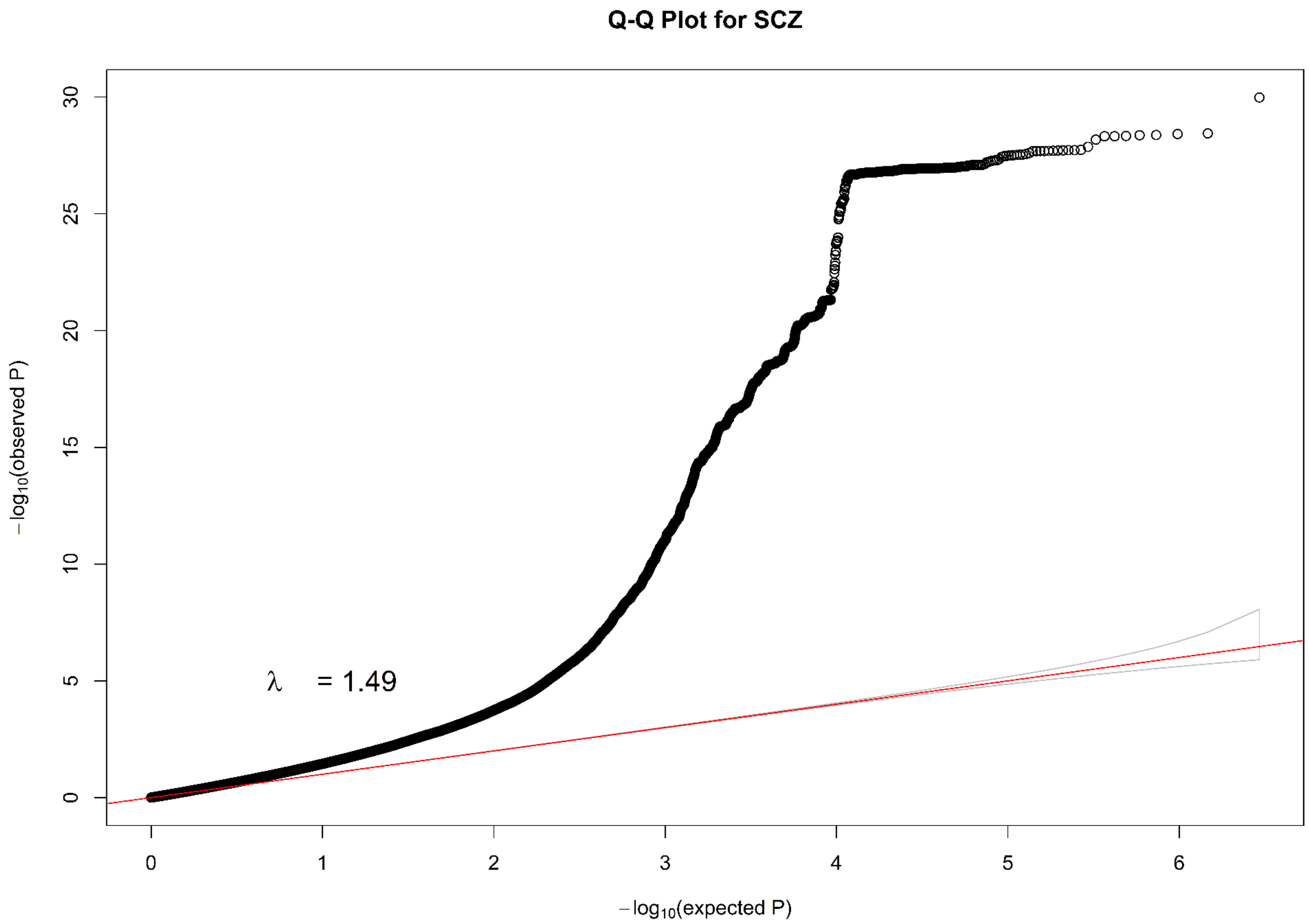


**Fig. S75 Q-Q plot for common SNPs of SCZ.**

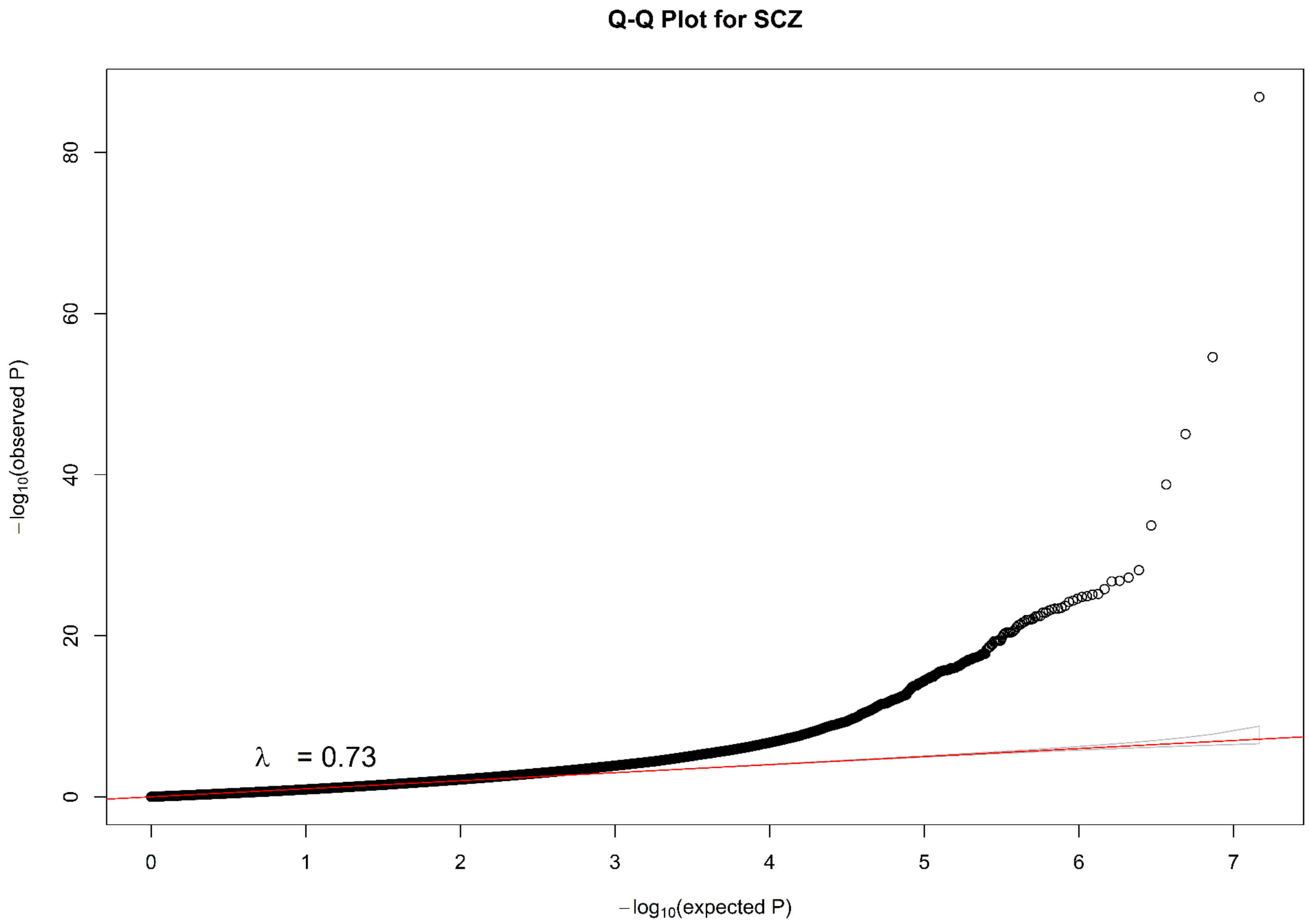


**Fig. S76 Q-Q plot for rare SNPs of SCZ.**

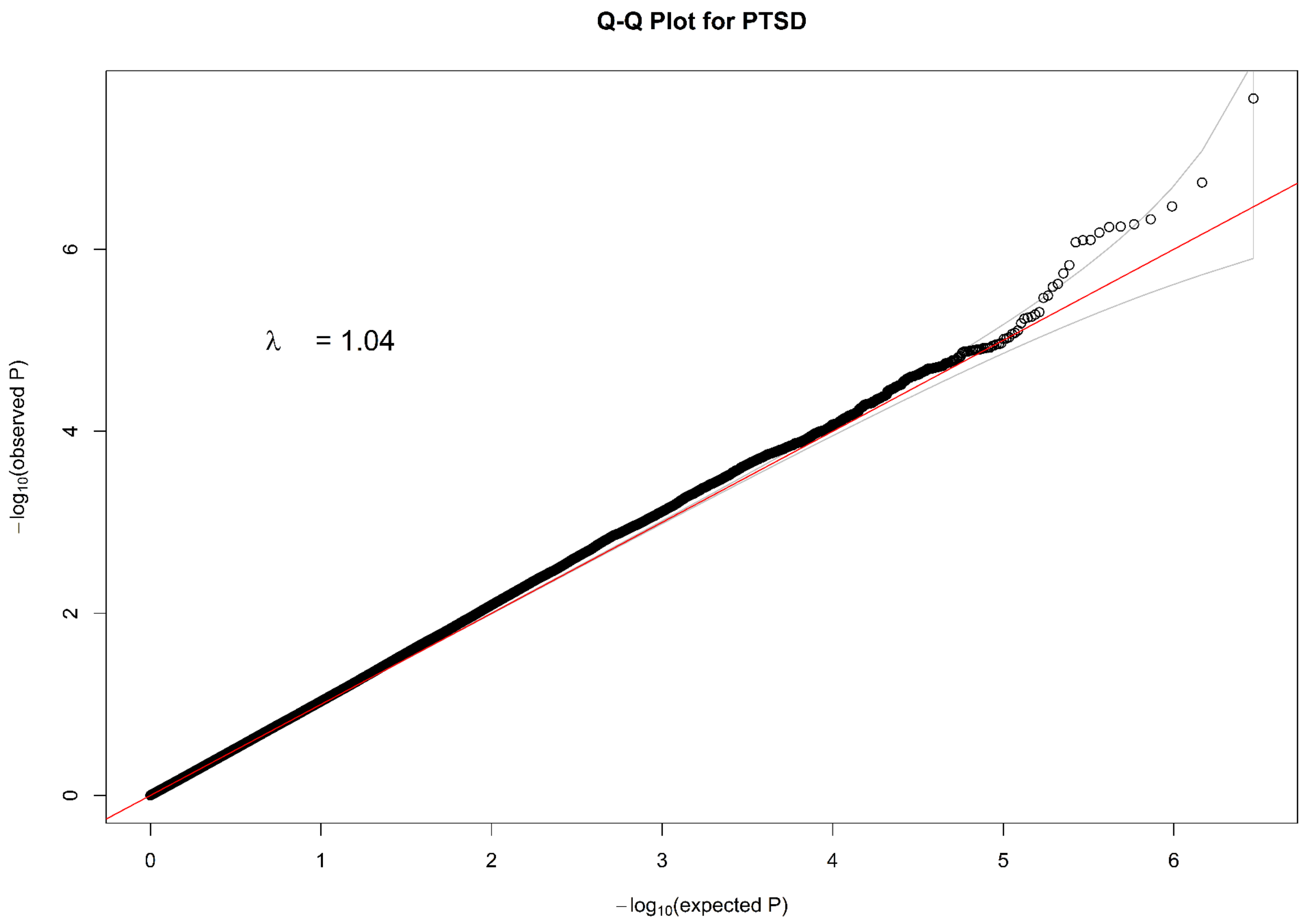


**Fig. S77 Q-Q plot for common SNPs of PTSD.**

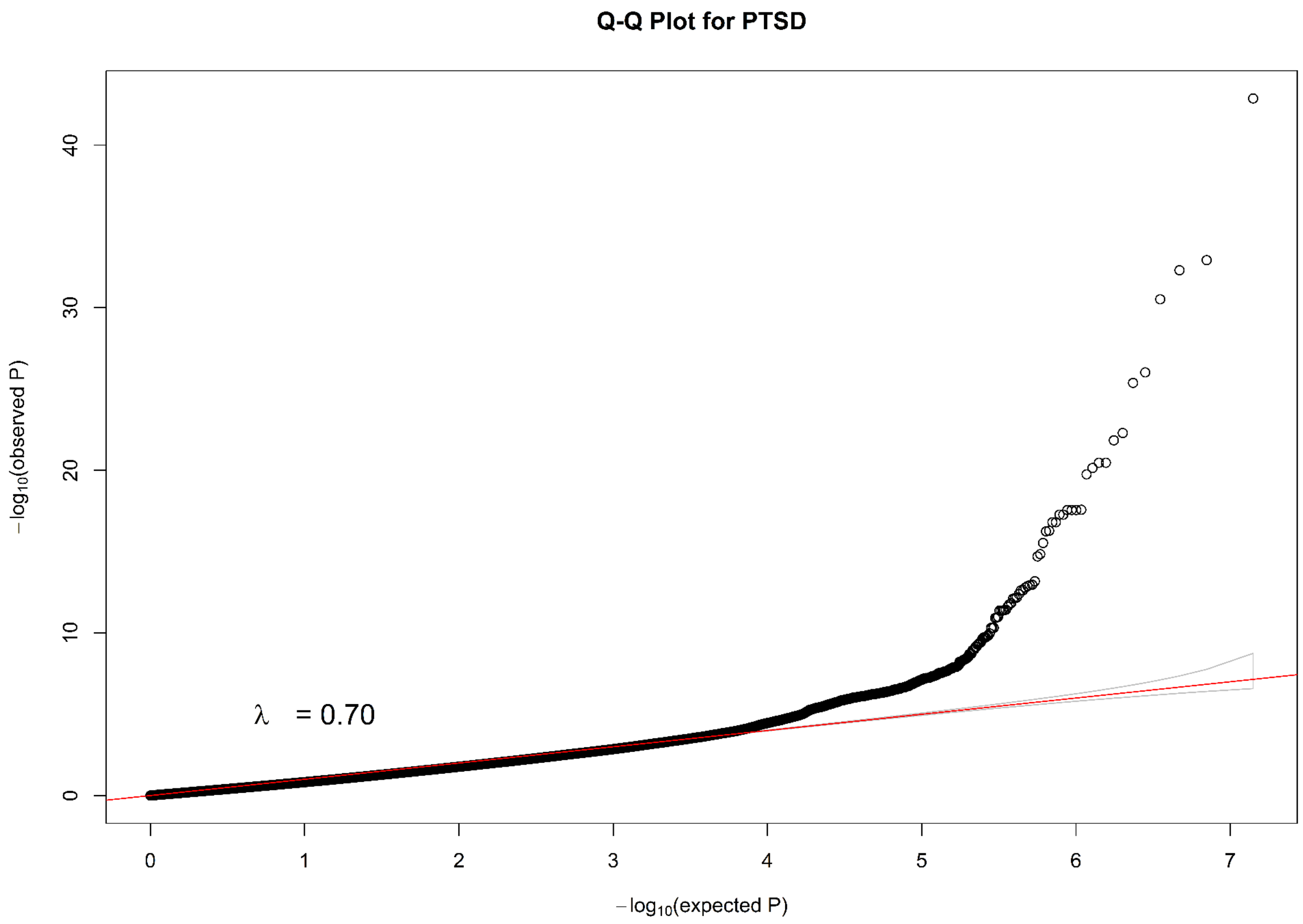


**Fig. S78 Q-Q plot for rare SNPs of PTSD.**

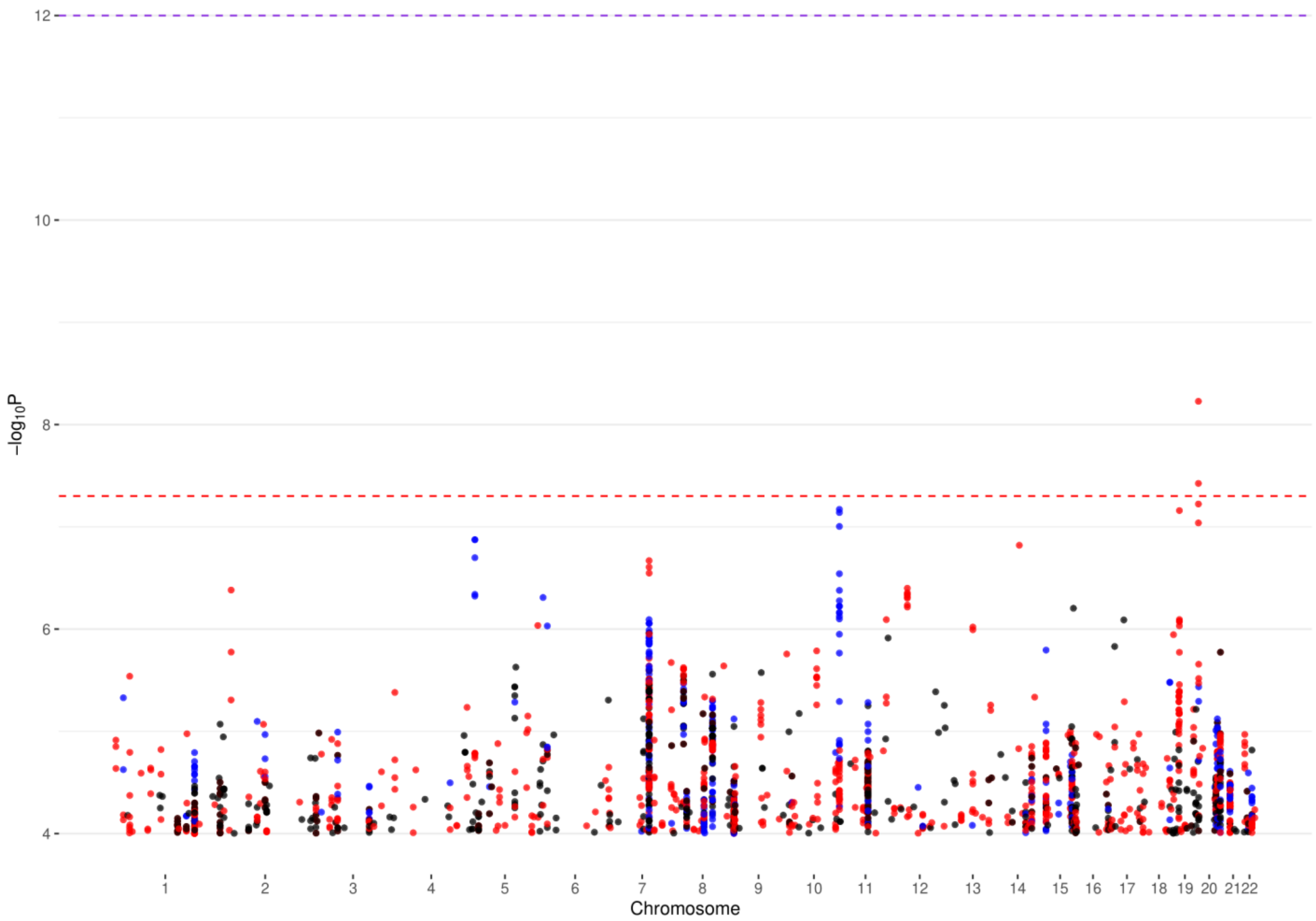


**Fig. S79 Manhattan plot of chromosomes 1-22 for ADHD. Black color denotes reported signals, red denotes DISTMIX2 signals and blue denotes ARDISS signals. The red line is the default genome-wide threshold of $p = 5 * 10^{-8}$, which is applicable common SNPs with moderate to large Info values. The purple line at $p = 10^{-12}$ is the threshold to be used for rare/very rare variants.**

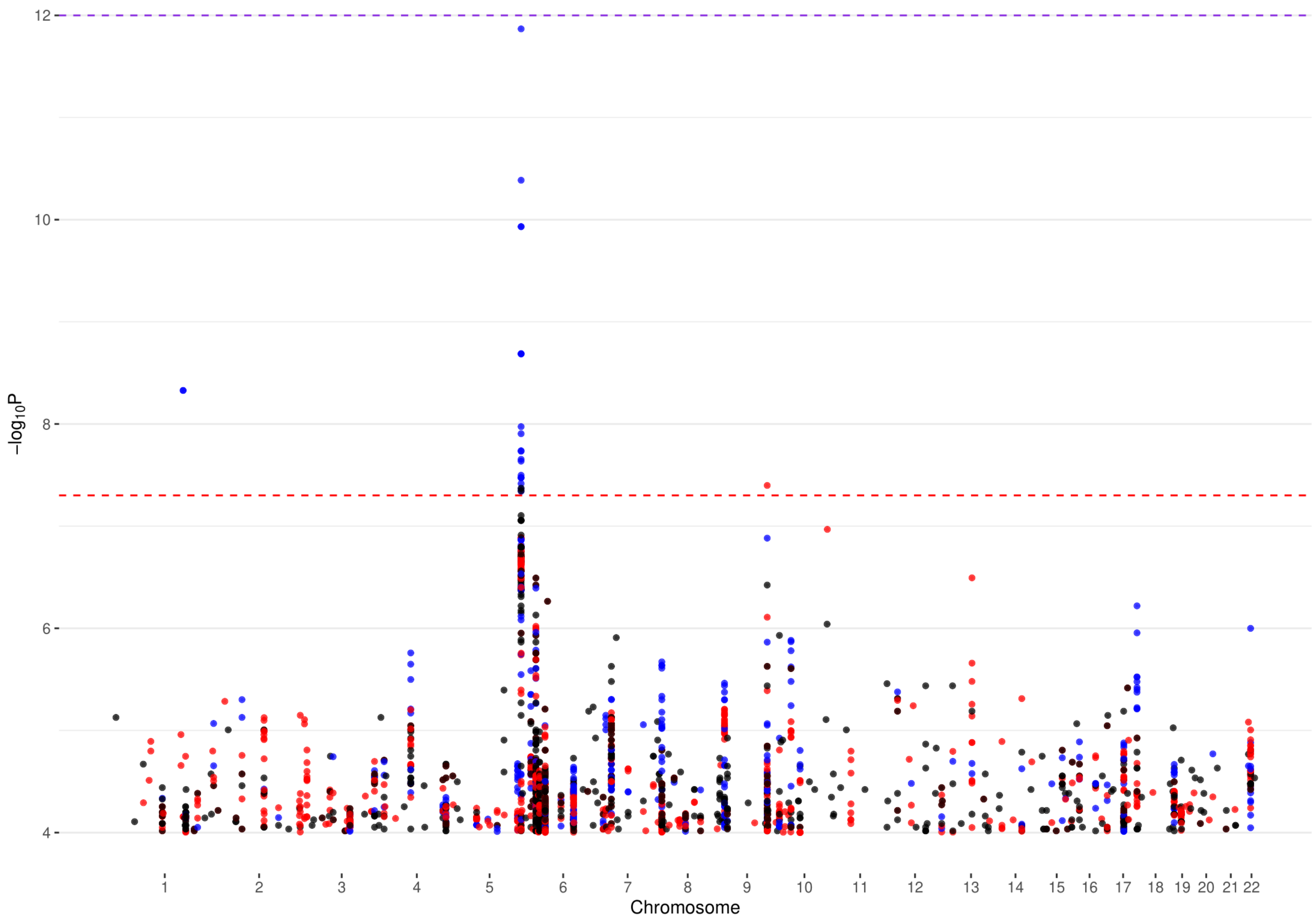


**Fig. S80 Manhattan plot of chromosomes 1-22 for AUT (see Fig. S79 for background).**

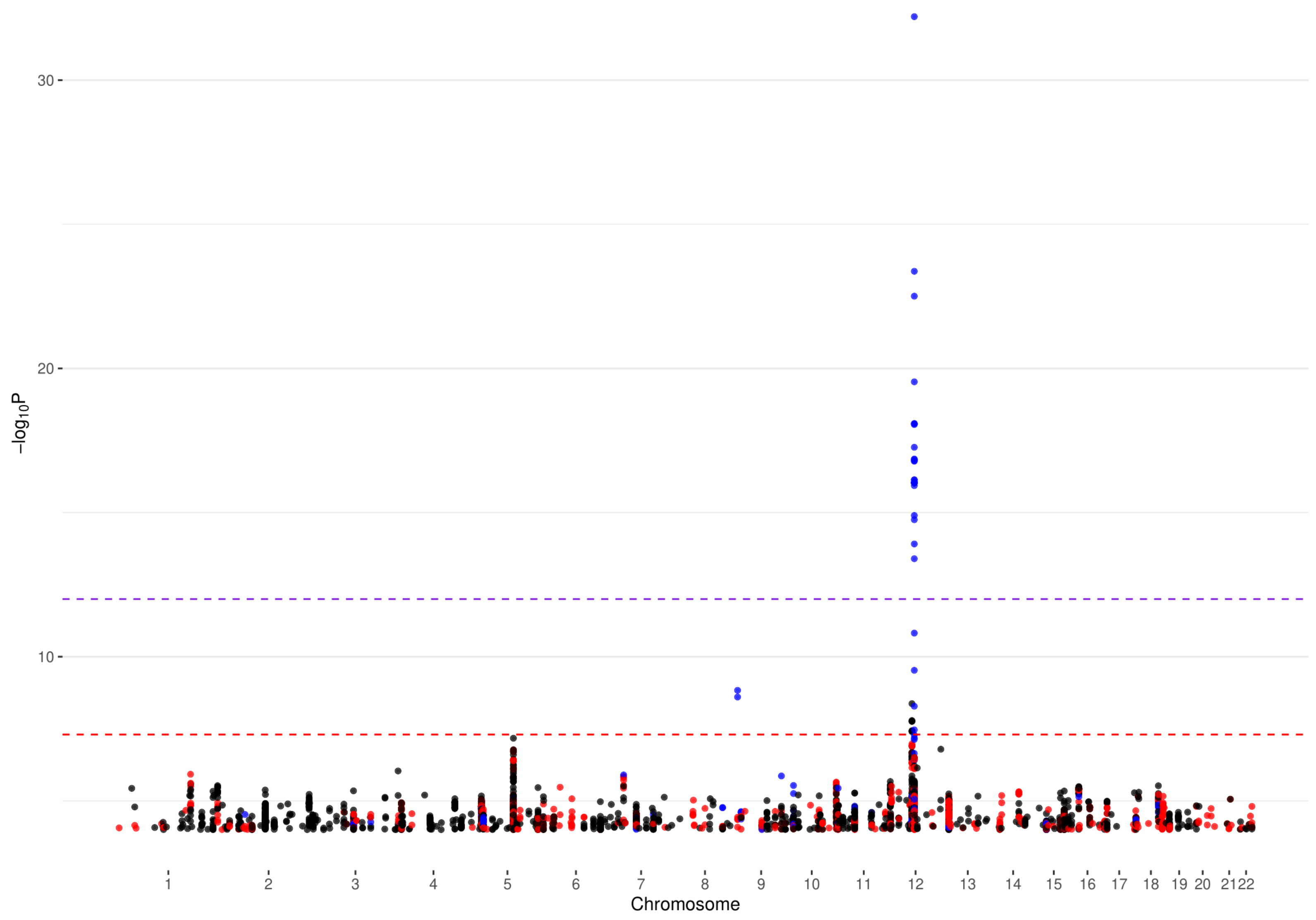


**Fig. S81 Manhattan plot of chromosomes 1-22 for BIP (see Fig. S79 for background).**

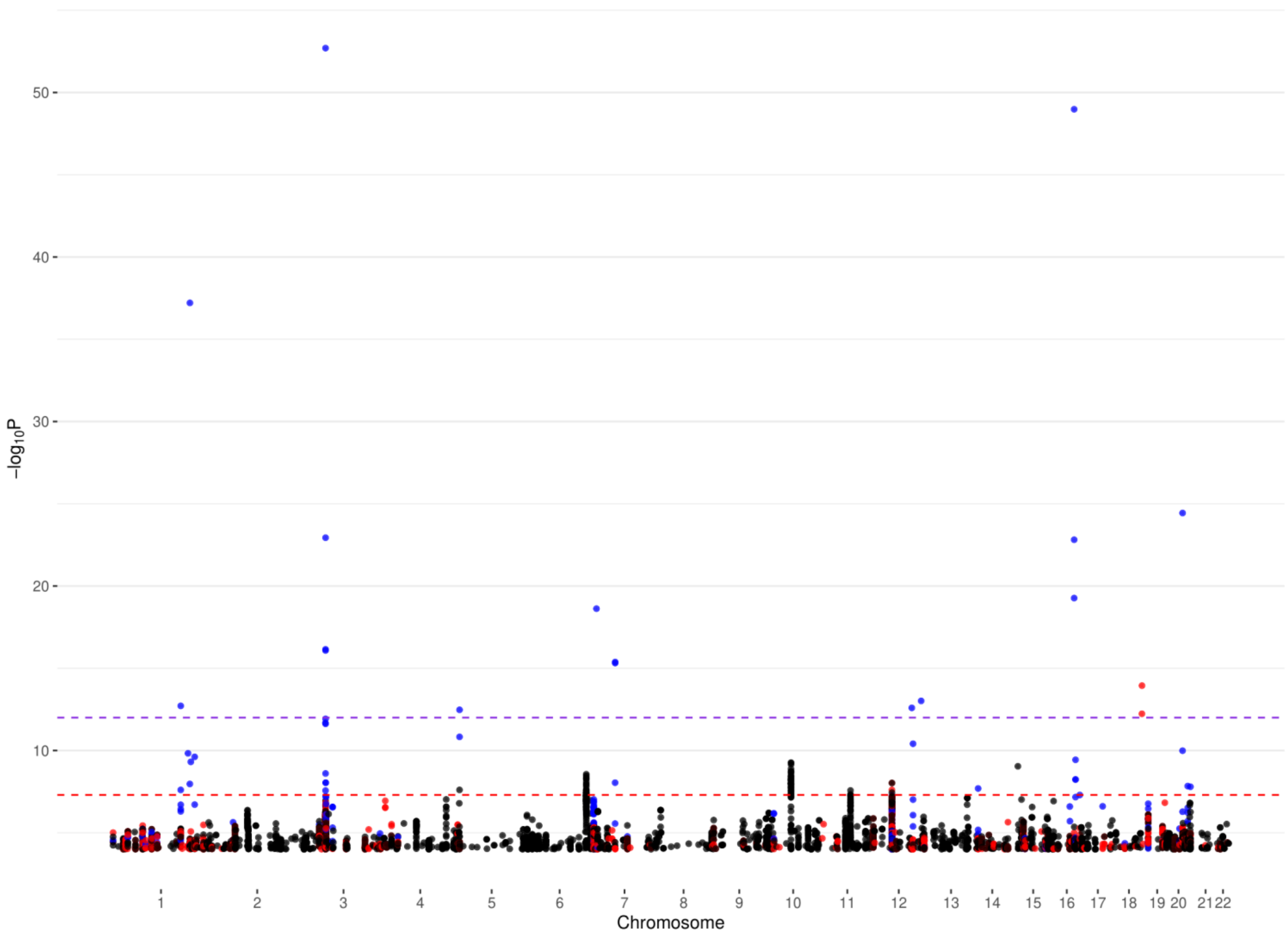


**Fig. S82 Manhattan plot of chromosomes 1-22 for ED (see Fig. S79 for background).**

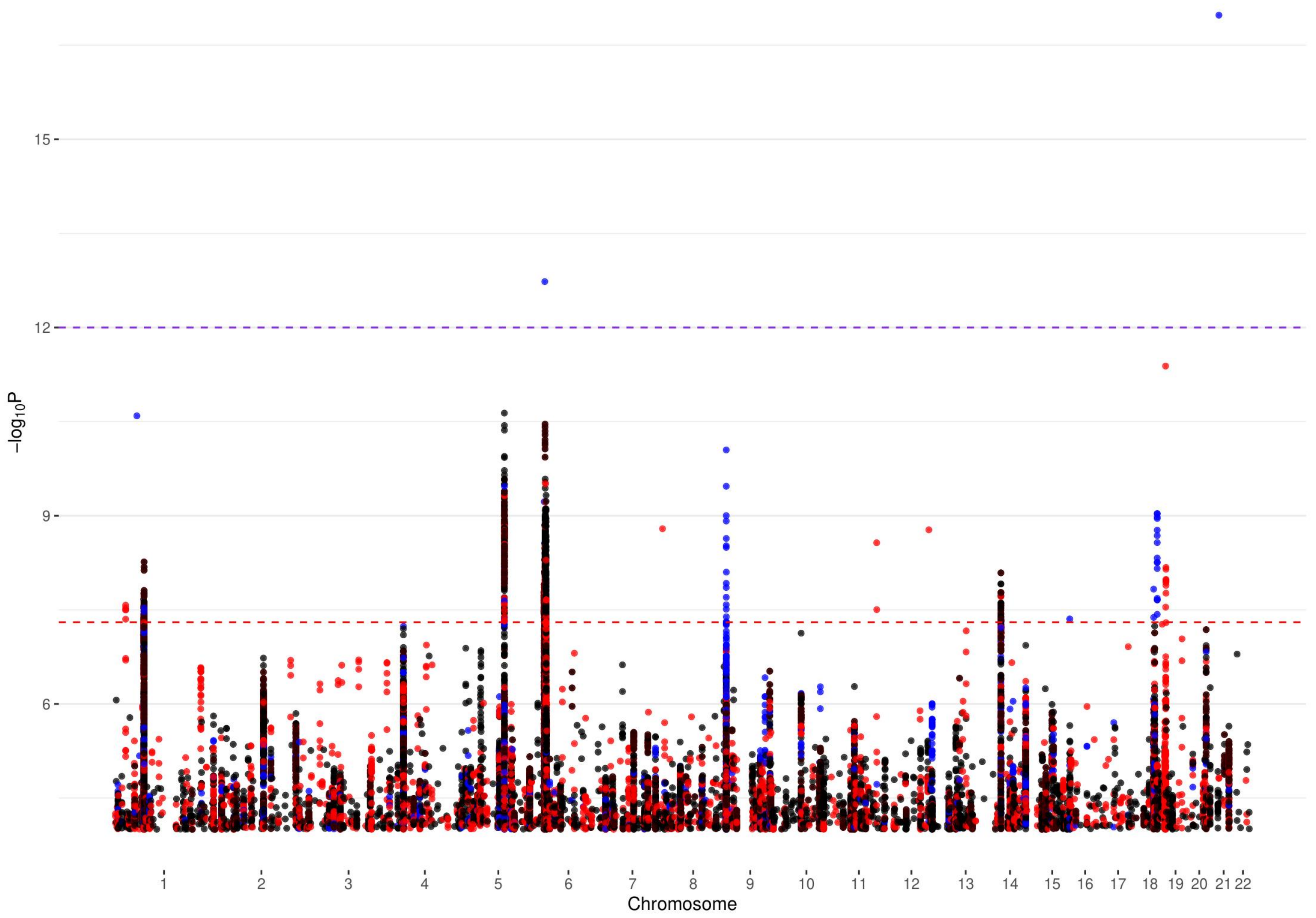


**Fig. S83 Manhattan plot of chromosomes 1-22 for MDD (see Fig. S79 for background).**

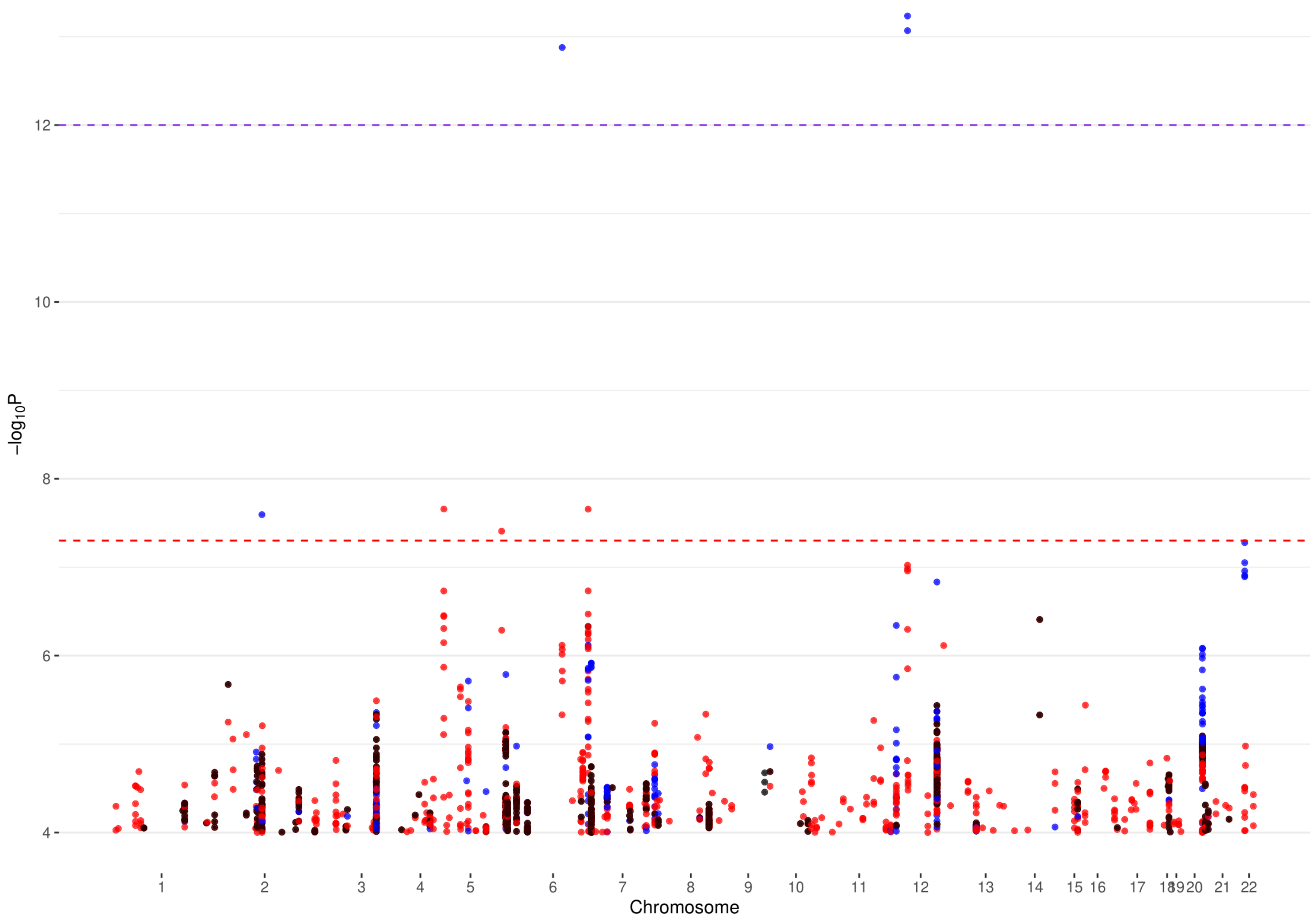


**Fig. S84 Manhattan plot of chromosomes 1-22 for PTSD (see Fig. S79 for background).**

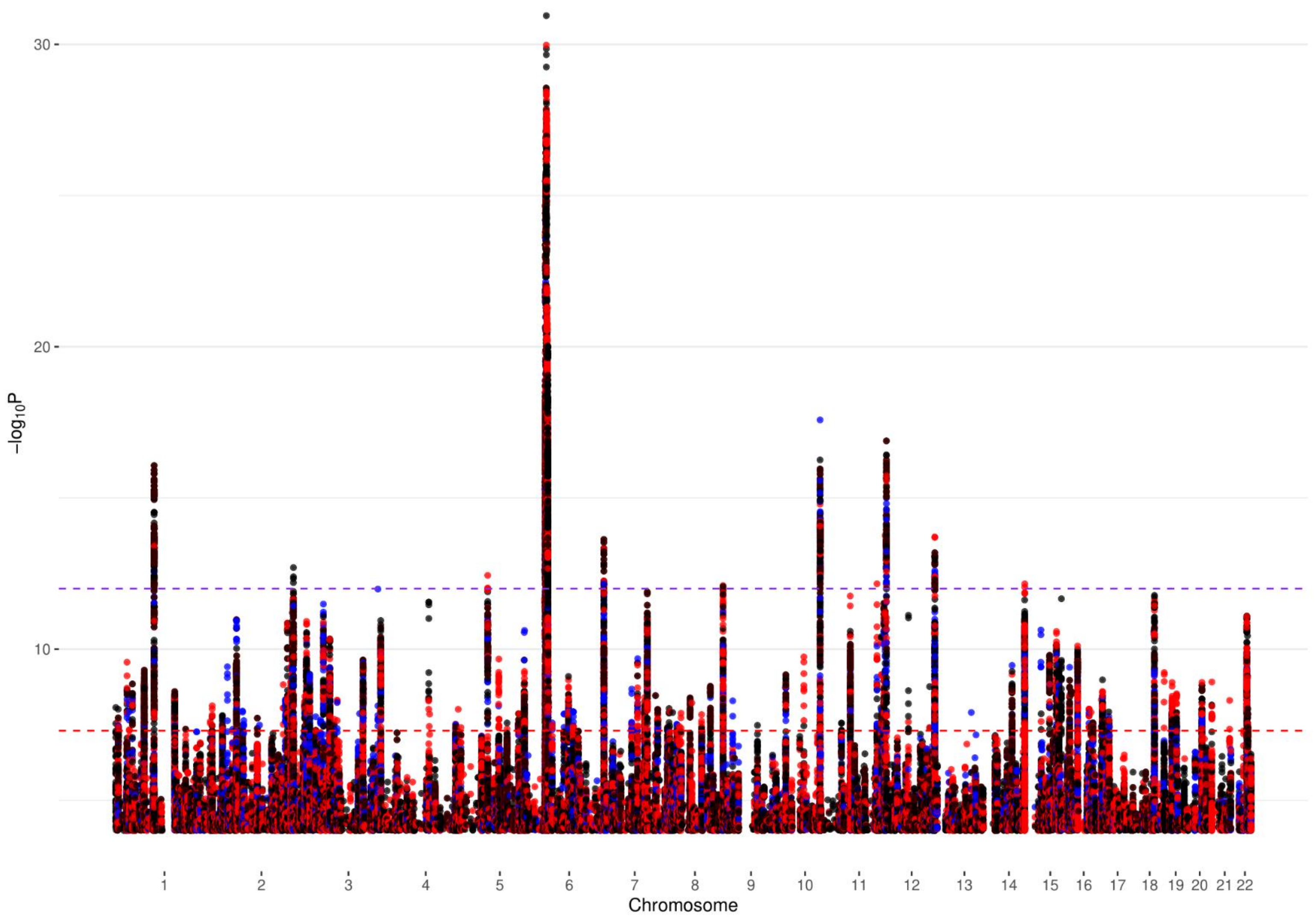


**Fig. S85 Manhattan plot of chromosomes 1-22 for SCZ (see Fig. S79 for background).**
